# Cell-type-resolved DNA methylation analysis using bulk long-read sequencing across human tissues

**DOI:** 10.64898/2026.11.18.767445

**Authors:** Hao Chen, Edward A. Mead, Yanchun Zhang, Nikolaos Karvelas, Zhangying Chen, Emma L. Thorn, Yu Fan, Mi Ni, Casidhe-Nicole Bethancourt, David A. Cohen, Gordon Fan-Huang, Zi Yan, Eric Bouhassira, Salima Soualhi, Alexey Kozlenkov, Jamshid Abdul-Ghafar, Slim Mzoughi, Ernesto Guccione, Stella Dracheva, Louis J. Cohen, Rachel Brody, Alexander W. Charney, Hao Wu, Rahul M. Kohli, Kurt W. Farrell, Jan Vijg, Joel T. Gabre, Deirdre J. Cohen, Fanny M. Elahi, John F. Crary, Gang Fang

## Abstract

DNA methylation shapes cellular identity and changes with aging and disease, but most human methylomes come from bulk tissue, which mixes signal across many cell types. A methylation change at the bulk tissue level can reflect a change in cellular composition or an intrinsic shift within a cell type, and the two cannot be separated from bulk data. Single-cell and sorted-cell methods resolve cell types directly, but their cost and workflow complexity pose challenges to large studies. Long-read sequencing of single DNA molecules can directly detect DNA methylation on each read. We built COMET (Cell-of-Origin from METhylation), a framework that uses cell-type-specific methylation to assign individual long reads to cell types from standard bulk long-read sequencing. COMET detects changes in cell-type composition robustly even from shallow bulk long-read data, quantifies methylation level of repetitive DNA within each cell type, and, with deeper sequencing, resolves methylation differences between health and disease in individual cell types. We applied COMET to 140 human blood samples, 104 human brain samples and 59 human colon samples. In blood, we found epigenetic aging is driven mainly by shifts in cell-type composition, and that APOE4 carriers have myeloid-specific methylation changes. In brain, we found sex-biased change in cell-type composition with age, resolved methylation and hydroxymethylation separately within each cell type. In colon, we found a consistent expansion of fibroblasts in tumors relative to matched normal tissue, localized methylation loss at LINE-1 in fibroblast cells, and identified tumor- and cell-type-specific hypermethylation in epithelial cells at the promoters of known colorectal cancer genes. By resolving cell-type-specific methylation directly from bulk long-read data, this approach opens broad study of cell-type epigenetics across human tissues in aging and disease.

## Introduction

DNA methylation is an epigenetic mark that shapes cellular identity and function across human tissues^1,2^, and its disruption is a molecular feature of both aging and a wide range of human diseases^1,3^. Changes in cytosine modification accompany, and in many cases help drive, the course of diseases and the gradual cellular decline of aging^4,5^. Built on large, consistently generated datasets from methylation arrays and short-read methylation sequencing, methylation-based age predictors (epigenetic clocks) and tumor classifiers have shown how reliable these associations are and how useful they can be in the clinic^6-9^.

Yet one interpretive problem has persisted. Most methylomes used to study human aging and disease come from bulk tissues, which combine signal from many cell types at once. A methylation change measured in bulk can mean that one or more cell types have changed internal methylation, or that the mix of cell types in the tissue has shifted, and these two explanations cannot be told apart from bulk data alone^10-13^.

Single-cell methylation sequencing^14-18^ and short-read sequencing of sorted cells^2,19,20^ measure cell-type methylation directly and remain the gold standard for cell-type specific DNA methylation analysis. Both, however, come at a high cost in expense and workflow complexity. As a result, methylation studies of aging and disease at the scale of large patient groups remain almost entirely based on bulk sequencing data, while cell-type-specific epigenetic biology stays confined to small, atlas-style cohorts.

These constraints motivate three goals of this study. First, we sought to determine whether cell-type composition and cell-type-specific methylation can be recovered directly from standard, unsorted bulk sequencing, without single-cell or sorted-cell input. Second, we asked whether this would allow the study of DNA methylation in human repetitive elements, which remain difficult to assay at cell-type resolution even with existing single-cell methods. Third, we asked whether resolving methylation by cell type would reveal disease- and aging-associated signals that are otherwise hidden or confounded in bulk sequencing data.

Here, we present COMET, a framework that recovers cell-type-resolved methylation from standard, scalable bulk long-read sequencing on Oxford Nanopore Technologies (ONT) and Pacific Biosciences single-molecule real-time (PacBio SMRT) platforms. These platforms read long, single molecules of DNA and detect methylation along each one^21-23^, and at sites where methylation differs between cell types, the pattern carried points to the cell it came from. COMET integrates these cell-type-specific methylation marks together to assign individual (bulk) long reads to their cell of origin, and then supports three kinds of analysis suited to how deeply a sample is sequenced. From shallow bulk data, COMET measures the cellular composition of a sample without sorting or single-cell input, and reports cell-type-resolved modification levels for the same sample. From deep bulk data, COMET identifies methylation differences between groups, such as healthy and disease, within individual cell types.

We applied COMET across three human tissues and multiple disease settings. In blood, we found age-related methylation change was explained largely by shifts in cell-type composition, and APOE4 carriers showed methylation differences specific to myeloid cells. In brain, cell-type proportions changed with age differently in males and females, and measuring methylation and hydroxymethylation separately within each cell type uncovered regions, specific to cognitive groups, where the two marks shift in opposite directions. In colon, tumors showed a consistent expansion of fibroblasts relative to matched normal tissue and LINE-1 (L1) methylation loss in fibroblasts, along with hypermethylation specific to both tumor and epithelial cells at the promoters of known colorectal cancer (CRC) genes. Collectively, these results establish a general, cost-effective strategy for studying cell-type epigenetics directly from bulk long-read sequencing, at a scope not previously attainable in these three tissues, and support a broad range of future studies of cell-type-resolved epigenetics across human tissues and diseases.

## Results

### COMET assigns individual long reads to their cell type of origin with high accuracy across three human tissues

We assembled samples from three human tissues (blood, colon, and brain) and generated native long-read sequencing data on the PacBio Revio or ONT P2 platforms (**Fig. 1A** and **Methods**). 297 samples were sequenced using the PacBio platform: average depth for shallow sequencing was 1.98x (range 0.30 to 15.64×) for 140 blood samples, 3.39x (0.41 to 10.98×) for 53 colon samples, and 3.09x (0.32 to 13.19×) for 104 brain samples (**Fig. 1A, Supplementary Fig. 1A**, and **Supplementary Table 1**). Median length ranged from 8,315 to 14,262 bp (**Supplementary Fig. 1B** and **Supplementary Table 1**). We also sequenced six colon samples using ONT P2 from our previous study^24^ (**Supplementary Fig. 1C** and **Supplementary Table 1**). Across the three tissues, 18 samples were additionally sequenced to high depth (21.16 to 60.15×) to support cell-type specific methylation analyses: 8 blood samples, 6 brain samples and 4 colon samples **(Supplementary Fig. 1D** and **Supplementary Table 1**). DNA modification was called from raw sequencing reads for PacBio and ONT data, followed by additional quality control steps (**Supplementary Fig. 2A–D** and **Methods**).

**Fig. 1.**
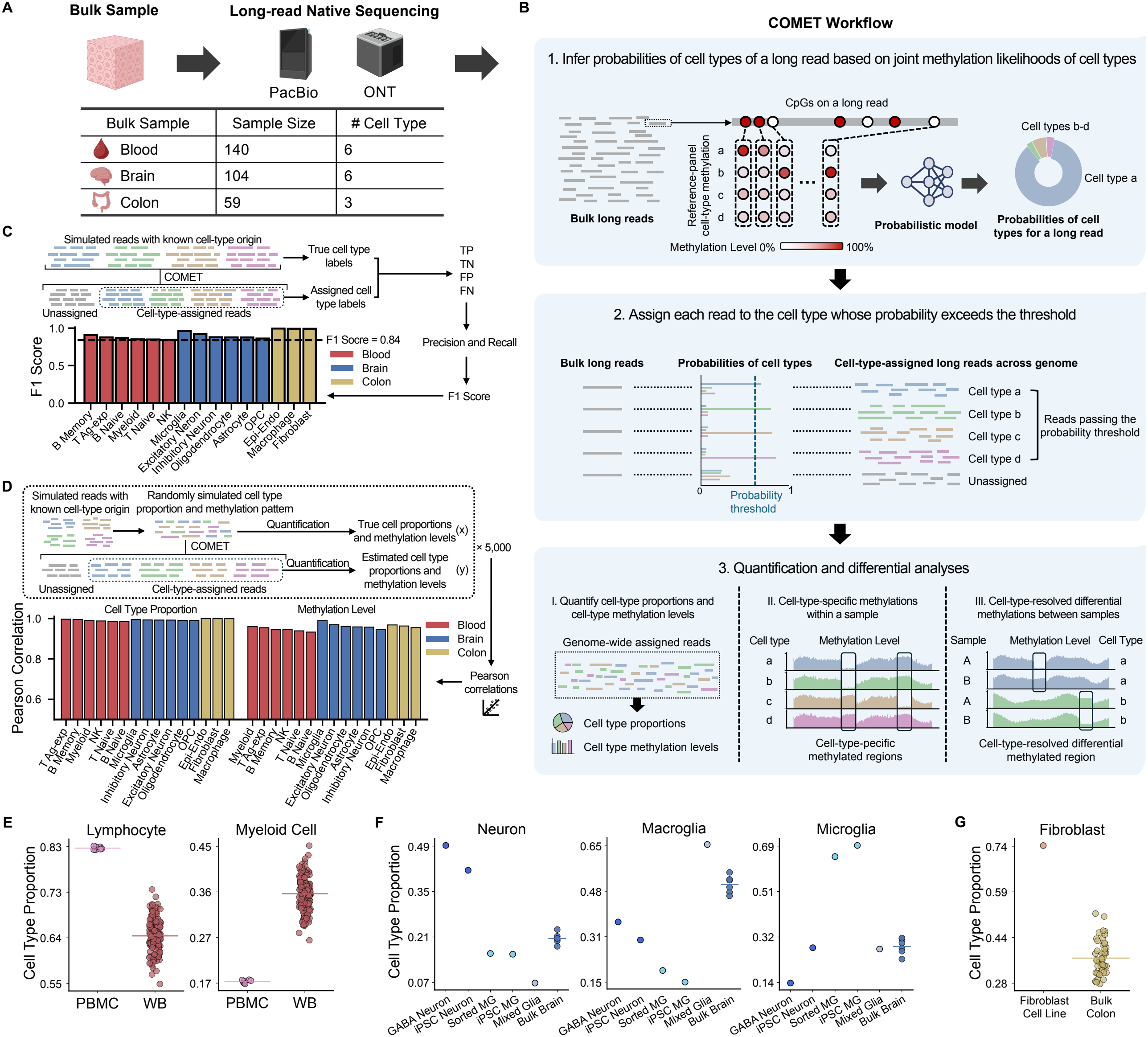
COMET workflow and evaluation. **(A)** Study design and cohort information. Bulk tissue samples from three human tissues were sequenced natively on PacBio Revio and ONT P2 platforms and analyzed with COMET. The table summarizes the sample size and the number of cell types resolved for each tissue, covering 140 blood, 104 brain and 59 colon samples. **(B)** Schematic of the COMET workflow. (1) For each long read in a bulk long-read dataset, cell-type probabilities are computed by a probabilistic model from the joint methylation likelihoods across all CpGs the read covers in the tissue-matched reference panel. In the example shown, color indicates methylation level of CpGs in the reference panel. (2) Across the bulk reads, those whose highest cell-type probability exceeds the threshold are assigned to that cell type, and the remainder are left unassigned. (3) Cell-type-assigned reads support three downstream analyses: (I) quantification of cell-type proportions and cell-type-resolved methylation levels, (II) identification of cell-type-differential or cell-type-specific methylation within a sample, and (III) identification of cell-type-resolved differential methylation between samples. **(C)** Simulation-based evaluation of cell type assignment. Long reads were simulated from each cell type’s reference methylome and carry that cell type as their true label. COMET was applied to each simulated read set, and the assigned labels were compared with the true labels to obtain true and false positives and negatives, from which precision, recall and the F1 score were calculated. Bars show the F1 score for every cell type across the three tissues at the selected probability thresholds, 0.65 for blood and brain and 0.50 for colon, with the dashed line marking F1 = 0.84. **(D)** Simulation-based evaluation of cell-type quantification. Simulated reads of known cell-type origin were combined into mixtures of defined composition and methylation level, with 5,000 mixtures generated per tissue at a minimum cell-type proportion of 5%. COMET was applied to each mixture, and cell-type proportions and cell-type-resolved methylation levels estimated from the assigned reads were compared with the known values of the mixture. Bars show the Pearson correlation between estimated and true values across the 5,000 mixtures, for each cell type and for both quantities. **(E)** COMET-estimated proportions of lymphocytes (B cells, T cells, and NK cells) and myeloid cells in PacBio-sequenced PBMC samples (n = 7) and whole-blood samples (n = 140), using the blood reference panel. **(F)** COMET-estimated proportions of neurons (excitatory, inhibitory, and ambiguous neurons), macroglia (astrocytes, oligodendrocytes, OPCs, and ambiguous macroglia) and microglia across ONT-sequenced sorted or derived brain cell populations and bulk brain samples, using the brain reference panel. These comprised GABAergic neurons (n = 1), iPSC-derived neurons (n = 1), sorted microglia (n = 1), iPSC-derived microglia (n = 1), mixed glia (n = 1) and bulk brain samples (n = 6). **(F)** COMET-estimated fibroblast proportion in PacBio-sequenced colon fibroblast cell line (n = 1) and bulk colon samples (n = 53), using the colon reference panel. In **(E–G)**, each point represents one sample and horizontal lines indicate the group mean.

**Fig. 2.**
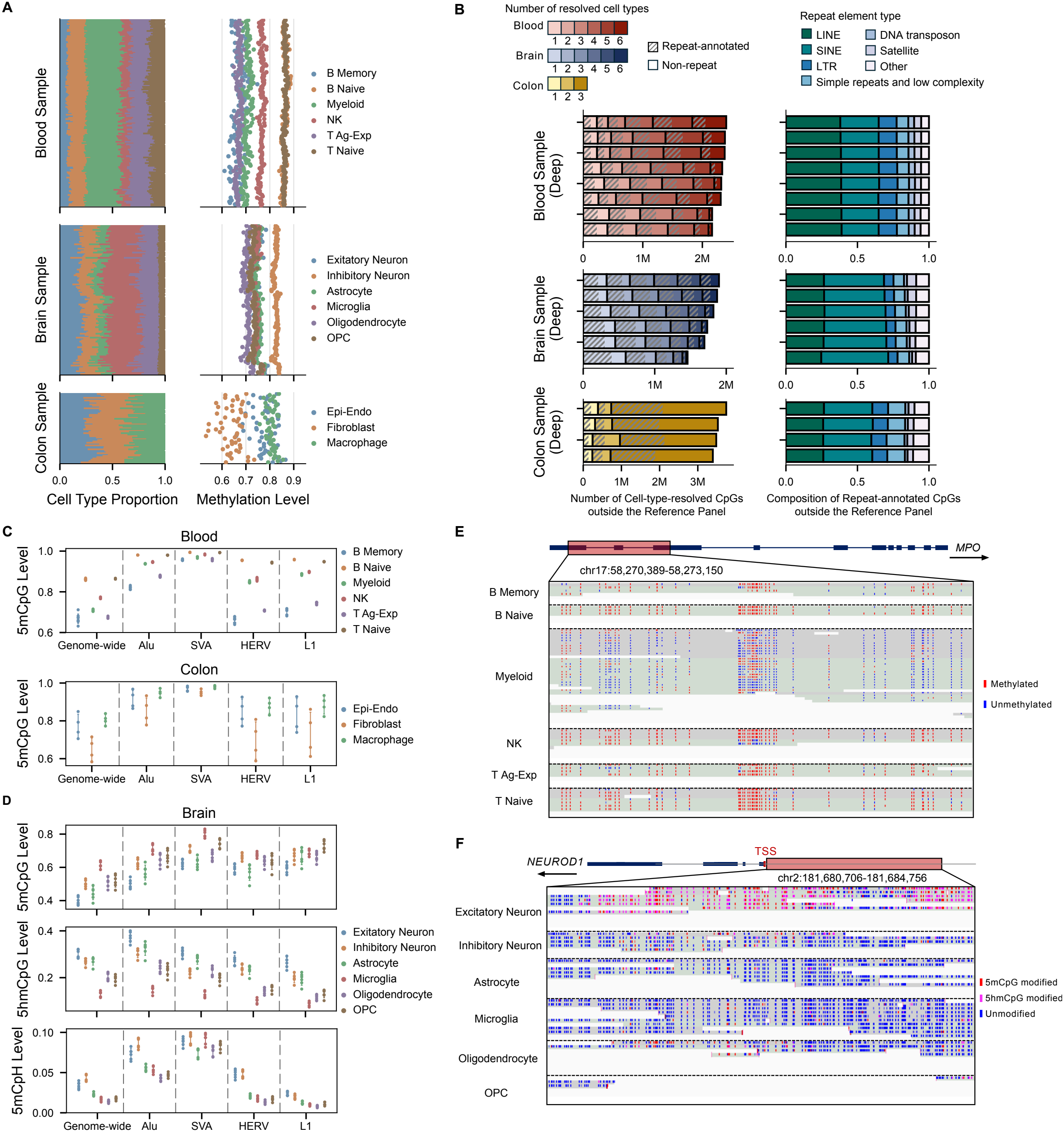
Cell-type-resolved genome-wide and repeat-element methylation across tissues. **(A)** Cell-type proportions (left) and cell-type-resolved genome-wide methylation levels (right) for the shallow PacBio-sequenced samples, shown for blood (n = 140), brain (n = 104), and colon (n = 53) from top to bottom. Each horizontal bar in the left panels represents one sample, and each row of points in the right panels gives the cell-type methylation levels for that same sample, with samples aligned across the two panels. **(B)** CpGs resolved to a cell type that fall outside the reference panel, quantified from the deep-sequenced samples. Left panels give the number of such CpGs, with blood (n = 8), brain (n = 6), and colon (n = 4) from top to bottom, and each bar representing one sample. Shading from light to dark indicates the number of cell types covering those CpGs, and hatching marks the subset overlapping an annotated repeat. Right panels give the composition of repeat classes among the repeat-annotated CpGs. Blood and colon were deep sequenced on PacBio, brain on ONT. **(C)** Cell-type-resolved methylation levels (5mCpG) genome-wide and at transposable element (TE) classes in the deep PacBio-sequenced samples, for blood (n = 8) and colon (n = 4) from top to bottom. Each point represents one sample and error bars give the standard deviation across samples. **(D)** Cell-type-resolved modification levels for 5mCpG, 5hmCpG and 5mCpH, genome-wide and at TE classes, in the deep ONT-sequenced brain samples (n = 6). Each point represents one sample and error bars give the standard deviation across samples. **(E)** Genome browser view of the *MPO* locus in a representative deep-sequenced blood sample, showing reads assigned to each of the six blood cell types. The gene model is shown above, with the displayed region (chr17:58,270,389–58,273,150) highlighted. Myeloid-assigned reads carry extensive unmethylated CpGs across the region, whereas reads assigned to the five lymphoid classes are predominantly methylated. The red and blue marks indicate methylated and unmethylated CpGs, respectively. **(F)** Genome browser view of a transcription start site (TSS) upstream of *NEUROD1* in a representative deep ONT-sequenced brain sample, showing reads assigned to each of the six brain cell types. The gene model is shown above with the displayed region (chr2:181,680,706– 181,684,756) highlighted. Excitatory neuron-assigned reads carry 5hmCpG modification across the region, whereas reads from the other five cell types are predominantly unmodified. The red, magenta and blue marks indicate 5mCpG-modified, 5hmCpG-modified and unmodified CpGs, respectively. In **(E–F)**, each row represents one read, grouped by assigned cell type and separated by dashed lines. The arrow under the gene name indicates the orientation of the gene.

For each tissue, we curated a cell-type methylation reference panel from data independent from our long-read cohorts, using sorted-cell bisulfite sequencing^2,19^ for blood and colon and single-cell methylation sequencing^25^ for brain (**Supplementary Fig. 3A** and **Methods**). To decide how many cell types to include in each panel, we performed principal component analysis of the genome-wide CpG methylation profiles of the candidate cell types, keeping cell types that separate into clear, distinct clusters and not distinguishing subtypes that the methylation data cannot yet reliably resolve, such as the T cell subtypes within the antigen-experienced T cell cluster (**Supplementary Fig. 3B–D** and **Methods**). This analysis led us to six cell types in blood: memory B cells (B Memory), naive B cells (B Naive), myeloid cells, natural killer cells (NK), antigen-experienced T cells (T Ag-exp, encompassing effector and memory T cells), and naive T cells (T Naive); six cell types in brain: excitatory neurons, inhibitory neurons, astrocytes, microglia, oligodendrocytes, and oligodendrocyte progenitor cells (OPC); and three cell types in colon: combined endocrine and epithelial cells (Epi-Endo), fibroblasts, and macrophages.

**Fig. 3.**
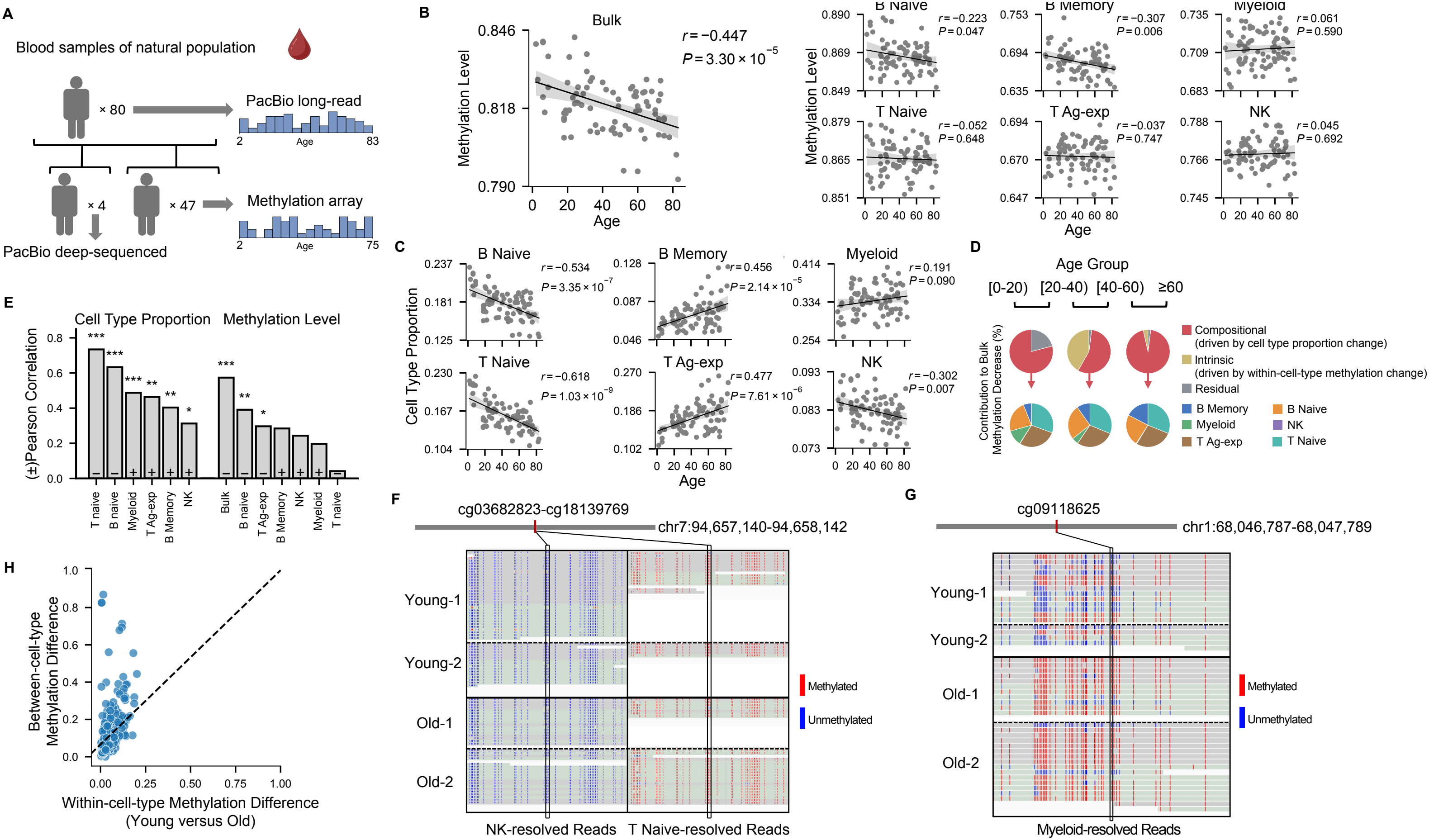
Cell-type composition and cell-type-resolved methylation changes in blood aging. **(A)** Cohort and sequencing design for the blood aging analysis. 80 blood samples spanning the indicated age range were analyzed by PacBio long-read sequencing. Four samples were further deep sequenced by PacBio, and methylation array data were generated for 47 samples. **(B)** COMET-estimated genome-wide bulk and cell-type-resolved methylation levels across age. Each plot shows the bulk or indicated cell-type methylation level (y-axis) as a function of subject age (x-axis). Solid lines indicate linear regression fits, with shaded areas representing 95% confidence intervals (CIs). Pearson’s *r* and *P* values, adjusted for sex, are shown in each plot. **(B)** COMET-estimated cell-type proportions across age. Each plot shows the proportion of the indicated cell type (y-axis) as a function of subject age (x-axis). Solid lines indicate linear regression fits, with shaded areas representing 95% CIs. Pearson’s *r* and *P* values, adjusted for sex, are shown in each plot. **(C)** Compositional and intrinsic contributions to bulk methylation decreases across age groups. Blood samples were divided into four age groups ([0,20), [20,40), [40,60), and ≥60) and each pair of adjacent groups was compared. Upper pie charts show the relative contributions of compositional changes driven by changes in cell-type proportions, intrinsic changes driven by within-cell-type methylation changes, and a residual term comprising unassigned reads and the covariance between proportion and methylation changes, to the bulk methylation decrease. Lower pie charts show the contributions of individual cell types to the compositional component. **(D)** Correlations between methylation array-derived Horvath clock age and COMET-estimated cell-type measurements in 47 samples with matched methylation array data. Each bar represents the absolute Pearson correlation coefficient (Pearson’s *r*) between Horvath clock age and a cell-type proportion (left) or bulk and cell-type-resolved methylation level (right). “+” and “−” signs below the bars indicate positive and negative correlations, respectively. Asterisks indicate sex-adjusted Pearson correlation *P* values: \**P*_adj_ < 0.05, \*\**P*_adj_ < 0.01, and \*\*\**P*_adj_ < 0.001. **(E)** Example of two adjacent Horvath clock CpGs showing distinct cell-type-specific methylation patterns. Read-level methylation within ±500 bp of the two CpGs is visualized in IGV for NK- and T naive-resolved reads across four deep-sequenced samples, including two young and two old subjects. Red and blue indicate methylated and unmethylated CpGs, respectively. **(F)** Example of a Horvath clock CpG showing an age-associated methylation difference within myeloid cells. Read-level methylation within ±500 bp of the CpG is visualized in IGV for myeloid-resolved reads across four deep-sequenced samples, including two young and two old subjects. Red and blue indicate methylated and unmethylated CpGs, respectively. **(G)** Between-cell-type and within-cell-type and methylation differences at Horvath clock CpGs based on the deep-sequenced young vs. old samples. Each dot represents a ±500-bp region centered on a Horvath clock CpG (n = 346). The y-axis was defined as the mean absolute methylation difference among cell types within samples, as illustrated by the representative example in **(F)**, whereas the x-axis was defined as the mean absolute methylation difference within cell types between young and old samples, as illustrated by the representative example in **(G)**.

The design of COMET leverages a property of long-read sequencing that an individual long read spans many more CpGs than short read sequencing. COMET integrates every CpG site a read covers, treating each as a partially informative vote, and combines the cell-type methylation likelihoods across all of them, drawn from the reference panel of the matching tissue, into a single probability for each candidate cell type (**Fig. 1B** and **Methods**). Reads whose combined probability for the top candidate cell type exceeds a defined threshold are called as confidently assigned; the remaining, more ambiguous reads are set aside rather than forced into a call (**Fig. 1B** and **Methods**). These read-level assignments anchor several downstream analyses. The fraction of reads assigned to each cell type yields an estimate of cellular composition, while reads confidently assigned to a specific cell type enables identification of cell-type-specific methylation within a sample, as well as cell-type-resolved differential methylation analysis between samples or between groups (**Fig. 1B** and **Methods**).

To benchmark assignment performance, we first simulated long reads of known cell-type origin from the reference panels and asked how accurately COMET recovered the identity of each read (**Fig. 1C** and **Methods**). Testing probability thresholds from 0.5 to 0.95, we observed the expected trade-off, with the F1 score, the harmonic mean of precision and recall, rising as the threshold increased (**Supplementary Fig. 4A**) while the proportion of assigned reads fell (**Supplementary Fig. 4B**). We therefore set the operating threshold for each tissue so that every cell type reached an F1 score above 0.84 (range 0.845 to 0.961; **Fig. 1C** and **Supplementary Fig. 4A**), and more than 40% of simulated reads from each cell type received an assignment (**Supplementary Fig. 4B**). Applying these thresholds to our cohort, the average proportion of assigned reads is 34.15% (27.87%–41.64%) for blood samples, 48.78% (44.48%– 59.67%) for brain samples, and 78.06% (66.93%–83.46%) for colon samples (**Supplementary Fig. 4C)**. For each tissue, we simulated 5,000 read mixtures with randomly drawn cell-type proportions (**Fig. 1D, Supplementary Fig. 5A**, and **Methods**), and estimated proportions tracked the true input composition closely for every cell type (**Fig. 1D** and **Supplementary Fig. 5B**), and estimated cell-type-resolved methylation levels agreed equally well (**Fig. 1D** and **Supplementary Fig. 5C**).

**Fig. 4.**
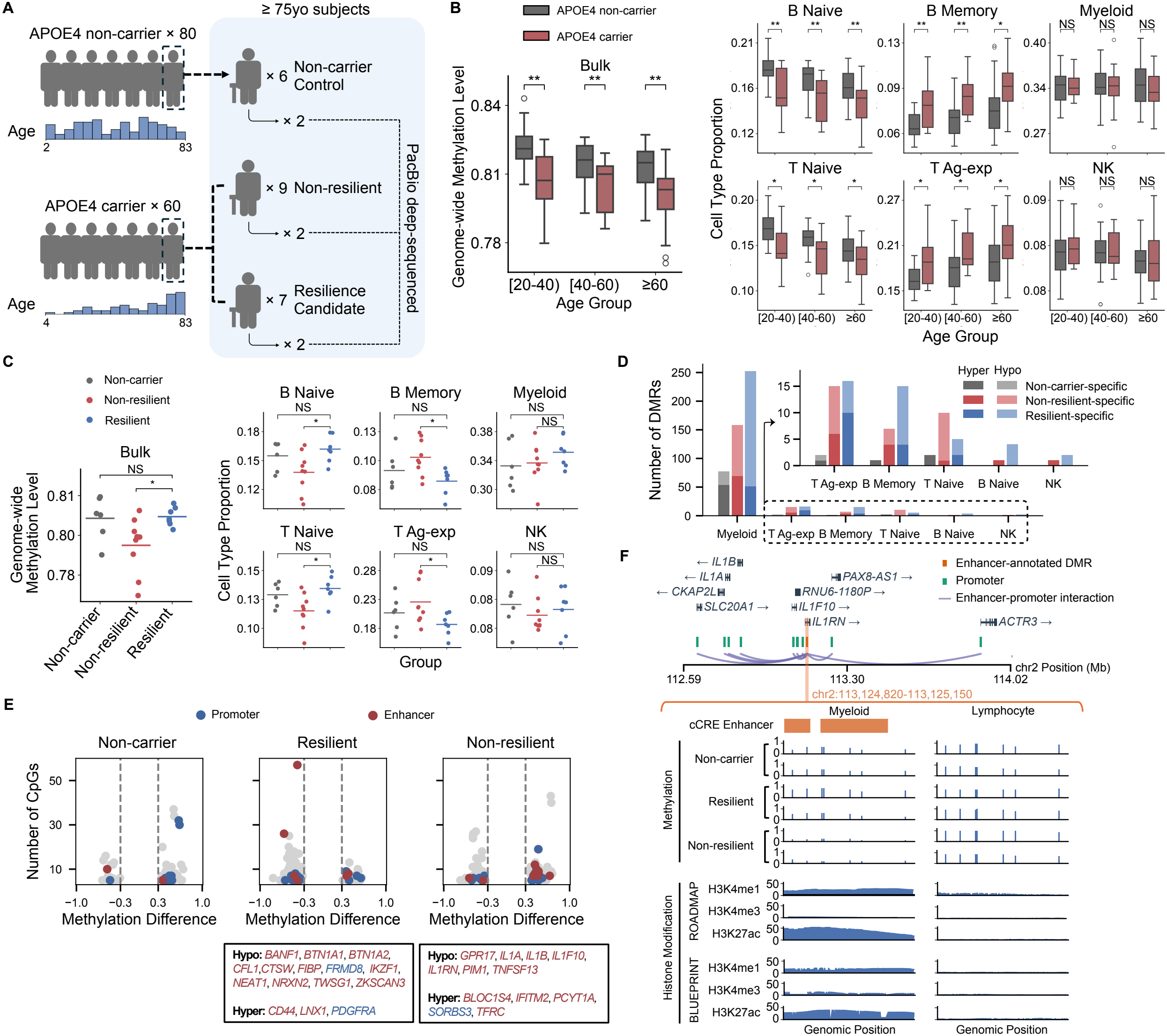
Blood cell-type composition shifts and myeloid-specific methylation changes in APOE4 carriage and AD resilience. **(A)** Cohort design and sequencing information for 140 blood samples. The cohort included 80 APOE4 non-carriers and 60 APOE4 carriers across the indicated age range. Among subjects aged ≥75 years, 6 APOE4 non-carrier controls, 9 non-resilient subjects, and 7 resilient subjects were selected for resilience analysis. Two samples from each group were further deep sequenced by PacBio. **(B)** COMET-estimated genome-wide bulk methylation levels and cell-type proportions in APOE4 carriers and non-carriers across age groups. Boxplots show genome-wide methylation levels (left) and cell-type proportions (right) for the indicated age groups. Grey and red indicate APOE4 non-carriers and carriers, respectively. Seven resilience candidates were removed from the ≥60 carrier group for this comparison, since they do not represent the typical carrier phenotype. Asterisks indicate Benjamini–Hochberg-adjusted *P* values from mixed-effects models adjusted for age and sex: \**P*_adj_ < 0.05 and \*\**P*_adj_ < 0.01. NS, not significant. **(C)** COMET-estimated genome-wide bulk methylation levels and cell-type proportions in non-carrier, non-resilient, and resilient groups. Dot plots show genome-wide methylation levels (left) and cell-type proportions (right) for non-carrier (grey), non-resilient (red), and resilient (blue) subjects. Horizontal lines indicate group means. Asterisks indicate Benjamini–Hochberg-adjusted *P* values from mixed-effects models adjusted for age and sex: \**P*_adj_ < 0.05. NS, not significant. **(D)** Group-specific, cell-type-resolved DMRs identified from PacBio deep-sequencing data for six samples, two from each group. DMRs were defined by a difference in methylation level of at least 0.3 between the two samples of one group and the remaining four. Bars show the number of group-specific DMRs across the six blood cell types in the non-carrier (grey), non-resilient (red) and resilient (blue) groups. Dark and light shades indicate hypermethylated and hypomethylated DMRs respectively. The inset shows the lymphocyte populations at an enlarged scale, as these contained far fewer group-specific DMRs than myeloid cells. **(E)** Group-specific, myeloid-resolved DMRs mapped to gene-linked regulatory elements. Each point represents a DMR, with the methylation difference between the indicated group and the other two groups on the x-axis and the number of CpGs in the DMR on the y-axis. Panels show the non-carrier, resilient, and non-resilient groups from left to right. Within each panel, points to the left of zero are hypomethylated and those to the right are hypermethylated in the indicated group, and vertical dashed lines mark the methylation-difference cutoffs used for DMR identification. Blue and red points indicate DMRs mapped to active myeloid promoters and enhancers respectively. Boxes below the plots list genes linked to these promoters or enhancers that have been associated with AD, APOE4 or inflammaging, separated by the direction of the methylation difference, with gene names colored blue for promoters and red for enhancers. **(F)** Example of a group-specific, myeloid-resolved DMR mapped to an active enhancer linked to the inflammaging-related genes *IL1A, IL1B* and *IL1RN*. The DMR at chr2:113,124,820– 113,125,150 overlaps an ENCODE candidate cis-regulatory element (cCRE) annotated as an enhancer. The upper panel shows the genomic context, with the DMR highlighted in orange, annotated gene promoters in green, and promoter-capture Hi-C interactions linking the DMR to those promoters as arcs. The lower panel shows the DMR itself for myeloid-assigned reads (left) and aggregated lymphocyte-assigned reads (right). Methylation tracks give per-CpG methylation levels in the six deep-sequenced samples, two from each group. Histone tracks give H3K4me1, H3K4me3, and H3K27ac signal in myeloid cells and lymphocytes from ROADMAP and BLUEPRINT, supporting an active enhancer state in myeloid cells but not in lymphocytes.

**Fig. 5.**
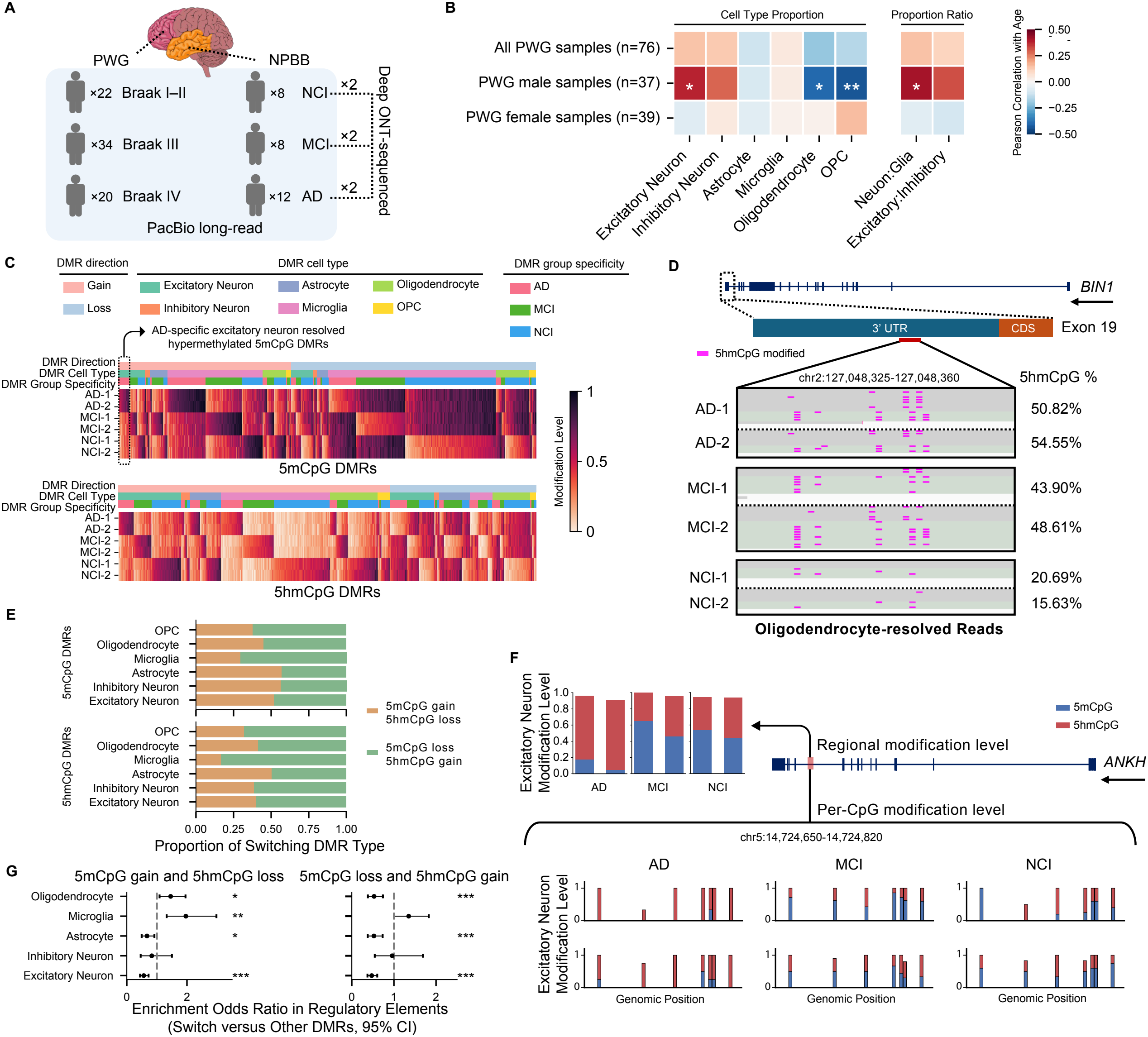
Cell-type composition change and cell-type-resolved modification in brain. **(A)** Cohort design and sequencing strategy. The PWG cohort comprised 76 prefrontal cortex (PFC) samples spanning Braak stages I–II (n = 22), III (n = 34), and IV (n = 20). The NPBB cohort comprised 28 temporal lobe samples adjacent to the hippocampus, including donors with no cognitive impairment (NCI; n = 8), mild cognitive impairment (MCI; n = 8), and AD (n = 12). All samples were sequenced on PacBio, and two samples from each NPBB group, one younger and one older donor with ages approximately matched across groups (younger, 65–66 years; older, 73–79 years), were additionally sequenced to high depth on ONT. **(B)** Associations between brain cell-type proportions or proportion ratios and age, shown for all PART samples (n = 76), male samples (n = 37) and female samples (n = 39). Colors indicate Pearson correlation coefficients (Pearson’s *r*). Asterisks indicate Braak stage-adjusted Pearson correlation *P* values: \**P*_adj_ < 0.05, \*\**P*_adj_ < 0.01. **(C)** Group-specific, cell-type-resolved 5mCpG and 5hmCpG DMRs identified from the six ONT-sequenced NPBB samples, two per clinical group. DMRs were defined by a difference in modification level of at least 0.2 between the two samples of one clinical group and the remaining four. Heatmap rows represent samples, columns represent DMRs, and color indicates modification level. The annotation bars above each heatmap give the direction of modification change, the cell type in which the DMR was identified, and the clinical group to which it is specific. **(D)** Representative group-specific, cell-type-resolved 5hmCpG DMR in the 3′ UTR of exon 19 of *BIN1* (chr2:127,048,325–127,048,360), identified as NCI-specific in oligodendrocyte-resolved reads. Magenta marks in the read-level tracks indicate modified 5hmCpGs, and percentages give the regional 5hmCpG level in each sample. The arrow under the gene name indicates the orientation of *BIN1*. **(E)** Classification of cell-type-resolved DMRs by modification pattern, shown for 5hmCpG DMRs (top) and 5mCpG DMRs (bottom). Left panels give the proportion of each cell type’s DMRs that are non-switching gain, non-switching loss, or switching. Right panels give the proportion of the switching DMRs falling into each of the two switching configurations, 5mCpG gain with 5hmCpG loss, and 5mCpG loss with 5hmCpG gain. **(F)** Representative switching DMR in *ANKH* (chr5:14,724,650–14,724,820), illustrating opposing 5mCpG and 5hmCpG changes that can cancel at the combined modification level. Bars at the upper left give regional 5mCpG and 5hmCpG levels for the six ONT-sequenced NPBB samples, two per clinical group. Lower panels give per-CpG 5mCpG and 5hmCpG levels across the region for each sample in each group. The arrow under the gene name indicates the orientation of *ANKH*. **(G)** Enrichment of each switching DMR type in regulatory elements relative to other DMRs, shown for 5mCpG gain with 5hmCpG loss (left) and 5mCpG loss with 5hmCpG gain (right) across cell types. Points give odds ratios and bars give 95% confidence intervals, with the dashed line marking an odds ratio of 1. Asterisks indicate statistical significance: \**P* < 0.05, \*\**P* < 0.01, and \*\*\**P* < 0.001.

To test whether variation in sequencing coverage might confound the quantification of cell-type proportion and cell-type-resolved methylation, we selected the highest-depth sample from each tissue cohort and downsampled it 2,000 times to each of three targets, two intermediate depths and the lowest depth observed in that cohort. Cell-type proportions and cell-type-resolved methylation levels estimated from the downsampled data closely recapitulated those obtained at full depth across all replicates, depth targets, and tissues (**Supplementary Fig. 6A–C**), indicating that COMET recovers both measures reliably even at the lowest coverage represented in our cohorts.

**Fig. 6.**
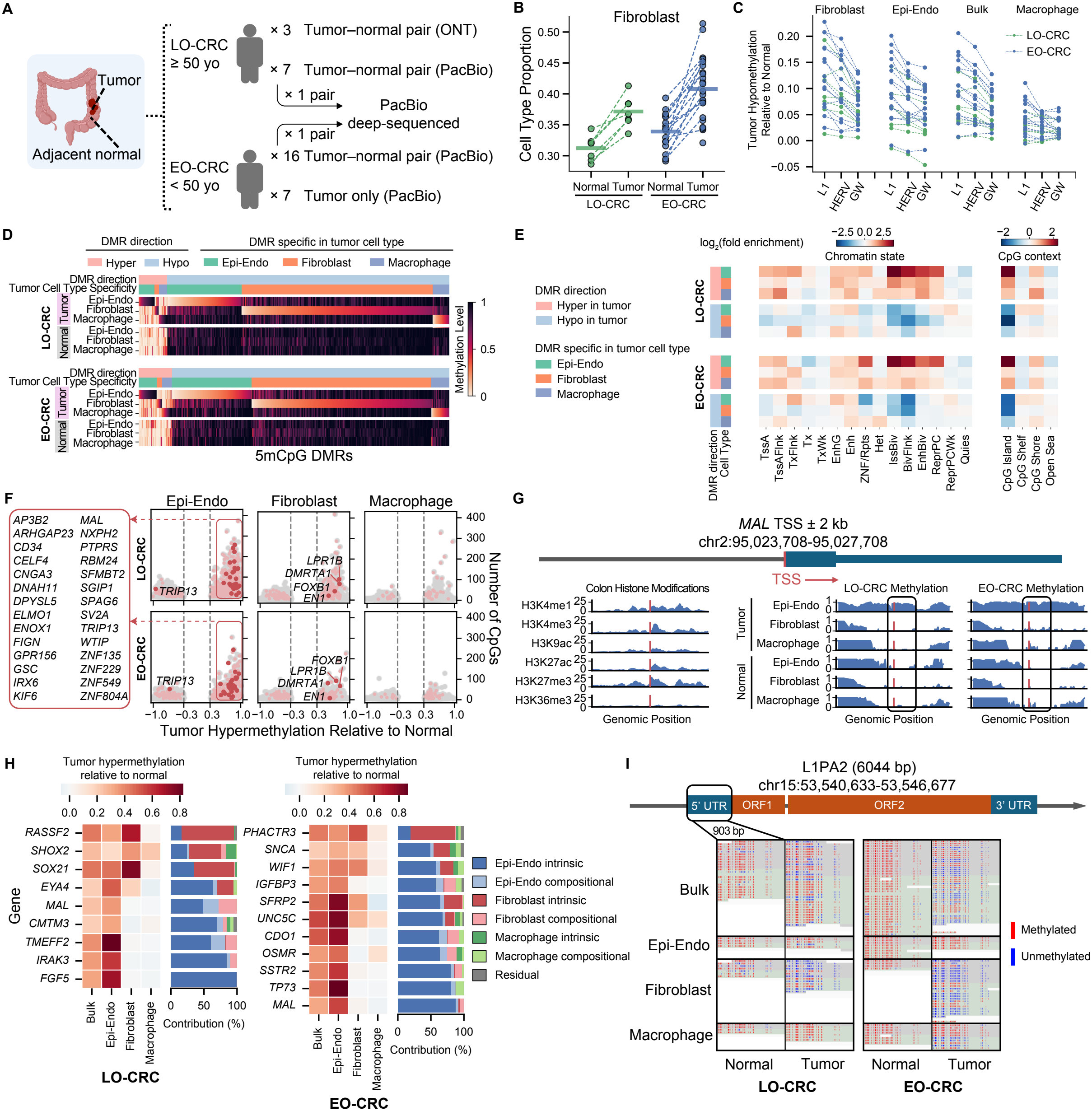
Cell-type composition and cell-type-resolved methylation changes in CRC. **(A)** Cohort design and sequencing information for colon samples. Tumor and adjacent normal colon tissues were analyzed. Ten tumor–normal pairs from individuals aged ≥50 years, defined as late-onset CRC (LO-CRC), were sequenced by ONT (n = 3 pairs) or PacBio (n = 6 pairs). Seventeen tumor–normal pairs from individuals aged <50 years, defined as early-onset CRC (EO-CRC), and seven additional tumor-only samples were sequenced by PacBio. One LO-CRC pair and one EO-CRC pair were further deep-sequenced by PacBio. **(B)** COMET-estimated fibroblast proportions in PacBio-sequenced tumor and matched normal samples. Y-axis shows fibroblast proportions of PacBio-sequenced samples, the dashed lines connect matched normal and tumor samples from the same subject. Green and blue indicate LO-CRC and EO-CRC, respectively. Horizontal lines indicate group means. **(C)** Loss of methylation in tumor relative to matched adjacent normal, calculated from COMET-estimated bulk and cell-type-resolved methylation levels in the PacBio-sequenced pairs. The methylation loss (y-axis) is shown for bulk tissue and each cell type across genome-wide (GW), L1 and HERV regions (x-axis), with positive y-axis values indicating tumor hypomethylation relative to matched adjacent normal. Dashed lines connect measurements from the same tumor–normal pair. Green and blue dots indicate LO-CRC and EO-CRC, respectively. **(D)** Overview of tumor- and cell-type-specific differentially methylated regions (tctDMRs) identified in the deep-sequenced LO-CRC and EO-CRC pairs. A tctDMR was defined as a region where one cell type in the tumor differed in methylation level by at least 0.3 from every other cell type in that tumor and from all cell types in the matched normal tissue. The upper and lower heatmaps show the LO-CRC and EO-CRC pairs, respectively. Each column represents a 5mCpG DMR, and each row represents the methylation level of a cell type in tumor or matched normal tissue. Annotation bars above each heatmap indicate the direction of DMR and the tumor cell type in which the DMR is specific. **(E)** Enrichment of chromatin states and CpG contexts among tctDMRs in the LO-CRC and EO-CRC pairs. Columns represent annotation categories, and rows represent tctDMR groups defined by methylation direction and tumor cell-type specificity. Heatmap colors indicate log_2_(fold enrichment). **(F)** Promoter-mapped tctDMRs shared between the LO-CRC and EO-CRC pairs. Each dot represents a tctDMR, with the gain of methylation in tumor on the x-axis and the number of CpGs on the y-axis. Upper and lower panels show the LO-CRC and EO-CRC pairs, respectively, with columns corresponding to Epi-Endo, fibroblast, and macrophage tctDMRs. Positive values on the left indicate hypermethylation in tumor and negative values on the right indicate hypomethylation. Dark red indicates promoter-mapped tctDMRs shared between the LO-CRC and EO-CRC pairs, with the associated genes labeled; light red indicates other promoter-mapped tctDMRs, and grey indicates other tctDMRs. Vertical dashed lines indicate the methylation-difference cutoffs used for DMR identification. **(G)** Example of a promoter-mapped tctDMR at the *MAL* locus. Colon histone-modification profiles and cell-type-resolved methylation profiles from the matched normal and tumor samples of the LO-CRC and EO-CRC pairs are shown across the *MAL* TSS ± 2 kb (chr2:95,023,708– 95,027,708). The arrow under the gene name indicates the orientation of *MAL*. **(H)** Bulk and cell-type-resolved methylation increase and cell-type contributions to bulk hypermethylation in tumor for candidate promoters associated with CRC screening genes. For each panel, the heatmap shows the methylation increase in tumor relative to matched normal tissue for bulk tissue and individual cell types, with rows representing genes and positive values indicating methylation gain in tumor. Adjacent stacked bars show the contribution of each cell type to the bulk methylation increase in tumor, separated into intrinsic methylation changes (dark shades) and compositional changes (light shades), with grey indicating the residual. LO-CRC and EO-CRC results are shown on the left and right, respectively. **(I)** Example of an L1-mapped tctDMR. Read-level methylation patterns across the 5′ UTR of an L1PA2 element at chr15:53,540,633–53,546,677 are shown for bulk tissue and three cell types in matched normal and tumor samples. IGV views are shown for the LO-CRC pair (left) and EO-CRC pair (right), with methylated and unmethylated CpGs indicated in red and blue, respectively.

Next, we evaluated COMET using real samples whose expected cell-type content is known (**Fig. 1E–G, Supplementary Table 2**, and **Methods**). For blood, we compared whole-blood samples with seven peripheral blood mononuclear cell (PBMC) samples profiled by PacBio sequencing. Because PBMC preparation removes granulocytes, PBMCs are expected to carry fewer myeloid cells and a correspondingly higher fraction of lymphocytes, B cells, NK cells, and T cells, than whole blood. COMET recovered exactly this pattern, assigning PBMCs a higher lymphocyte proportion and a lower myeloid proportion than whole blood (**Fig. 1E** and **Supplementary Fig. 7A**). The seven PBMC samples comprised one primary preparation and six preparations derived from induced pluripotent stem cells (iPSCs); as expected, the iPSC-derived samples carried a higher proportion of naive cells (B Naive and T Naive) and a lower proportion of antigen-experienced cells (B Memory and T Ag-exp) than the primary sample (**Supplementary Fig. 7A**). In brain, we examined 11 ONT-sequenced samples, comprising six bulk brain samples and five samples of defined cell-type composition: one GABAergic neuron sample, one iPSC-derived neuron sample, one sorted microglia sample, one iPSC-derived microglia sample, and one mixed-glia sample. For these samples, we additionally applied lineage-level assignment, which groups reads that cannot be resolved to a single cell type into the neuronal or macroglia lineage (**Methods**). Assigned compositions matched the known identity of every sample. The two neuronal samples both showed the highest neuron proportions, but differed in an informative way: the GABAergic sample was dominated by inhibitory neurons, while the iPSC-derived sample carried a larger share of reads that could not be confidently resolved between inhibitory and excitatory identity (**Fig. 1F** and **Supplementary Fig. 7B**), consistent with an immature population captured before fate commitment. The mixed-glia sample showed elevated glial content, and the sorted and iPSC-derived microglia samples carried the highest microglia proportions of all (**Fig. 1F** and **Supplementary Fig. 7B**). Taking advantage of the high depth of the GABAergic neuron and mixed-glia samples, we mixed reads from the two in defined, graded proportions to build a neuron-to-glia titration series from real data; COMET recovered the imposed composition gradient across the series, confirming on real biological samples the quantification accuracy established in simulation (**Supplementary Fig. 7C**). In colon, we compared colon tissue samples with a colon fibroblast cell line: as anticipated, the cell line showed a higher fibroblast proportion than any bulk sample, a contrast recovered consistently between PacBio and ONT data (**Fig. 1G** and **Supplementary Fig. 7D–E**).

In addition, to evaluate whether COMET captures relative differences in cell type proportions across samples, we used an independent set of 10 blood samples profiled by both cytometry by time-of-flight (CyTOF) and PacBio sequencing (**Methods**). COMET-estimated proportions are correlated with CyTOF-estimated proportions across samples within each cell type (**Supplementary Fig. 8A**; Spearman’s ρ ≥ 0.65, *P* < 0.05), supporting the use of COMET for comparing cell-type composition across samples. We also compared two ways of assigning reads to cell types: using only a small set of highly cell-type-specific CpGs on each read, or integrating all CpGs on the read using a probabilistic model (**Methods**). The two strategies gave similar cell-type proportions and similar assignment confidence, but integrating all CpGs assigned substantially more reads (**Supplementary Fig. 8B–D**).

### COMET recovers a genome-wide, cell-type-resolved methylation landscape, including repeat elements

We first characterized the genomic landscape resolvable by COMET at cell-type resolution. For each cell type, we mapped the genomic intervals at which reads could be assigned to a cell type and recorded how many of that tissue’s cell types each interval could distinguish (**Supplementary Fig. 9**). The pattern is not uniform along the genome, since it depends on how cell-type-specific the methylation at a locus is. The resolvable genomic intervals are distributed across every chromosome, and a large share of them separate more than one cell type. The unresolved gaps correspond largely to centromeric and acrocentric sequence, regions inaccessible to the short-read-based reference panels themselves.

Cell-type composition and cell-type-resolved methylation were stable across shallow-sequenced samples within each tissue, indicating that COMET’s assignment reports a property of the tissue rather than of the individual library (**Fig. 2A** and **Supplementary Tables 3–5**). The recovered methylation levels reproduced established biology. In blood, the two naive cell types (B Naive and T Naive) carried the highest genome-wide methylation, the two antigen-experienced cell types (B Memory and T Ag-exp) the lowest, with myeloid and NK cells in between (**Fig. 2A** and **Supplementary Table 3**), recapitulating the progressive methylation loss known to accompany lymphocyte differentiation^26-28^. In brain, inhibitory neurons carried a higher combined modification level (5mCpG plus 5hmCpG) than excitatory neurons or glia (**Fig. 2A** and **Supplementary Table 4**), consistent with prior work^18,29,30^. In colon, fibroblasts were the least methylated of the three cell types^2^ (**Fig. 2A and Supplementary Table 5**).

A reference-panel method is normally confined to the sites that define the panel. Long reads lift that constraint, because a read anchored at panel sites carries modification calls at every CpG it covers. Across the deep-sequenced samples, this yielded more than 2 million additional cell-type-resolved CpGs per blood sample, more than 1 million per brain sample, and more than 3 million per colon sample, all outside the reference panel itself (**Fig. 2B**). A large share of these additional CpGs fall inside repeat elements, which are usually hard to resolve with short-read or microarray-based methods (**Fig. 2B**). Transposable elements (TEs) such as LINE and SINE elements accounted for the majority in every tissue, roughly in proportion to their share of the genome (**Fig. 2B**).

Because repeat element methylation has been largely inaccessible at cell-type resolution due to limitations of array-based and short-read methods, we used these long-read-resolved CpGs in deep-sequenced samples to quantify cell-type-resolved modification levels for the major TE families alongside the genome-wide value (**Fig. 2C–D**). TEs carried higher 5mCpG than the genome-wide average across every cell type, consistent with their constitutive silencing (**Fig. 2C–D**). SVA (SINE-VNTR-Alu) elements carried the highest 5mCpG level and the smallest variation across cell types whereas L1 and HERV showed greater variation (**Fig. 2C–D**), consistent with the evolutionarily recent origin and ongoing retrotransposition activity of SVAs, as well as their strong epigenetic repression across human tissues^31-33^. In blood and colon, the 5mCpG rank order of cell types was preserved from the genome-wide value to repeat families, with naive cell types above antigen-experienced cells in blood, and fibroblasts below Epi-Endo and macrophages in colon (**Fig. 2C**), matching the cell-type ordering seen in the shallow-sequenced cohort (**Fig. 2A**).

Because Nanopore sequencing can detect 5mCpG, 5hmCpG as separate marks, we resolved them by cell type and repeat family in the brain samples (**Fig. 2D**). Genome-wide, non-CpG (CpH) methylation accumulated in neurons and was near-absent in glia, with higher levels in inhibitory neurons than in excitatory neurons (**Fig. 2D**), matching the neuronal accumulation of 5mCpH and its cell-type-specific distribution described in human cortex^15,30^. For 5hmCpG, excitatory neurons carried the highest levels while oligodendrocytes had lower levels (**Fig. 2D**), consistent with the ordering reported among sorted human excitatory neurons, inhibitory neurons, and oligodendrocytes^30^. Across the six cell types, microglia carried the highest 5mCpG and neurons the lowest, with the reverse relationship for 5hmCpG and 5mCpH (**Fig. 2D**), recapitulating patterns previously observed between neurons and microglia in mouse brain^18,34^. Extending this analysis to repeat families revealed mark-specific patterns: the 5hmCpG cell-type ordering held within every repeat family, whereas Alu elements carried higher 5hmCpG levels than the genome-wide average, and L1 and HERV elements carried lower levels (**Fig. 2D**). For 5mCpH, the signal concentrated in Alu and SVA and was lowest in L1 (**Fig. 2D**).

COMET’s read-level resolution also allows direct visualization of cell-type-resolved reads at individual loci from deep-sequenced data. We examined loci where the expected answer is already established by independent evidence and visualized at read level in a representative deep-sequenced sample for each tissue (**Fig. 2E–F** and **Supplementary Fig. 10A–B**). In blood, *MPO* encodes myeloperoxidase, an enzyme stored in the azurophilic granules of neutrophils and monocytes; it is among the most lineage-restricted genes in hematopoiesis, used clinically as a myeloid marker, and known to be hypomethylated in myeloid cells relative to lymphoid cells^35,36^. Cell-type-resolved reads showed this pattern as expected: myeloid-assigned reads were mostly unmethylated across the region, while reads assigned to all five lymphoid classes were methylated (**Fig. 2E, Supplementary Fig. 11A**). It is worth noting that, even though the five lymphoid cell types do not appear different within the displayed region, they can be resolved based on the CpG methylation status on the flanking regions of the reads (**Supplementary Fig. 12**). In brain, *GPR17* marks OPC and newly formed, pre-myelinating oligodendrocytes and is downregulated as oligodendrocytes mature, making it a marker of a transition state rather than a terminal identity^37,38^; OPC-assigned reads carried the lowest modification level of the six cell classes at this locus, while other cell types remained uniformly modified (**Supplementary Fig. 10A**). Independent samples of known identity supported this pattern, with the same *GPR17* region largely modified in neuron and microglia populations (**Supplementary Fig. 11B**). A second brain example shows how separating 5mCpG from 5hmCpG, a distinction inaccessible to bisulfite-based assays, since both marks resist bisulfite conversion and are therefore read as the same signal, can expose cell-type-specific regulatory biology that would otherwise stay hidden. *NEUROD1* encodes a neurogenic transcription factor central to excitatory neuronal differentiation; at a candidate regulatory region near its transcription start site (TSS), we observed pronounced 5hmCpG specifically on excitatory-neuron-assigned reads, while reads from other cell types remained largely unmodified (**Fig. 2F**). This pattern fits the established role of 5hmCpG in the nervous system, where it is deposited by TET-family enzymes and accumulates preferentially at the regulatory regions and gene bodies of actively transcribed neuronal genes in a cell-type-specific, expression-correlated manner^18,39^. The enrichment of 5hmCpG to excitatory-neuron-assigned reads points to a cell-type-restricted, TET-mediated regulatory state at *NEUROD1* that tracks its lineage-specific expression. The same region were largely unmodified in independent GABAergic neuron, iPSC-derived microglia and mixed glia samples (**Supplementary Fig. 11C**). In colon, *FZD1* encodes a receptor linked to fibroblast activation and expressed predominantly in colonic stroma^40,41^; fibroblast-assigned reads were markedly less methylated across the region than Epi-Endo- or macrophage-assigned reads (**Supplementary Fig. 10B**). Consistently, reads over this *FZD1* region in the colon fibroblast cell line were mostly unmethylated (**Supplementary Fig. 11D**).

### Cell-type compositional change accounts for most of the blood epigenetic aging

We sequenced blood samples from 80 individuals with no recorded disease diagnosis in the BioMe cohort^42^, spanning ages 2 to 83 (**Fig. 3A** and **Supplementary Table 6**). All samples were sequenced on PacBio, and for 47 samples we also generated matched methylation array data from the same blood draw (**Fig. 3A** and **Supplementary Table 6**). We wanted to test whether blood epigenetic aging reflects a cell-intrinsic methylation program or a shift in cell type composition. To test this, we first confirmed the expected bulk signal: genome-wide bulk methylation declined significantly with age (**Fig. 3B**; Pearson’s *r* = −0.447, *P* = 3.30×10^−5^), consistent with the progressive global hypomethylation long associated with blood aging. When we resolved the same analysis into the six constituent cell types, however, cell-type-resolved correlation between methylation and age was much weaker than the bulk correlation, and only B Naive and B Memory reached nominal significance (**Fig. 3B**; B Naive: Pearson’s *r* = −0.223, *P* = 0.047; B Memory: Pearson’s *r* = −0.307, *P* = 0.006; other four cell types: |Pearson’s *r*| < 0.1, *P* > 0.05).

We then tested how well cell-type composition change accounts for the bulk decline in DNA methylation. Two naive populations, B Naive and T Naive, declined significantly in proportion with age (**Fig. 3C**; B Naive: Pearson’s *r* = −0.534, *P* = 3.35×10^−7^; T Naive: Pearson’s *r* = −0.618, *P* = 1.03×10^−9^), while two antigen-experienced populations, B Memory and T Ag-exp, increased with age (**Fig. 3C**; B Memory: Pearson’s *r* = 0.456, *P* = 2.14×10^−5^; T Ag-exp: Pearson’s *r* = 0.477, *P* = 7.61×10^−6^). NK proportion declined modestly (**Fig. 3C**; Pearson’s *r* = −0.302, *P* = 0.007), and myeloid proportion stayed essentially flat (**Fig. 3C**; Pearson’s *r* = 0.191, *P* = 0.09). Because naive cells carry higher methylation level than their antigen-experienced counterparts, this steady replacement of naive by experienced cells explains why bulk methylation declines significantly even though no single cell type shows a comparably strong change of its own methylation level.

To quantify the relative contributions of cell-type composition and intrinsic change to the bulk decline, we partitioned the age-related decrease in bulk methylation into a compositional term and an intrinsic term for every cell type (**Methods**). Blood samples were grouped into four age bins ([0,20), [20,40), [40,60), and ≥60), and each comparison between consecutive bins was quantified this way. In all three comparisons, the compositional term accounted for the largest share of the bulk decline (56.19%–97.37%; **Fig. 3D** and **Supplementary Table 7**). The intrinsic term was small in the [0,20) to [20,40) and [40,60) to ≥60 comparisons, but contributed 41.80% between the [20,40) and [40,60) groups, where the decline in bulk methylation was largest. Above age 60, bulk methylation changed little. Within the compositional term, B Naive, T Naive, B Memory and T Ag-exp, the four cell types with the largest age-related change in proportion, accounted for most of the contribution (88.20%–99.54%; **Fig. 3D** and **Supplementary Table 7**). Together, these results indicate that composition is the main contributor to the age-related decline in bulk methylation, with a smaller within-cell contribution that is clearest in middle age.

We next asked whether this compositional change also explains the behavior of the Horvath epigenetic clock. For the 47 samples with matched array data, array-based global methylation declined with age (**Supplementary Fig. 13A;** Pearson’s *r* = −0.444, *P* = 0.002), reproducing the relationship seen in the long-read data, and Horvath’s clock age tracked chronological age well (**Supplementary Fig. 13B**). Horvath’s clock is highly correlated with all six cell-type proportions, most strongly with T Naive and B Naive (**Fig. 3E** and **Supplementary Fig. 13C**), and with bulk methylation level (**Fig. 3E** and **Supplementary Fig. 13D**). Associations between Horvath’s clock and cell-type-resolved methylation were weaker and largely non-significant, reaching significance only for B Naive and T Ag-exp (**Fig. 3E** and **Supplementary Fig. 13D**). Horvath epigenetic age therefore behaves more like a readout of the shifting cellular mixture than of a cell-intrinsic methylation program.

To see what these two mechanisms look like at individual loci and single-molecule resolution, we selected two young (24 and 25 years old) and two old (75 and 78 years old) male blood samples for deep sequencing: the two older samples showed lower bulk methylation level, lower naive cell proportions, and higher experienced cell proportions compared to the two younger ones (**Fig. 3A** and **Supplementary Fig. 14A–B**). From these four samples we generated high-coverage data and examined cell-type-resolved reads spanning ±500 bp around the Horvath clock CpGs (**Fig. 3F–G**). Both mechanisms were evident within the clock CpG set.

At the region surrounding two clock CpGs, cg03682823 and cg18139769, NK-resolved and T Naive-resolved reads carried clearly different methylation levels in all four samples, while within each cell type, young and old reads looked alike (**Fig. 3F)**; bulk methylation change at loci like these depends mainly on shifting proportions of the cell types involved. By contrast, at the region surrounding a third clock CpG, cg09118625, reads assigned to myeloid cells differed between young and old samples, an intrinsic, age-related methylation change (**Fig. 3G)**.

We then quantified both terms across the full set of Horvath clock CpGs we could evaluate (n = 346). For each CpG, we used the mean absolute methylation difference between cell types within a sample as a measure of compositional difference, and the mean absolute methylation difference within a cell type between young and old samples as a measure of intrinsic difference (**Methods**). The compositional difference corresponding to **Fig. 3F** exceeded the intrinsic difference corresponding to **Fig. 3G** for the majority of clock CpGs, with most points falling well above the identity line (**Fig. 3H**). These observations suggest that a substantial part of the Horvath clock signal in blood reflects CpGs that distinguish one cell type from another.

### Compositional and myeloid-specific methylation changes mark APOE4 risk and cognitive resilience

APOE4 is a major genetic risk factor of Alzheimer’s disease (AD) and is thought to accelerate neurodegenerative disease in part through heightened systemic inflammation, with effects traceable to specific immune cell programs^43^, making cell-type-resolved blood profiling well suited to characterizing its peripheral correlates. We included 60 APOE4-carrier blood samples from the BioMe biobank and Davos Alzheimer’s Collaborative (DAC), alongside the 80 APOE4 non-carrier samples used in the aging analysis above (**Fig. 4A** and **Supplementary Table 6**). Within this cohort, we identified a subset of APOE4 homozygous (APOE44) individuals aged 75 or older with no neurodegenerative disease (NDD) diagnosis on record; reviewing their plasma p-tau217 and neurofilament light (NfL) values, we defined resilience candidates as those with neither an NDD diagnosis nor elevated p-tau217 and NfL, yielding 7 such individuals (**Fig. 4A, Supplementary Fig. 15A, Supplementary Table 6**, and **Methods**). We compared these resilience candidates with two age-matched groups drawn from the same cohort: 9 non-resilient APOE4 homozygous carriers and 6 APOE4 non-carrier controls (**Fig. 4A**).

We first confirmed that the carrier population reproduced the aging pattern seen in non-carriers, including the decline of bulk methylation with age and the naive-to-antigen-experienced compositional drift (**Supplementary Fig. 15B–C**), while additionally showing an age-associated methylation decline in B Memory and T Ag-exp that was not present in non-carriers (**Fig. 3B** and **Supplementary Fig. 15C**). Because carriers skewed older and were distributed less evenly across age than non-carriers, a direct comparison between the two groups would confound APOE4 genotype with age; to account for this, we grouped both populations into three matched age bins ([20,40), [40,60), ≥60) and compared genotypes only within each bin. In every bin, carriers showed the same features as aged samples in the broader cohort, relative to age-matched non-carriers: lower bulk methylation (*P*_adj_ < 0.01), lower B Naive and T Naive proportions and higher B Memory and T Ag-exp proportions (*P*_adj_ < 0.05), while myeloid and NK proportions remained indistinguishable (**Fig. 4B**). APOE4 carriers additionally showed reduced methylation at the level of individual cell types. The reduction was observed across all six cell types and was more pronounced in the ≥60 age group, which showed stronger difference than the other two age groups (**Supplementary Fig. 16A**). Together, this suggests APOE4 carriers have an accelerated aging in terms of cell-type compositional changes, along with an intrinsic, within-cell-type methylation loss.

Building on these aging-associated cell-type signatures, we examined the same features across the three defined groups of individuals aged 75 and older: non-carrier controls, non-resilient APOE4 carriers, and resilience APOE4 carriers (**Fig. 4A**). Bulk methylation in resilience candidates was significantly higher than in non-resilient carriers (*P*_adj_ < 0.05) but statistically indistinguishable from non-carrier controls (**Fig. 4C**). Cell-type composition showed the same pattern: resilience candidates carried higher B Naive and T Naive proportions and lower B Memory and T Ag-exp proportions than non-resilient carriers (*P*_adj_ < 0.05) and were statistically indistinguishable from non-carrier controls on every measure (**Fig. 4C**). Per-cell-type methylation, by contrast, was largely indistinguishable across the three groups, the one exception being higher B memory methylation in resilience candidates than in non-resilient carriers (**Supplementary Fig. 16B**). Composition, rather than global within-cell methylation state, therefore appears to be what separates resilience APOE4 carriers from their at-risk peers.

To further characterize cell-type-specific methylation at individual loci, we deep-sequenced 6 samples, two from each group, and called group-specific differentially methylated regions (DMRs) within each blood cell type (**Fig. 4D, Supplementary Table 8**, and **Methods**). Myeloid cells yielded more group-specific DMRs than any lymphocyte cell type (**Fig. 4D**), consistent with the larger fraction of reads assigned to myeloid cells (**Fig. 1E and Fig. 2A**), which gives more statistical power to detect DMRs in this cell type. Within myeloid cells, both non-resilient and resilient APOE4 carriers yielded more DMRs than non-carrier controls, with resilience carriers yielding the most. The direction of these DMRs was asymmetric: non-carrier-specific myeloid DMRs were predominantly hypermethylated, while resilient-specific myeloid DMRs were predominantly hypomethylated, relative to the other two groups (**Fig. 4D**). The two layers of APOE4-associated difference therefore map to different cell compartments: the compositional signal that separates carriers from non-carriers is found in lymphocytes, whereas deep sequencing resolved locus-level, group-specific methylation differences in myeloid cells.

Annotating the group-specific myeloid DMRs against myeloid histone modification data and promoter-capture Hi-C (PCHi-C) interactions (**Methods**), we found many to fall within regions marked as enhancers or promoters linked by PCHi-C to genes (**Fig. 4E** and **Supplementary Table 9**). The genes these regions mapped onto included several previously reported inflammaging markers: *CTSW, IKZF1*, and *NEAT1* among resilient-specific DMRs, and *IL1A, IL1B, IL1RN, PIM1*, and *GPR17* among non-resilient-specific DMRs (**Fig. 4E**), pointing to gene-level effects on inflammaging regulation. As one example, the non-resilient-specific hypomethylated region chr2:113,124,820-113,125,150 is annotated as a candidate cis-regulatory (cCRE) enhancer within *IL1RN* and is supported as an active enhancer by myeloid histone modification data from both the ROADMAP^20^ and BLUEPRINT^19^ consortia (**Fig. 4F, Supplementary Fig. 17A–B**, and **Methods**). In myeloid PCHi-C, the enhancers annotated in this region contact the promoters of several nearby genes, including the IL-1 gene cluster (*IL1A, IL1B, IL1RN*), encoding IL-1α, IL-1β and the IL-1 receptor antagonist, a signaling axis that serves as a central mediator of inflammaging^44^ (**Fig. 4F)**. The same region showed no group difference in aggregated lymphocyte-resolved reads, and lymphocyte histone data carried no active marks across it (**Fig. 4F and Supplementary Fig. 17C–D)**, indicating this is a myeloid-specific enhancer whose methylation differs specifically in non-resilient carriers.

Collectively, these findings point to two layers of APOE4-associated difference in blood: a compositional difference shared across carriers, with fewer naive and more antigen-experienced cells, and a myeloid-specific methylation signature that further distinguishes resilient from non-resilient carriers, at loci supported as active myeloid regulatory elements by ROADMAP and BLUEPRINT histone modification data and PCHi-C interactions.

### Sex-biased brain aging and 5mCpG-5hmCpG switching emerge at cell-type resolution

We sequenced 104 postmortem brain samples from two cohorts (**Fig. 5A and Supplementary Table 10**). To investigate cell-type changes associated with brain aging, we leveraged 76 prefrontal cortex (PFC) samples from donors with primary age-related tauopathy (PART), all with a CERAD score of 0 and spanning Braak stages I–II (n = 22), III (n = 34) and IV (n = 20), obtained from PART Working Group (PWG) cohort (**Fig. 5A** and **Methods**). Because PART is characterized by age-related tau pathology in the absence of substantial amyloid deposition, this cohort provides a valuable setting for studying cellular and epigenetic changes associated with brain aging with minimal confounding from amyloid pathology. The second cohort comprises 28 samples from temporal lobe, adjacent to the hippocampus, from the Mount Sinai Neuropathology Brain Bank (NPBB), spanning the clinical spectrum: 8 donors with no cognitive impairment (NCI), 8 with mild cognitive impairment (MCI), and 12 with AD (**Fig. 5A** and **Methods**).

We first quantified cell-type composition and cell-type-specific methylation levels for every sample in both cohorts and tested them against the available clinical traits. In the NPBB cohort, cell type composition is similar between groups (**Supplementary Fig. 18A**). Methylation level in oligodendrocytes was the only measure that is higher in MCI than in NCI (*P*_adj_ = 0.0306, **Supplementary Fig. 18B**), exhibiting a non-monotonic trajectory across diagnostic groups that aligns with the non-linear, stage-dependent involvement of the oligodendrocyte lineage previously described at the transcriptional level in AD^45,46^. No measure differed across Braak stage groups in the PWG cohort (**Supplementary Fig. 18C–D**). However, stratification of the PWG cohort by sex revealed a pronounced sex bias in age-associated cell-type composition that was not apparent in the pooled analysis, consistent with reported sex stratifications in brain aging trajectories and the potential for pooling across sexes to dilute or mask sex-specific signals^47,48^. Among the male donors (n = 37), neuronal proportion increased with age for excitatory neurons, whereas proportions of the oligodendrocyte lineage decreased, with significant declines in oligodendrocytes and OPC (**Fig. 5B** and **Supplementary Fig. 19A**).

Consistently, the neuron-to-glia ratio also increased with age (**Fig. 5B** and **Supplementary Fig. 19A**). In contrast, none of these age-associated correlations reached significance among female donors (n = 39; **Fig. 5B** and **Supplementary Fig. 19A**). Age-related decline of OPC and myelin integrity in the human brain are well documented^49-51^, and this analysis suggests the decline is more pronounced in male donors. Age-related methylation changes in the PWG cohort were modest across cell types, with significant associations detected only in bulk tissue (**Supplementary Fig. 19B**). Neither the age associations nor the sex bias was apparent in the NPBB cohort (**Supplementary Fig. 19C**), which may reflect its smaller sample size, different donor ascertainment, and the distinct cell-type composition of the brain region sampled.

To illustrate the study of cell-type specific DNA methylation changes at individual loci in AD, we selected two samples, one younger and one older donor with ages approximately matched across groups (younger, 65–66 years; older, 73–79 years), from each NPBB group and sequenced them to high coverage **(Fig. 5A)**. Because nanopore sequencing detects 5mCpG and 5hmCpG as distinct modifications, we called group-specific, cell-type-resolved DMRs separately for each mark, yielding two distinct DMR landscapes (**Fig. 5C**). Each landscape was further stratified by cell type and clinical group; the highlighted example shows the AD-specific, excitatory-neuron-resolved, hypermethylated 5mCpG DMRs (**Fig. 5C**). Several of these DMRs fall in genes with established roles in brain function and AD risk (**Supplementary Table 11**). The strong AD risk locus *BIN1* carries a 5hmCpG DMR in the 3’ UTR within exon 19 (chr2:127,048,325-127,048,360), resolved specifically on oligodendrocyte-assigned reads, the cell type in which *BIN1* is predominantly expressed in human brain^52,53^ (**Fig. 5D**). 5hmCpG over this region was 15.6% and 20.7% in the two NCI donors, 43.9% and 48.6% in the two MCI donors, and 50.8% and 54.6% in the two AD donors (**Fig. 5D**).

Because 5hmCpG is derived from 5mCpG via TET-mediated oxidation, we hypothesized that at some loci the two marks might change in opposite directions, a pattern that would be invisible to assays that report their sum such as bisulfite sequencing. To test this, we measured both 5mC and 5hmC at each group-specific and cell-type-resolved DMR. The two were negatively correlated in every cell type and across both DMR sets, with regions losing 5mCpG tending to gain 5hmCpG and vice versa (**Supplementary Fig. 20A–B**). Their combined level, which is what a bisulfite-based assay reports, remained largely stable (**Supplementary Fig. 20C**). We therefore defined switching DMRs as those where both the 5mCpG and the 5hmCpG difference exceeded the 0.2 threshold used for brain DMR identification (**Methods**) and investigated how they are distributed across cell types.

Among the 5mCpG-5hmCpG switching DMRs, 5mCpG loss paired with 5hmCpG gain was the more common pattern, which is consistent with TET-mediated oxidation. This pattern was most dominant in microglia, followed by OPC and oligodendrocytes (**Fig. 5E**). Detecting these modification-specific changes at cell-type resolution depends on measuring 5mCpG and 5hmCpG separately, which native long-read sequencing does directly, and COMET extends this to the cell-type level in bulk samples. As one example, a DMR at chr5:14,724,650-14,724,820, within the AD-associated gene *ANKH*^54-56^, illustrates the 5mCpG and 5hmCpG variations on reads assigned to excitatory neurons (**Fig. 5F**). AD donors carried markedly lower 5mCpG and correspondingly higher 5hmCpG than MCI and NCI donors, yet the combined 5mCpG plus 5hmCpG level was nearly identical across the three groups (**Fig. 5F**). The per-CpG view showed the same reciprocal relationship site by site (**Fig. 5F**), indicating that the cancellation holds at single-CpG resolution as well^57^. Consistent with the positive association between gene-body 5hmCpG and transcription reported within individual brain cell types^18,39^, re-analysis of public snRNA-seq and scRNA-seq data from human cortex (**Methods**) showed *ANKH* is upregulated in AD compared with controls specifically in excitatory neurons (**Supplementary Fig. 21A–B**), the same cell type in which the switching DMR was detected (**Fig. 5F**).

Motivated by the cell-type-specific regulatory patterns described above, we asked whether 5mCpG-5hmCpG switching regions tend to fall in regulatory elements. We annotated the group-specific, cell-type-resolved DMRs against matched cell-type regulatory signal, including H3K4me3, H3K27ac, and ATAC-seq, and compared the two classes of switching region, 5mCpG gain with 5hmCpG loss, and 5mCpG loss with 5hmCpG gain, against all other DMRs (**Methods**). In microglia, switching regions were enriched in regulatory elements, roughly twofold for 5mCpG gain with 5hmCpG loss (**Fig. 5G**), which suggests that the two modifications are more likely to change together at regulatory sites in this cell type. In excitatory neurons and astrocytes, switching regions were instead depleted in regulatory elements (**Fig. 5G**), meaning that at regulatory sites in these cell types, changes tend to involve one modification rather than a coordinated 5mCpG-5hmCpG switching. Inhibitory neurons showed no preference in either direction (**Fig. 5G**). Oligodendrocytes were the only cell type with a direction-biased pattern, enriched in regulatory elements for 5mCpG gain with 5hmCpG loss and depleted for the reverse (**Fig. 5G**), indicating that the two switch directions are not mirror images of a single process and that gaining and losing 5hmCpG are separately regulated.

### Cell-type-resolved methylation reveals tumor-intrinsic and stromal signatures in early- and late-onset colorectal cancer

To examine how cell-type composition and methylation change in CRC, we assembled a cohort of paired tumor and adjacent normal colon tissue spanning both early- and late-onset disease (**Fig. 6A and Supplementary Table 12**). The late-onset group (LO-CRC, diagnosed at 50 or older) comprised 10 patients, all with matched pairs, sequenced on ONT (3 pairs) or PacBio (7 pairs) platforms. The early-onset group (EO-CRC, diagnosed before the age of 50) comprised 23 patients, with 16 matched pairs and 7 tumor-only samples, all sequenced on PacBio. Two pairs, one from an LO-CRC patient and one from an EO-CRC patient, were additionally deep PacBio-sequenced to enable locus-level differential methylation analysis at cell-type resolution (**Fig. 6A**).

Applying COMET across the PacBio cohort, we found a consistent compositional shift: fibroblast proportion was higher in tumor than in matched adjacent normal tissue in every paired sample (**Fig. 6B**), a pattern also reproduced in the ONT data (**Supplementary Fig. 22A**). This mirrors the well-established expansion of cancer-associated fibroblasts (CAFs), the dominant stromal constituent of most solid tumors and an active participant in CRC progression^58^ and shows that COMET recovers a known tumor-biology signal directly from unsorted tissue. The increase was more pronounced in EO-CRC than in LO-CRC (**Supplementary Fig. 22B**), in line with our earlier single-cell RNA sequencing (scRNA-seq) quantification^59^ and with a recent snRNA-seq atlas of colorectal tumors^60^. For the other two cell types, macrophage shows cell type proportion decrease in most pairs, while Epi-Endo did not show a consistent change across pairs (**Supplementary Fig. 22C**).

Genome-wide methylation was lower in tumor than in matched adjacent normal in all pairs for bulk, fibroblast, and macrophage, and in majority of pairs for Epi-Endo (**Supplementary Fig. 22D–E**). This loss was not distributed evenly across the genome or across cell types. Retrotransposable L1 and HERV elements lost methylation more than the genome-wide average, and the loss was more pronounced in fibroblast and Epi-Endo cells than in macrophages (**Fig. 6C** and **Supplementary Fig. 22F**), indicating that the global hypomethylation characteristic of the CRC methylome is not a uniform property of the tissue.

We generated methylation array data for the seven PacBio-sequenced LO-CRC pairs and found that the arrays (**Supplementary Table 12**), which largely exclude repetitive regions by probe design, detected a decrease in average tumor methylation in only five of the seven pairs (**Supplementary Fig. 22G**). This observation is consistent with our long-read-based finding that repetitive regions contribute disproportionately to the tumor hypomethylation signal.

We then moved to individual loci by deep-sequencing two tumor and adjacent-normal pairs, one from an LO-CRC and one from an EO-CRC patient (**Fig. 6A**). These samples recapitulated the cohort-level findings, showing the same fibroblast proportion increase in tumor and the same methylation decrease at both bulk and cell-type level (**Supplementary Fig. 22H**). Within each cell type, we compared tumor against its own matched adjacent normal tissue, a paired design that controls for inter-patient genetic background, and retained regions specifically methylated in one cell type but not the other two (**Fig. 6D** and **Supplementary Table 13)**, which we term tumor- and cell-type-specific DMRs (tctDMRs). Consistent with the global tumor hypomethylation shown in **Fig. 6C**, hypomethylated tctDMRs substantially outnumbered hypermethylated ones (**Fig. 6D** and **Supplementary Fig. 23A)**. The two directions also differed in their cellular origin: Epi-Endo dominated the hypermethylated set, whereas the hypomethylated set was split between Epi-Endo and fibroblasts, with macrophages contributing comparatively few tctDMRs in either direction (**Fig. 6D** and **Supplementary Fig. 23A)**.

Annotating these tctDMRs against colon chromatin states, CpG context, gene-centric features, and repeat elements, and computing fold enrichment by cell type and direction **(Supplementary Table 13** and **Methods)**, we found that hypermethylated tctDMRs were strongly enriched at bivalent chromatin domains (TssBiv, BivFlnk, EnhBiv), repressive polycomb domains (ReprPC), CpG islands, and promoters and 5’ UTRs across all three cell types and both patient groups, with the strongest enrichment in Epi-Endo cells (**Fig. 6E** and **Supplementary Fig. 23B**). Hypomethylated tctDMRs showed the reciprocal pattern in all three cell types: depletion at bivalent chromatin and CpG islands, with a shift toward repeat elements instead (**Fig. 6E** and **Supplementary Fig. 23B**). This focal-hypermethylation-versus-dispersed-hypomethylation asymmetry recapitulates the classic observation that cancer hypermethylation preferentially targets developmentally poised polycomb and bivalent loci, while hypomethylation spreads through late-replicating regions of the genome^61^. Cell-type resolution adds a dimension bulk analysis cannot provide: this regulatory-element hypermethylation is concentrated disproportionately in Epi-Endo cells rather than spread evenly across the tumor microenvironment.

Mapping tctDMRs onto candidate gene promoters (**Supplementary Table 13**), since promoter methylation is a central mechanism of gene silencing in CRC^62,63^, Epi-Endo tumor-hypermethylation formed the largest and most extreme class (**Supplementary Fig. 23C**): 27 genes were hypermethylated in Epi-Endo cells in both patients (**Fig. 6F**). This shared list includes established CRC methylation markers such as *MAL, GSC, PTPRS*, and *RBM24*, alongside genes not previously reported in CRC, including *GPR156, KIF6*, and *ZNF549* (**Fig. 6F**). Fibroblast tumor-hypermethylation formed a smaller shared set of 4 genes, *LRP1B, DMRTA1, FOXB1*, and *EN1* (**Fig. 6F**), indicating that the two cell types run distinct gene-silencing programs in tumor tissue. Macrophages showed far fewer promoter tctDMRs, none shared between the two patients. As one example, the candidate promoter region of *MAL* (TSS ± 2 kb, chr2:95,023,708-95,027,708) carries histone marks of both an active promoter and an enhancer, indicating it is functionally active in colon tissue (**Fig. 6G**). In both patients, this region gained methylation in tumor specifically on Epi-Endo-assigned reads, while fibroblast and macrophage reads from the same samples remained unmethylated (**Fig. 6G**), illustrating that COMET enables profiling tumor- and cell-type-specific methylation signals at a single locus directly from bulk tissue.

To ask whether the cell-type-level methylation changes are consistent with changes in expression, we examined *C5AR2, CEACAM6, ERBB3, LILRB4, PTPRE*, and *TOP2A* which are highly expressed in an epithelial cell state that is enriched in EO-CRC relative to LO-CRC in our recent scRNA-seq data^59^ and which retained sufficient coverage after cell-type resolution in both deep-sequenced samples. In our deep-sequenced EO-CRC sample, five of six genes carried regions with reduced methylation in tumor compared to normal in Epi-Endo-resolved reads, whereas no such hypomethylation was seen at the same regions in the deep-sequenced LO-CRC sample (**Supplementary Fig. 24A**). These observations connect the cell-type-resolved methylation changes we detect from bulk sequencing to transcriptional changes seen in an independent single-cell dataset, and they support the functional relevance of the epithelial methylation changes. We then examined the clinical relevance of these findings by focusing on 65 genes commonly used in current CRC screening panels (**Supplementary Table 14** and **Methods**). Mapping bulk and cell-type methylation onto these gene promoters, 55 of 65 panel genes in EO-CRC and 51 of 65 panel genes in LO-CRC show tumor hypermethylation at the bulk level (**Supplementary Fig. 24B** and **Supplementary Table 14**). Among the subset with sufficient coverage across all three cell types (**Methods**), 9 genes in LO-CRC and 11 in EO-CRC showed bulk-level hypermethylation in tumor, with Epi-Endo reads contributing disproportionately more methylation than the other two cell types at most of these genes (**Fig. 6H**). By quantifying each cell type’s contribution to the tumor-associated bulk hypermethylation, rather than relying on the bulk signal alone, we found that Epi-Endo cells dominate it (**Fig. 6H**). This is consistent with our earlier analyses showing that promoter hypermethylation in colon tumors is predominantly attributable to Epi-Endo cells (**Fig. 6E–F** and **Supplementary Fig. 23B–C**). This is consistent with the design of stool- and blood-based screening assays, which target methylation markers of colorectal neoplasia arising from the colonic epithelium^64,65^. It therefore confirms, at cell-type resolution, that the methylation signals targeted by these panels originate primarily from the intended epithelial compartment.

Turning to repeat elements in the deep-sequenced pairs, we found that most full-length L1 and HERV elements were hypomethylated in tumor relative to matched normal in all three cell types, with the loss most pronounced in fibroblasts (**Supplementary Fig. 25A**). This is consistent with the pattern seen in the shallow-sequenced cohort (**Fig. 6C** and **Supplementary Fig. 22F**) and the annotation result where hypermethylated tctDMRs were depleted at L1 and HERV while hypomethylated tctDMRs showed no such depletion (**Supplementary Fig. 23B**). We also observed that the methylation loss in tumor on L1 and HERV was similar for young and old subfamilies (**Supplementary Fig. 25A**). Beyond gene promoters, tctDMRs themselves fell within L1 and HERV elements (**Supplementary Fig. 25B–C** and **Supplementary Table 13**). At the promoter region of one L1PA2 locus (chr15:53,540,633–53,546,677) for example, bulk tumor methylation loss was evident in both patients, and read-level assignment showed this loss concentrated on fibroblast-assigned reads (**Fig. 6I**). L1 hypomethylation is a long-recognized feature of CRC^66,67^, and because it has been measured almost entirely in bulk tissue, the cell types driving it have remained unresolved. Our data place the stronger signal in fibroblasts, which contribute to the bulk measurement both through their intrinsic hypomethylation and through their expansion in tumor.

Collectively, these results show that COMET separates tumor-intrinsic epithelial reprogramming from stromal and immune contributions within the same bulk tumor sample, at both regulatory loci, in clinically used screening genes, and at retrotransposable elements, without requiring cell sorting of primary tumor tissue.

## Discussion

Cell-type composition and cell-type-specific methylation have long been inseparable from bulk tissue methylation measurements, forcing aging and disease studies to choose between population-scale bulk profiling and small, atlas-style single-cell or sorted-cell cohorts. Here, we show that COMET addresses this challenge from standard bulk long-read sequencing. Applied across blood, colon, and brain, COMET recovered cell-type composition and cell-type-specific methylation from shallow sequencing, cell-type-resolved modification of repetitive DNA by pooling reads across repeat copies, and, from deeper sequencing, cell-type-specific differential methylation between health and disease. In each tissue, this resolution deepened the interpretation of bulk-level DNA methylation changes: blood epigenetic aging traced largely to a shift in cell-type composition, with intrinsic change contributing mainly in middle age; APOE4-associated blood aging separated into a compositional layer shared by carriers and a myeloid-specific intrinsic layer that distinguished resilient from non-resilient carriers; colorectal tumor hypermethylation related to cancer biology concentrated disproportionately in the epithelial compartment rather than the surrounding stroma or immune infiltrate; and Alzheimer’s-associated brain methylation change consisted of reciprocal 5mCpG and 5hmCpG shifts that cancel at the bulk level.

These results point to a general principle: many bulk methylation signals long treated as intrinsic, cell-autonomous change are, at least in part, signals of a shifting cellular population, and disentangling the two requires cell-type resolution rather than deeper bulk sequencing. This distinction matters beyond methodology: it also bears on how we understand disease risk itself. Genetic risk, such as APOE4 genotype, is fixed at conception and cannot by itself account for why some carriers show the aging and methylation signature we describe while others, the resilience candidates, do not; the compositional and myeloid-specific differences we identify sit downstream of genotype and upstream of clinical outcome, in the dynamic, cell-type-resolved layer where genetic risk is translated into phenotype.

COMET complements single-cell and sorted-cell methylation profiling. These methods remain the gold standard for per-cell resolution, and the reference atlases COMET relies on for assignment are themselves products of them, but their per-sample cost limits how many donors can be profiled. COMET extends cell-type resolution from the atlas-building regime, where these methods excel, into more scalable studies using long read sequencing. Because single-cell methylation atlases now span a wide range of human organs^68^, reference panels can be assembled for tissues beyond the three examined here as these resources grow, which would broaden the aging and disease questions COMET can address. Other groups have shown that long-read methylation can be traced to cell type or tissue of origin^69-72^. The present work builds on this concept and extends it beyond cell-type assignment to composition estimation, cell-type-resolved repeat-element methylation, and cell-type-specific differential analysis across three human tissues.

The translational implications differ by tissue but share a common shape: cell-type resolution turns a diffuse bulk signal into a candidate biomarker or mechanistic target tied to a specific cell type. In blood, the myeloid-specific methylation signature that distinguished resilient from non-resilient APOE4 carriers, at loci independently supported as active myeloid regulatory elements, raises the possibility that myeloid inflammaging state, rather than APOE4 genotype alone, could inform individual risk stratification for AD, and that the specific loci involved, including *IL1A, IL1B*, and *IL1RN*, may warrant further study as modifiable targets. In colon, confirming that the bulk methylation signal used by current screening panels originates predominantly from the epithelial compartment supports the biological rationale behind these assays, and cell-type-resolved profiling may help identify panel loci less diluted by stromal and immune contribution in heterogeneous clinical samples. In brain, the discovery that Alzheimer’s-associated methylation change consists of reciprocal 5mCpG and 5hmCpG switching suggests that markers built on total modification level, as most bisulfite-based assays report, may systematically miss disease-associated signal at switching loci; markers that separate the two modifications could recover this signal. More broadly, the sex-biased compositional shift we observed in brain aging illustrates that cell-type composition can itself confound comparisons that are not explicitly stratified or adjusted for it, a consideration relevant to bulk DNA methylation studies of postmortem brain samples.

Several limitations warrant consideration in future work. First, the deep-sequencing analyses in this study, spanning the APOE4 resilience, brain AD, and CRC comparisons, drew on small numbers of samples per group. We present these analyses as demonstrations of how COMET enables cell-type-specific differential methylation analysis from deep bulk long-read sequencing, illustrating the kind of locus-level, cell-type-resolved signal this approach can recover, rather than as fully powered tests of any single disease association. Establishing the specific loci we highlight as robust, generalizable markers of APOE4 resilience, AD progression, or CRC subtype will require replication in larger deep-sequenced cohorts, a goal that becomes increasingly practical as long-read sequencing costs continue to fall. Second, COMET assigns reads only to the cell types in each tissue’s reference atlas, so its resolution is limited to the cell types and states the atlas distinguishes. This current study resolved six cell types in blood and brain and three in colon, and did not capture finer subtypes such as T-cell memory states, neuronal subtypes beyond the excitatory-inhibitory division, or epithelial subtypes in colon. A cell type missing from a panel may also be assigned to the most similar class or left unassigned. Because COMET places no constraint on the number or source of reference cell types, expanding the atlases would allow it to resolve more cell types and assign more reads, without changes to the framework. Third, COMET sets aside reads whose cell-type assignment falls below a confidence threshold rather than forcing a call, which keeps the assigned read set specific but means that composition and methylation estimates are computed on a subset of the sequenced reads. If ambiguous reads are not missing at random with respect to cell state, this could introduce bias; we chose the threshold to maximize the balance between F1 score and assignment rate in simulation, favoring specificity, and the same threshold and reference panels are applied uniformly across samples and conditions within a tissue, so any resulting bias should affect groups comparably rather than manufacture a group difference. Finally, every cohort in this study was profiled cross-sectionally, so we cannot establish the temporal order between a compositional or methylation change and the clinical outcome it accompanies, nor rule out that both arise from a shared upstream cause; longitudinal cohorts, in which the same individuals are profiled over time, would be needed to test whether the compositional and myeloid-specific signatures we describe precede, rather than follow, cognitive change in APOE4 carriers.

## Acknowledgements

We thank all participants in the Mount Sinai BioMe Biobank, as well as the recruiters and staff who assisted in data collection and management. We thank Panos Roussos for providing sorted microglia samples; Kimberly Hartl for isolating the colon fibroblast cell line; Jinglong Zhang, Bin Hu, and Aiqun Li for providing iPSC-derived neuron and microglia samples. We thank HistoGenetics LLC for performing PacBio sequencing, and Hwa Ran Kim (HistoGenetics) for additional guidance on optimizing PacBio library preparation. This work was primarily supported by a seed fund (G.F.) from the Icahn School of Medicine at Mount Sinai. Rigorous detection of 5mC and 5hmC from long read sequencing data is supported by grant no. R01 HG010646 (G.F. and R.K.) from the National Institutes of Health. Association analysis between DNA methylation and aging is supported by grant no. P01 AG047200 (G.F. and J.V.) from the National Institutes of Health. This research was conducted with the support of the Biorepository and Pathology CoRE at Icahn School of Medicine at Mount Sinai. This work was also supported in part through the computational resources and staff expertise provided by the Department of Scientific Computing at the Icahn School of Medicine at Mount Sinai.

## Author Contributions

H.C. and G.F. designed the study and developed COMET. G.F. supervised the study. H.C. performed all data analyses. E.A.M. prepared DNA samples for long-read sequencing and methylation array profiling, constructed sequencing libraries, and performed sequencing. H.C. and Y.Z processed the raw sequencing data. N.K., Z.C., and F.M.E. collected the DAC blood samples, performed chart review analyses for blood samples, and generated the blood CyTOF data. C.N.B., D.A.C., R.B., J.A.G., J.T.G., and D.J.C. collected colon samples from patients with CRC. E.L.T. and J.F.C. collected NPBB brain samples. K.W.F. and J.F.C. collected PWG brain samples. G.F.H., Z.Y., E.B., and J.V. generated the iPSC-derived PBMC samples. A.W.C assisted with BioMe blood sample collection. S.S. and L.C. generated the colon fibroblast cell line. A.K. and S.D. generated the ONT data for GABAergic neurons and mixed glia. Y.F. and M.N. assisted with methods and software testing. H.C. and G.F. wrote the manuscript. Y.F., M.N., E.A.M., J.T.G., E.G., S.M., R.M.K., H.W., G.F.H., J.V., K.W.F., S.S., and L.C. contributed to data interpretation and manuscript editing. All authors reviewed and approved the final manuscript.

## Methods

### Whole blood samples

We profiled 140 whole blood samples, including 134 from the Mount Sinai BioMe Biobank and 6 from the Davos Alzheimer’s Collaborative (DAC) at the Icahn School of Medicine at Mount Sinai (**Supplementary Table 6**). Blood samples were collected under the protocols STUDY-20-01197 and STUDY-22-01203, approved by the Institutional Review Board (IRB) of the Icahn School of Medicine at Mount Sinai. Phenotypic information was obtained from linked electronic health records (EHRs). Sample age was defined as the donor’s age at the date of blood collection, and APOE genotypes were called from whole-exome sequencing (WES) data based on the two defining single-nucleotide polymorphisms (SNPs), rs429358 and rs7412. Further details on the BioMe cohort, including its associated data and IRB approval, have been described previously^42^.

Whole blood was centrifuged at 2,000 × g for 10 min at 4 °C to separate plasma, and the plasma-depleted fraction was flash-frozen in liquid nitrogen and stored at −80 °C. Plasma was further centrifuged at 12,000 × g for 2 min at 4 °C to pellet exosomal material, and both the pellet and supernatant were flash-frozen. Genomic DNA was isolated from aliquots of plasma-depleted blood, typically 1.5 mL, with the Wizard Genomic DNA Purification Kit (Promega), following the whole blood sub-procedure (3A) of the manufacturer’s protocol including the optional RNase treatment, and scaling up the volumes at each step for larger inputs. DNA was eluted in 0.1× IDTE (IDT), prepared as a 1:9 dilution of IDTE stock in nuclease-free ultrapure water (Thermo Fisher Scientific).

### PBMC samples

Primary PBMCs were obtained from ZenBio (Durham, cat# SER-PBMC-200, lot no. PBMC012721B) and contained more than 2×10^8^ cells. DNA was isolated using the DNeasy Blood and Tissue kit (Qiagen, cat# 69504) according to the manufacturer’s protocol, including the optional RNase A treatment, and eluted in AE buffer.

Induced pluripotent stem cell (iPSC) lines were generated from PBMCs collected from one 24-year-old female donor and one 74-year-old female donor and stored in liquid nitrogen. PBMCs were thawed one day before sorting and cultured in Week 1 medium consisting of StemSpan SFEM II (cat# 09655) supplemented with 1 µM hydrocortisone (cat# H4001), 50 ng/mL SCF (cat# 300-07), 16.6 ng/mL FLT3L (cat# 300-19), 6.6 ng/mL IL-3 (cat# 200-03) and 1.3 U/mL EPO (cat# 55513-0126-10). After 18 to 24 h, CD49f? long-term hematopoietic stem cells (LT-HSCs), defined as Lin^−^CD34^+^CD38^−^Thy1^+^CD45RA^−^CD49f^+^Rhodamine123low^73^ (**Supplementary Table 15**), were sorted into 96-well plates. Wells containing single live cells were reprogrammed with the CytoTune-iPS 2.0 Sendai Reprogramming Kit (cat# A16517). Successfully reprogrammed colonies were identified by alkaline phosphatase staining (cat# SK-5100), transferred to 6-well plates, and maintained in Essential 8 medium with daily medium changes. Generated iPSCs were shown to express pluripotency markers, are transgene free, and differentiate into three germ layers using an embryoid body hanging drop assay, as described previously^74^. DNA was extracted with the Quick-DNA Miniprep Plus Kit (Zymo Research, cat# D4069).

### Brain samples

76 primary age-related tauopathy (PART) brain samples were obtained from the prefrontal cortex (PFC) of donors in the PART Working Group (PWG) cohort (**Supplementary Table 10**). Fresh-frozen tissue was collected from the contributing centers detailed in a previous study^75^, and all tissue was used in accordance with institutional guidelines and regulations.

Neuropathological assessment included Braak neurofibrillary tangle staging^76^ and CERAD neuritic plaque scoring^77^. Inclusion required a Braak stage of I–IV and a CERAD score of 0. Individuals with motor neuron disease, parkinsonism, frontotemporal dementia or other neurodegenerative tauopathies were excluded, although aging-related tau astrogliopathy (ARTAG) was permitted. Genomic DNA was isolated from 20–40 mg of fresh-frozen PFC tissue using the MagMAX DNA Multi-Sample Ultra 2.0 Kit on a KingFisher Flex system (Thermo Fisher Scientific). Tissue was digested overnight at 65 °C in 480 µL of proteinase K mixture, followed by magnetic bead-based purification. DNA quality was assessed by NanoDrop spectrophotometry, with samples required to have a concentration above 50 ng/µL and a 260/280 ratio between 1.7 and 2.2. 28 postmortem frozen brain tissue samples from the temporal lobe adjacent to the hippocampus were obtained from the Neuropathology Brain Bank & Research CoRE (NPBB; RRID: SCR_027565) at the Icahn School of Medicine at Mount Sinai (**Supplementary Table 10**). Donors’ legal next-of-kin consented to brain donation in accordance with ethical guidelines and IRB approval at the Icahn School of Medicine at Mount Sinai. Diagnoses were made by the attending neuropathologist from review of clinical records together with external and microscopic examination of the brain, following a standardized NPBB workflow that records data points compatible with the National Alzheimer’s Coordinating Center (NACC) database. In this workflow, each donated brain is bisected sagittally in the fresh state, with one hemibrain parcellated and rapidly frozen. The other is immersion-fixed in 10% neutral buffered formalin (Leica Microsystems, cat# 3800600) at 4 °C for two to three weeks, transferred to 1× phosphate-buffered saline with 0.01% sodium azide (Sigma Aldrich, cat# S2002-100G), then sampled to produce formalin-fixed paraffin-embedded (FFPE) blocks. All cases underwent H&E staining, with or without Luxol fast blue counterstaining, to assess morphology. Chromogenic immunohistochemistry was performed on 5 µm FFPE sections using the Ventana Discovery Ultra platform (Roche Diagnostics) following the manufacturer’s protocols, with CC1 antigen retrieval and hematoxylin counterstaining. The stains and diagnostic staging criteria applied to each brain comprised the distribution of phosphorylated-tau neurofibrillary tangles by Braak criteria^76^ with p-tau IHC (AT8, 1:1,000; Roche Diagnostics), the density of beta-amyloid neuritic plaques by CERAD criteria with Bielschowsky silver staining, and the combined Alzheimer’s disease neuropathologic change (ADNC) ABC score^77,78^. Donors were classified as AD when intermediate or high ADNC was accompanied by a clinical cognitive status of dementia, and as MCI when clinical cognitive change was documented without meeting these criteria. NCI donors were controls selected for the absence of any documented cognitive change. Genomic DNA was extracted from up to 500 mg of frozen brain tissue using the Wizard Genomic DNA Purification Kit (Promega). For larger tissue inputs, reagent volumes were increased proportionally at each step to maximize yield. DNA was eluted in 0.1× IDTE buffer (IDT), prepared by diluting the stock TE solution 1:10 in nuclease-free ultrapure water (Thermo Fisher Scientific).

### Sorted and derived brain cell populations

Sorted microglia were isolated from the fresh dorsolateral prefrontal cortex (DLPFC) tissue of an 81-year-old female, with sample collection and processing described previously^79^.

The GABAergic neuron sample comprised medial ganglionic eminence (MGE)-derived neurons from the DLPFC of a 58-day-old male infant and is enriched for inhibitory neurons. The mixed glia sample was taken from the DLPFC of a 25-year-old female donor and contains all major non-neuronal cell types, including microglia, astrocytes, oligodendrocytes and oligodendrocyte precursor cells (OPC), with oligodendrocytes constituting the majority. Nuclei for both populations were isolated by fluorescence-activated nuclei sorting (FANS). Further details of sample collection and processing have been described previously^80^.

iPSC-derived microglia sample is day 45 microglia obtained from the 2607-2C3-VPR-iMGL line, a male control CRISPR activation line. At harvest the cells had not undergone a recent media change and showed abnormal morphology, indicating suboptimal culture condition. Genomic DNA was isolated using the Wizard kit (Promega).

iPSC-derived neuron sample is the 553-3 hiPSC line established from fibroblasts of a healthy male donor by SV-KOSM reprogramming, modified by lentiviral integration of a Cas9 expression cassette (V2-Puro), and subcloned to give the daughter line 553-3C4. For neuronal differentiation, 553-3C4 cells were transduced with separate lentiviruses carrying NGN2 and M2rtTA expression cassettes. Genomic DNA was isolated using the Wizard kit (Promega).

### Colon samples

Early-onset colorectal cancer (EO-CRC) specimens comprised 17 tumor and adjacent normal tissue pairs together with 6 additional tumor samples without matched normal tissue, all provided by the Biorepository and Pathology Core at the Icahn School of Medicine at Mount Sinai (**Supplementary Table 12**). Late-onset colorectal cancer (LO-CRC) specimens comprised 7 tumor and adjacent normal tissue pairs obtained from surgical resection specimens at the Herbert Irving Comprehensive Cancer Center, Columbia University Irving Medical Center, under IRB protocol AAAT8778, together with 3 tumor and adjacent normal tissue pairs from the Mount Sinai Biorepository and Pathology Core that were ONT-sequenced in our previous study^24^ (**Supplementary Table 12**). Collection of the Mount Sinai specimens and the associated IRB approval have been described previously^24^. Each specimen was classified at collection as either tumor core or adjacent normal, and for adjacent normal specimens the distance from the tumor margin was recorded, ranging from 1 to 9 cm. All specimens were flash-frozen in liquid nitrogen and stored at −80 °C until genomic DNA extraction with the Wizard Genomic DNA Purification Kit (Promega).

### Colon fibroblast cell line

Primary colonic stromal cells were isolated from ten rectal pinch biopsies of histologically healthy mucosa from a 28-year-old male donor with Crohn’s disease (15 years since onset, maintained on risankizumab). Biopsies were rinsed in ice-cold PBS and incubated in PBS with 10 mM EDTA for 20 min at 37 °C to release epithelial crypts, which were shaken off and discarded. Remaining fragments were washed repeatedly until the supernatant was free of epithelium, then dissociated in HBSS containing Liberase TL and DNase I in three 20-min incubations at 37 °C, with pipetting every 10 min. Supernatant was collected into medium containing 10% FCS after each incubation to quench the Liberase. Released cells were filtered through a 70 µm strainer, pelleted at 1,000 g for 4 min at 4 °C, and seeded in stromal medium (αMEM with 5–10% PLT Gold, penicillin–streptomycin and Normocin). Cultures were expanded in the same medium and passaged with TrypLE. Cells were cryopreserved in Cell Recovery Medium (Gibco) and stored at −80 °C. Genomic DNA was extracted using the Wizard Genomic DNA Purification Kit (Promega).

### PacBio long read sequencing

Genomic DNA was sheared to approximately 20 kb using a g-TUBE at 4,200 rpm. Following AMPure PB bead cleanup, SMRTbell libraries were prepared with the SMRTbell Prep Kit 3.0 with several modifications to the standard protocol. End repair was carried out for 1 h, ligation for 3 h to overnight, and bead elution was extended to 45 min at 37 °C to improve recovery.

AMPure PB beads were used at 0.5× for the final cleanup in place of the dilution step specified by the manufacturer, which improved library retention. Eluted material was combined with a Tween-20 rinse of the well using the manufacturer-supplied solution, and all samples were cleaned up with AMPure PB beads. Libraries were typically prepared from pools of two samples. Size selection was performed on a BluePippin instrument with the S1 external marker, giving a sharp cutoff that excluded fragments shorter than 8 kb. PacBio sequencing data collection and processing were carried out at HistoGenetics. Whole-genome sequencing was performed on the PacBio Revio platform according to the manufacturer’s standard workflow. SMRTbell libraries were annealed with sequencing primers and bound to DNA polymerase using the Revio Polymerase Kit (PacBio, cat# 102-739-100) or Revio SPRQ Polymerase Kit (PacBio, cat# 103-496-900). After purification, polymerase-bound SMRTbell complexes were quantified on a Qubit fluorometer (Thermo Fisher Scientific) and diluted to a final concentration of 300 pM. Libraries were combined with loading buffer and a sequencing control, then loaded onto SMRT Cells 25M. Data were acquired over 30 h runs per SMRT Cell under standard conditions with adaptive loading enabled.

### Nanopore long read sequencing

R10 libraries were prepared with the Native Barcoding Kit 24 V14 (SQK-NBD114.24) and loaded onto PromethION flow cells. The manufacturer’s instructions were followed except as noted. DNA repair and end preparation were extended to 30 min at 20 °C, with no corresponding increase at 65 °C, since 5 min is sufficient to denature the enzyme. Barcoding and adapter ligation were each carried out for 1 h. Bead elution was extended to 45 min to improve yield, and AMPure PB beads were used at 0.37× for the final cleanup, a stricter cutoff that reduced the proportion of short fragments. Flow cells were run for up to 96 h, with washing and reloading on each day of the run using the Flow Cell Wash Kit XL (EXP-WSH004-XL) to maximize output.

### DNA methylation array

Plates of diluted DNA samples were prepared by aliquoting 600 ng of each sample into a PCR plate in a total volume of 40 µl of 0.1× Tris-EDTA buffer. DNA methylation was profiled using the Illumina Infinium Methylation Screening Array (cat# 20112611) on Tecan Freedom EVO liquid handlers. Genomic DNA was bisulfite-converted with the Qiagen EpiTect Bisulfite kit (cat# 59104), loading 40 µl of DNA at no more than 12.5 ng/µl. Converted DNA was denatured, neutralized, and isothermally amplified overnight, then fragmented by end-point fragmentation and precipitated with isopropanol. The fragmented DNA was resuspended in hybridization buffer and hybridized to the array overnight in an Illumina Hybridization Oven. Unhybridized and non-specifically hybridized DNA was then washed away, and single-base extension with labelled nucleotides was performed on the array XStain. Arrays were imaged on an Illumina iScan. Raw methylation array data normalization was performed by minfi v1.56.0^81^.

### Read alignment and quality control

Long reads were aligned to the GRCh38 human reference genome. PacBio reads were mapped (https://github.com/PacificBiosciences/pbmm2) with pbmm2 v1.13.0 and ONT reads with Dorado v1.0.2 (https://github.com/nanoporetech/dorado). Alignments with a mapping quality below 30 were discarded.

### Modification calling and pass number bias correction for PacBio sequencing data

Modified base calling of PacBio data was first performed with jasmine v2.4.0 (https://github.com/PacificBiosciences/jasmine). In the resulting data, the modification probabilities encoded in the ML tag (0–255) depended strongly on read pass number, with low-pass reads yielding systematically lower and more ambiguous values than high-pass reads (**Supplementary Fig. 2A**). When reads were stratified by pass number, the genome-wide methylation level estimated from each stratum rose with pass number and stabilized from 15 passes onward (**Supplementary Fig. 2A**). We therefore took the genome-wide methylation level of reads with ≥15 passes, called at a fixed probability threshold of 180, as the reference (**Supplementary Fig. 2C**). For reads with 3 to 14 passes, the threshold was calibrated separately at each pass number so that the resulting genome-wide methylation level matched this reference. Both the reference and the calibrated thresholds were derived independently for each sample. Where the calibrated threshold reached the limit of its adjustable range without matching the reference, reads of that pass number were discarded.

Modified bases were then called a second time, applying the calibrated threshold for each pass number from 3 to 14 and the fixed threshold of 180 for reads with ≥15 passes. Only reads with at least three passes were retained (**Supplementary Fig. 2C**). For this step, modification calls were decoded directly from the MM and ML SAM tags of the jasmine output: for each read, the MM tag was used to locate modified positions relative to the canonical bases, accounting for read orientation, and the corresponding ML values were converted from their 8-bit encoding into modification probabilities. At each cytosine, the 5mCpG and canonical probabilities were compared to assign a modification state, and read positions were mapped to reference coordinates using the alignment. Bases whose ML value fell between the applicable threshold and its complement were treated as uncertain and excluded from downstream analysis, so that only confident methylated and unmethylated calls were retained.

### Modification calling for ONT sequencing data

Modified base calling of ONT data was performed using Dorado v1.0.2 with the 5mC_5hmC or (https://github.com/nanoporetech/dorado; 5mCG_5hmCG models **Supplementary Tables 1–2**). Read-level base modification information was then extracted with modkit v0.3.1 (https://github.com/nanoporetech/modkit) using the *modkit extract* function. A modification likelihood threshold of 180 was used to assign the modified state of a 5mCpG and 5hmCpG base and 225 for a 5mCpH base, the more stringent cutoff applied because 5mCpH calls carry a higher false discovery rate^82^.

### Curation of cell-type methylation reference panels across human tissues

Cell-type methylation reference panels were curated for three tissues, blood, colon, and brain. Principal component analysis (PCA) based on genome-wide methylation profiles within each panel was performed to confirm the cell types were well separated, supporting their use as classification references.

The blood and colon panels were built from high-depth bisulfite sequencing (BS-seq) of sorted cell populations. For blood, cell-type-level methylation profiles were drawn from Loyfer et al.^2^ and BLUEPRINT^19^ to assemble six cell types: B Memory, B Naive, myeloid, NK, T Ag-exp (antigen-experienced T cells, encompassing T effector and T memory cells), and T Naive. For colon, we used cell-type-level profiles from Loyfer et al.^2^ to assemble three cell types: Epi-Endo, fibroblast, and macrophage. Epithelial and endocrine cells were combined into a single Epi-Endo class because their genome-wide methylation profiles clustered together by PCA. The brain panel was built from single-cell methylation sequencing (snmC-seq) data from Tian et al.^25^, from which we assembled six cell types: excitatory neurons, inhibitory neurons, astrocytes, microglia, oligodendrocytes, and oligodendrocyte precursor cells (OPC). For the brain panel, single cells were aggregated into cell-type-level (pseudobulk) methylation profiles. Non-neuronal cells were merged directly within each cell type. Neuronal cells were first merged within each neuronal subtype, and because PCA showed that subtypes clustered by class, these were then merged into single excitatory neurons and inhibitory neuron profiles. For every tissue panel, only CpGs covered by at least 50 reads in all cell types were retained.

### Development of COMET

#### Binarization of methylation calls

From each bulk long-read BAM file, only primary alignments that mapped to autosomes and passed read alignment quality control were retained for cell type assignment. CpG methylation calls are stored per read in the MM and ML tags. A call was kept only if the read sequence at that position was CG and the base aligned to the reference as a match, and it was then mapped to its reference coordinate. Methylation states were called using a pass-number-calibrated cutoff for PacBio data (see “Modification calling and pass number bias correction for PacBio sequencing data” in **Methods**) and a fixed cutoff for ONT data (see “Modification calling for ONT sequencing data” in **Methods**). Each retained call was then binarized, with calls above the cutoff coded as methylated (1) and those below as unmethylated (0).

#### Read-level likelihood model

For a read *r*, let {1,…,*N*_*r*_} be its informative CpGs, those covered by the reference panel whose call passed binarization under the modification calling cutoff. Let 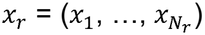, with *x*_*i*_ ∈ {0,1}, be its observed binary methylation states. For each candidate cell type *k* ∈ {1,…,K}, let β_*k,i*_ be the reference methylation level of cell type *k* at CpG *i*. A read was scored only if *N*_*r*_ ≥ 5. If a read originates from cell type *k*, the methylation state at each CpG is modelled as an independent Bernoulli draw with success probability β_*k,i*_. The log-likelihood of the read under cell type *k* is

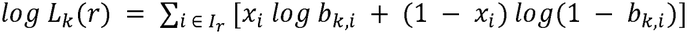

where

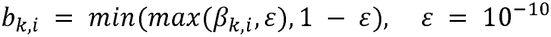

Clipping *β*_*k,i*_ at *ε* prevents a single CpG that is fully methylated or fully unmethylated in the reference from driving the likelihood to −∞. Such a CpG still contributes a large penalty when the read disagrees with it, so CpGs that are invariant within one cell type and discordant with the read weigh strongly against that cell type.

#### Posterior probability and read assignment

With a prior *π*_*k*_ over cell types, the posterior probability that read *r* originates from cell type *k* is

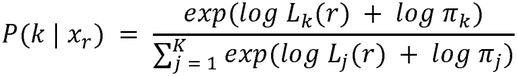

This is the softmax of the log-posterior scores and is computed in log space for numerical stability. In this study we used a uniform prior, 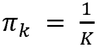. The prior therefore cancels and the posterior reduces to the normalised likelihood. A uniform prior means that read-level assignment does not depend on any assumed tissue composition, so the cell-type proportions that follow are estimated from the data rather than set by a prior. Let *k*^*^ = *argmax*_*k*_ *P*(*k* | *x*_*r*_), and let *P*_(1)_ ≥ *P*_(2)_ be the largest and second-largest posteriors. Read *r* was assigned to *k*^*^ only if both of the following conditions held: (i) *P*_(1)_ ≥ *θ*, where *θ* is the tissue-specific posterior probability cutoff selected from the simulation analysis (see “Simulation-based evaluation of cell type assignment by COMET” in **Methods**), with *θ* = 0.65 for blood and brain and *θ* = 0.5 for colon, and (ii) *P*_(1)_ ≥ 2 × *P*_(2)_. Reads that failed either condition were labelled unassigned. For each scored read, COMET reports the read identifier *N*_*r*_, the assigned label and the full posterior vector (*P*(1 | *x*_*r*_), …, *P*(*K* | *x*_*r*_)). The absolute cutoff removes reads whose CpGs carry little information.

The ratio rule removes reads that cannot be distinguished from a closely related cell type even when the top posterior is high. Assignment uses only each read’s methylation pattern and the reference panel, never sample age, diagnosis, genotype or group label, so it is blind to sample condition and any systematic difference it produced would act equally across groups.

#### Lineage-level assignment

When the reference panel contains closely related cell types, many unassigned reads have posterior mass that is split almost entirely within one lineage. These reads were assigned to a lineage-level group *G* (a set of two or more related cell types) when all of the following conditions held: (i) the cell types in *G* are the top-| *G* | ranked posteriors of the read, (ii) ∑_*k ∈ G*_ *P*(*k* | *x*_*r*_) ≥ *θ*, where *θ* is the tissue-specific posterior probability cutoff selected from the simulation analysis (see “Simulation-based evaluation of cell type assignment by COMET” in **Methods**), and (iii) *min*_*k ∈ G*_ *P*(*k* | *x*_*r*_) > *max*_*j ∉ G*_ *P*(*j* | *x*_*r*_) + *δ*, with *δ* = 0.10. Lineage-level assignment was applied only in the brain real-sample validation analysis, for the neuronal lineage with *G* comprising excitatory and inhibitory neurons, and for the macroglia lineage with *G* comprising astrocytes, oligodendrocytes, and OPC, using *δ* = 0.65.

#### Cell-type proportion and cell-type-resolved methylation quantifications

<TC-F-BlockquoteIndent>For each cell type, the identifiers of its assigned reads were used to extract the corresponding alignments from the original bulk BAM file, yielding one cell-type-specific BAM per sample. Cell-type composition was then estimated from the number of reads assigned to each cell type. For cell type *k*, the cell type proportion is 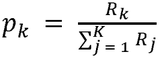, where *R*_*k*_ is the number of reads assigned to cell type *k* and the sum runs over all *K* cell types in the reference panel. Unassigned reads were excluded from the denominator. Genome-wide methylation was computed from all autosomal CpG calls. For cell type *k*, the methylation level *ml*_*k*_ was obtained by pooling binary calls across reads 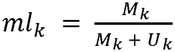, where *M*_*k*_ and *U*_*k*_ are the numbers of methylated and unmethylated calls among reads passing the modification calling cutoff. To exclude the possibility that a sample’s estimated composition and its cell-type methylation are two views of the same signal rather than independent measurements, assignment draws only on the reference panel sites while methylation quantification uses genome-wide sites, keeping the two separable.

#### Cell-type-resolved methylation output for differential methylation analysis

Position-level methylation calls were generated from the read set of each cell type, using the pass-number-aware calibration pipeline integrated in COMET for PacBio data (see “Modification calling and pass number bias correction for PacBio sequencing data” in **Methods**) and modkit for ONT data (see “Modification calling for ONT sequencing data” in **Methods**). These calls are written in standard BED format accepted by differential methylation tools and support two complementary comparisons. Comparing different cell types within one sample identifies cell-type-specific differential methylation internal to that sample, while comparing the same cell type across samples or groups identifies cell-type-resolved differential methylation between conditions.

### Simulation-based evaluation of cell type assignment by COMET

To measure how accurately COMET assigns individual reads to their cell of origin, we generated simulated long reads of known cell-type identity for each tissue. Autosomal reads were simulated genome-wide at approximately 2× coverage, chosen to match the median depth of the shallow PacBio-sequenced cohort (1.88×), with read lengths drawn from a normal distribution *N*(*µ* = 10,000 bp, *σ* = 2,500 bp). For each cell type in a tissue’s reference panel, the full set of simulated reads was treated as originating from that cell type. Read coordinates were intersected with the reference panel to identify the CpGs each read covers, and the methylation state of every covered CpG was drawn as a Bernoulli variable using that cell type’s reference methylation level at that site as the success probability. Repeating this for every cell type yielded a labelled read set in which the true cell type of each read is known by construction.

We then applied COMET to these reads across probability thresholds from 0.50 to 0.95 in increments of 0.05. At each threshold, a read was assigned to a cell type when its highest probability exceeded the threshold and was otherwise left unassigned. Accuracy was evaluated per cell type on the assigned reads, with the unassigned fraction recorded separately. For cell type *i*, true positives (TP) are reads simulated as *i* and assigned to *i*, false positives (FP) are reads simulated as another cell type but assigned to *i*, false negatives (FN) are reads simulated as *i* but assigned elsewhere, and true negatives (TN) are reads neither simulated as nor assigned to *i*. From these we computed 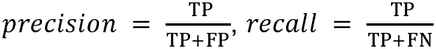, and their harmonic mean, 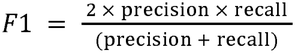, for call type *i*.

Raising the threshold improves F1 but reduces the fraction of reads assigned. Balancing these, we required each tissue’s threshold to satisfy two criteria simultaneously: every cell type reaching an F1 score above 0.84, and more than 40% of simulated reads from every cell type receiving an assignment. For downstream analysis of real samples, we took the lowest threshold meeting both, giving 0.50 for colon and 0.65 for blood and brain.

### Simulation-based evaluation of cell-type quantifications by COMET

We built an *in silico* benchmark to test how accurately COMET recovers cell-type proportions and cell-type-resolved methylation levels. The benchmark uses computationally constructed read sets in which both the cellular composition and the methylation level of each cell type are defined by construction rather than drawn from real tissue, allowing us to span a far wider range of compositions and methylation levels than real samples provide, each with an exactly known value.

For each tissue, we generated 5,000 random cell-type mixtures, covering six cell types for blood, three for colon and six for brain. Cell-type proportions were drawn from a uniform Dirichlet distribution, with *p*_*i*_ denoting the proportion of cell type *i*. To ensure that all cell types were represented at appreciable proportions, mixtures in which any cell type fell below a predefined minimum proportion were discarded and redrawn. Two sets of 5,000 mixtures were generated using minimum cell-type proportions of 1% and 5%, respectively. For each cell type, a pool of 100 million reads was simulated and reused across mixtures. For each mixture, a total read count *N* was randomly drawn between 1 and 10 million. Within each mixture, *N* × *p*_*i*_ reads representing cell type *i* were sampled without replacement from the corresponding read pool.

The benchmark was designed to test whether COMET recovers cell-type methylation levels that differ from the reference panel used for assignment. If simulated reads carried exactly the reference methylation profile, recovery would be trivial, so we generated read sets whose methylation levels deviate from the reference while assignment continued to use the unchanged reference panel. For each cell type, we took the mean *µ* and variance *σ*^2^ of methylation averaged over non-overlapping blocks of 100 consecutive CpGs in the reference methylome, using only sites with at least 50× coverage, so that *σ*^2^ reflects the regional methylation heterogeneity seen in real data. Each read from a cell type was modelled as carrying its own methylation rate *r*∼*N*(*µ, σ*^2^), with each of its CpGs methylated with probability *r*.

This gives read sets whose overall methylation varies from mixture to mixture, including cases where a cell type is substantially less methylated than its reference profile. The rates at which reads of each true cell type are assigned to each cell-type class were taken from COMET run on real reads of known origin, rather than re-derived from the simulated reads. Simulated assignment therefore carries the site-specific behavior of the real classifier. Total methylated and covered CpG counts per read set were drawn directly from the mean and variance implied by the model above, rather than by simulating individual CpG sites.

COMET was applied to each simulated read set. Reads whose highest cell-type posterior probability fell below the tissue threshold, 0.50 for colon and 0.65 for blood and brain, according to simulation-based evaluation of cell type assignment; only reads above the threshold were retained for quantification of cell-type proportions and cell-type-resolved methylation levels. Cell-type proportions and cell-type-resolved methylation levels constructed from the simulation and estimated from the COMET assignments, were compared by Pearson correlation across all simulated mixtures.

### Downsampling analysis

For each tissue, the sample with the highest sequencing depth within the shallow-coverage cohort was selected for downsampling, since this cohort is used for cell-type proportion and methylation quantification and the analysis was designed to test whether depths in this range are sufficient. Three target depths were defined: the lowest depth in the corresponding tissue cohort and two intermediate depths between the lowest and original depths. At each target depth, reads were randomly downsampled 2,000 times with samtools v1.22^83^, and cell-type proportions and cell-type-resolved methylation levels were quantified and compared with those at full depth.

### Evaluation of COMET using real samples

Cell type assignment was evaluated using samples whose cellular composition is known to differ from that of the corresponding bulk tissue. In each case the comparison samples are enriched for, though not necessarily pure in, the expected cell type. For blood, PBMC samples, which are depleted of myeloid cells, were compared against the whole-blood samples in our cohort, using PacBio data throughout. For colon, a fibroblast cell line was compared against the colon samples in our cohort, using both PacBio and ONT data. For brain, ONT data were used to compare six deep-sequenced bulk brain samples against five populations each dominated by a single cell type: sorted microglia, iPSC-derived microglia, GABAergic neurons, iPSC-derived neurons, and mixed glia.

To test whether including lower-specificity CpGs preserves assignment quality while recovering more reads, we compared assignment using all reference-panel CpGs against specificity-ranked subsets. For a tissue with *n* cell types, let *m*_*i*_ denote the methylation level of cell type *i* at a given CpG. The mean methylation level of that CpG across all cell types is: 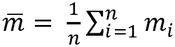 The cell-type specificity *s* of a CpG was defined as the population standard deviation of its reference methylation levels across the *n* cell types, with higher *s* indicating greater divergence among cell types: 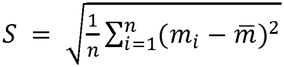. CpGs were ranked separately for blood, colon and brain, and nested panels were built from the top 20%, 40%, 60% and 80% of scores, retaining ties at each cutoff. Each subset was applied to one sample per tissue dominated by a single known cell type, namely PBMCs for blood, a colon fibroblast cell line for colon and sorted microglia for brain. For each subset we recorded the number of assigned reads, the estimated cell-type proportions and the distribution of assignment probabilities, and compared these against the values obtained with the complete reference panel, which is what COMET uses.

### CyTOF profiling

Mass cytometry was performed on 10 whole blood samples from donors without known disease, collected independently of the 140-sample blood cohort and also profiled by PacBio sequencing. Immune profiling used the Maxpar Direct Immune Profiling Assay (Standard BioTools, cat# 201325), a 30-marker core panel expanded to 42 markers with 12 spiked-in add-on antibodies (**Supplementary Table 15**). Add-on antibodies were combined undiluted in cell staining buffer (Standard BioTools, cat# 201068), centrifuged (15,000 × g, 15 min, 4 °C) to remove aggregates, aliquoted, and stored at −80 °C. 270 µL fresh blood (1–3 × 10□ cells) was added to a Maxpar Direct assay tube (Standard BioTools, cat# S10310) with the add-on cocktail, incubated 30 min at room temperature, stabilized with 420 µL Prot1 for 10 min, and stored at −80 °C. Samples were thawed, lysed in Thaw-Lyse Buffer (Smart Tube, cat# 501351696), fixed in 1 mL 4.4% paraformaldehyde, labeled with 125 nM iridium-193 (Standard BioTools, cat# S00093), and resuspended at 1 × 10□ cells/mL in Cell Acquisition Solution (Standard BioTools, cat# 201240) containing EQ normalization beads (1:20; Standard BioTools, cat# 201078). Acquisition used a Helios mass cytometer with a wide-bore injector at <400 events/s.

FCS files were read with flowio v1.4.0 and ion counts transformed as arcsinh(x/5). Intact single live leukocytes were retained by sequential gating on ^1^□^3^Ir DNA, event length, ^1^□□Ce (normalization beads), ^1^□□Pt (viability) and CD45, and all proportions are expressed as a percentage of CD45? events. Positivity thresholds were set once on a pooled subsample drawn equally from the 10 samples, placed at the valley of the smoothed pooled histogram, and applied unchanged to every sample; implausible thresholds were replaced by a fixed cutoff.

Differentiation markers were thresholded within their parent lineage. For comparison with COMET, these were collapsed into the six reference-panel cell types: B Naive (CD20^+^CD27^−^), B Memory (CD20^+^CD27^+^), T Naive (CD3^+^CD45RA^+^CCR7^+^), T Ag-exp (remaining CD3^+^ cells), NK, and myeloid (neutrophils, eosinophils, basophils, all monocyte subsets, mDC and pDC); remaining CD45^+^ events were assigned as other.

### Decomposition of genome-wide blood bulk methylation differences between age groups into cell type contributions

Six cell types were assigned and analyzed in blood aging. Within each sample, cell-type proportions were normalized to the total number of assigned reads. For each age group containing *n* samples, and for cell type *i*, the group-level methylation level and cell-type proportion are 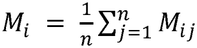 and 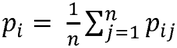, respectively. The group-level observed bulk methylation level, computed over all reads including those not assigned to a cell type, and the bulk level reconstructed from the six cell types, are 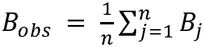 and 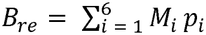, respectively.

Differences between the two age groups are taken as older minus younger. *ΔB*_*obs*_ and *ΔB*_*re*_ represent the observed and reconstructed methylation level difference between the two age groups, respectively. For cell type *i, ΔM*_*i*_ and *Δp*_*i*_ represent the differences in mean methylation level and cell-type proportion between the two age groups, respectively, and the 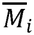 and 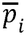 denote the averages of the two group means, with 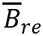 the average of the two reconstructed bulk levels.

The intrinsic contribution (*I*_i_), reflecting within-cell-type methylation change, and compositional contribution (*C*_i_), reflecting change in that cell type’s abundance, are: 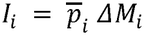 and 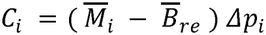, respectively. Two residual terms reconcile the measured bulk difference with this reconstruction. The covariance residual is the change between groups in the covariance across samples between a cell type’s abundance and its methylation level: 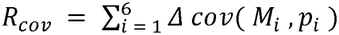, where 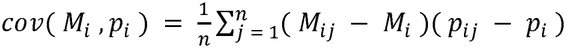; the measurement residual is the difference between the measured bulk level and the level reconstructed from assigned reads, reflecting unassigned reads and any residual weighting difference: *R*_2_ = *ΔB*_*obs*_ − *ΔB*_*re*_ − *R*_*cov*_.

Each component was classified by the direction in which it moved bulk methylation. Writing the twelve contributions and two residuals, the components lowering bulk methylation were summed to give the total decrease, against which each component’s share was taken, with components acting in the opposite direction with bulk methylation level decrease assigned a share of zero.

### Quantifying Between-cell-type and within-cell-type methylation differences at Horvath clock CpG regions

To characterize the relative contribution of within-cell-type age-associated methylation change and between-cell-type methylation heterogeneity at each Horvath clock^6^ CpG region (±500 bp around each Horvath clock CpG), we computed two absolute-difference-based summary metrics per region in two young and two old deep-sequenced blood samples. Both metrics are defined over whichever cell types and sample pairs are measurable at a given region, where a cell type was considered measurable in a sample if it was covered at a depth of at least 3× at that region. Only regions for which both metrics could be computed were retained.

Mean absolute within-cell-type methylation difference between young and old samples, averaged first across all valid young–old pairings within each cell type and then across all contributing cell types. Cell types with no valid pairing at the region are excluded from this region. For each target region, let *M*_*i*_ denote the regional methylation level of cell type *i* in a sample *s, Y* = {*s*_*Y*1_,*s*_*Y*2_} and *o* = {*s*_*o*1_,*s*_*o*2_} denote the sets of young and old samples, respectively. For each cell type *i*, let *P*_*i*_ be the set of young–old sample pairs for which both samples are measurable at region in cell type *i*. Let *C* be the set of cell types with at least one such pair.

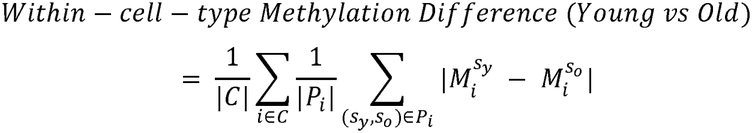

Mean absolute between-cell-type methylation difference within samples, computed across all available cell-type pairs in each sample and then averaged across samples. Samples with fewer than two measurable cell types are excluded for this region. For each target region, let *M*_*i*_ denote the regional methylation level of cell type *i* in a sample *s* regardless of young or old. For each sample *s*, let *Q*_*s*_ be the set of measurable cell type pairs in that sample at the region. Let *D* be the set of samples with at least two measurable cell types.

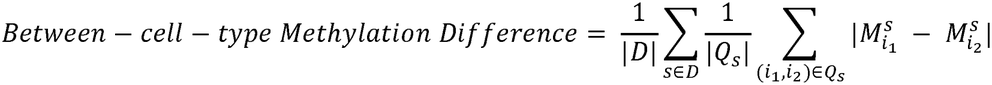

### Identification of AD resilience candidates by chart review for blood samples

Among participants with blood samples, APOE4 homozygotes (APOE44) aged 75 years or older at enrollment were selected to determine AD resilience status, as this age exceeds both the mean age at symptom onset (65.6 years) and the mean age at dementia diagnosis (73.6 years) reported for APOE4 homozygotes based on prior clinicopathological work^84^. Chart review of the linked EHR was then performed for these candidates, and resilience was defined by two criteria. First, no diagnosis of a neurodegenerative disease was recorded and no cognitive impairment was documented within the 5 years following sample collection, as determined by ICD-9/ICD-10 billing codes, documented prescription of medications used to treat dementia or upon independent chart review, ensuring that resilient individuals had neither latent pathology at the time of sampling nor subclinical deficits that emerged shortly afterwards. Second, plasma pTau217 and NfL concentrations, retrieved from the same records and profiled by Quanterix Simoa, were non-elevated, defined as pTau217 below 0.66 pg/mL and NfL below 22 pg/mL^85^.

### Identification of DMRs

For each cell type, modifications were called from the reads assigned to it, giving a separate set of methylation calls per cell type as input to DMR analysis. DMRs were identified with metilene v0.2.9^86^ in *de novo* segmentation mode, requiring at least 5 CpGs per region and no more than 300 bp between consecutive CpGs. DMRs were retained only where the region was covered at a depth of at least 3× in every sample used for the differential comparison.

Condition-specific and cell-type-resolved DMRs were then called by applying a methylation difference threshold to the retained regions, with the threshold and comparison defined per tissue. For blood, group-specific and cell-type-resolved DMRs were those where the target group differed by at least 0.3 in methylation level from the remaining samples. For colon, tctDMRs were those where one tumor cell type differed by at least 0.3 from every other tumor cell type and from all normal cell types. For brain, group-specific and cell-type-resolved 5mCpG and 5hmCpG DMRs were called at a threshold of 0.2, since the anti-correlation between the two marks can dilute the apparent difference in either one.

### Repeat annotation

CpGs that were resolved to a cell type but lie outside the reference panel were annotated with repeat elements from RepeatMasker (https://www.repeatmasker.org/). Modification levels for each transposable element category were quantified by pooling calls across all elements of that category. Full-length L1 elements were defined as L1 annotations of at least 5,500 bp and full-length HERV elements as HERV annotations of at least 7,000 bp.

### Integration with epigenomic and transcriptomic datasets

For blood, group-specific and myeloid-resolved DMRs were annotated as candidate promoters or enhancers using blood cell-type histone modification data from ROADMAP^20^. For the individual DMR shown as an example, regional histone mark signal of blood cell types was examined in both ROADMAP and BLUEPRINT^19^ data, and overlap with ENCODE candidate cis-regulatory elements (cCREs) of myeloid cells was assessed using the SCREEN annotations^87^. Candidate regulatory elements were linked to target genes using promoter-capture Hi-C (PCHi-C) data from Javierre et al.^88^, where interactions with a CHiCAGO score^89^ of at least 5 were retained as significant promoter-enhancer contacts.

For brain, group-specific and cell-type-resolved DMRs were annotated with candidate gene promoters, defined as the interval spanning 1,500 bp upstream to 500 bp downstream of transcription start site (TSS), and with gene bodies (**Supplementary Table 11**), based on GENCODE v45^90^. Switching and non-switching DMRs were further annotated as regulatory elements with echoannot v0.99.11^91^, using brain cell-type histone modification and chromatin accessibility data from Nott et al.^92^. Cell-type-resolved expression of *ANKH* was assessed using single-nucleus and single-cell RNA-seq data from seven public human cortex studies of AD and control donors^93-99^, retrieved from the CZ CELLxGENE Discover Census^100^ with cellxgene-census v1.18.0 and tiledbsoma v2.3.0, yielding 5,159,226 cells from 563 donors with Cell Ontology labels collapsed into the same six cell classes used in our brain reference panel and neurons additionally resolved into Allen Institute human cortical taxonomy subclasses. For each donor, counts were summed and expressed as counts per million. log_2_(CPM + 1) was modelled by ordinary least squares with diagnosis as the predictor of interest and study, region and sex as covariates, using donor-clustered robust standard errors, with Benjamini-Hochberg correction applied across cell types and significance assessed at FDR < 0.05.

For colon, tctDMRs were annotated with chromatin states from the ROADMAP 15-state ChromHMM^101^ segmentation of colonic mucosa. Gene-centric and CpG context annotations were based on GENCODE v45^90^, with the candidate promoter defined as the interval spanning 1,500 bp upstream to 500 bp downstream of the TSS. Genes identified by our earlier single-cell RNA-seq analysis^59^ as highly expressed in an epithelial cell state enriched in EO-CRC relative to LO-CRC were examined in the deep-sequenced colon samples, to test whether Epi-Endo-resolved methylation at these genes was reduced in EO-CRC but not in LO-CRC. Repeat element categories were annotated with RepeatMasker, and full-length L1 and HERV elements were defined by the length thresholds given above. L1HS and L1PA2 were classified as young L1s and all remaining L1 subfamilies as old, and HERVK was classified as young and all remaining HERV subfamilies as old. For the individual loci shown as examples, regional histone mark signal at the candidate promoter region of *MAL* was examined in the same ROADMAP colonic mucosa data, and the L1PA2 5′ UTR was identified by aligning the element against the L1PA2 consensus sequence.

### Evaluation of CRC screening marker genes at cell-type resolution

Screening marker genes were taken from the QIAseq Targeted Methyl Human Colorectal Cancer Panel (GeneGlobe ID MHS-002Z, QIAGEN), which comprises 65 genes reported to be hypermethylated in CRC (**Supplementary Table 14**). These include hypermethylation marker genes used in commercial stool-based CRC detection assays (Cologuard, ColoWell and Colocomf) and blood-based assays (Epi proColon and COLVERA).

For each gene, the candidate promoter region was defined as the interval spanning 1,500 bp upstream to 500 bp downstream of the TSS. Bulk and cell-type-resolved methylation levels were calculated across this candidate promoter region for each tumor and matched adjacent normal sample. Genes were classified as hypermethylated when bulk promoter methylation was higher in tumor than in matched normal tissue. For promoter regions with sufficient read coverage (≥3×) in all three colon cell types in both tumor and matched normal, the contribution of each cell type to the bulk methylation difference was decomposed as described below.

Reads were assigned to three cell types in colon analysis. *B*_*obs*_ denotes the regional bulk methylation level measured over all reads, including those not assigned to a cell type. At a given region and in a sample, the proportion of cell type *i* was taken as the fraction of cell-type-assigned reads in that region that were assigned to cell type *i*. Let the cell type proportion and cell-type resolved methylation level for cell type *i* be *p*_*i*_ and *M*_*i*_, respectively, and the regional bulk methylation level over assigned reads is 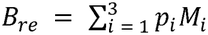. Differences between the two samples are taken as tumor minus normal. For cell type *i, ΔM*_*i*_ and *Δp*_*i*_ are the differences in regional methylation level and proportion between the two samples, and 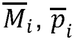, and 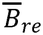 are the corresponding averages across the pair. The intrinsic contribution *I*_*i*_, reflecting change in that cell type’s own methylation, and the compositional contribution *C*_*i*_, reflecting change in its abundance, are 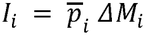 and 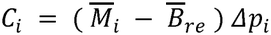. A single residual reconciles the measured bulk difference with this reconstruction, the difference between the regional bulk level measured over all reads and the level reconstructed from the assigned reads, reflecting reads not assigned to any cell type: *R* = *ΔB*_*obs*_ − *ΔB*_*re*_. No covariance residual arises because the decomposition is applied within each matched pair rather than to group means. Writing the six contributions and the residual, each component was classified at every region by whether it moved bulk methylation in the same direction as the observed regional change.

### Statistics

Pearson’s *r* was used for associations between continuous variables and Spearman’s ρ where variables were bounded or ordinal. Partial correlations were used to adjust by correlating residuals after regressing out the covariate. Group differences in methylation level and cell-type proportion were tested with linear mixed-effects models including group as the fixed effect of interest and age, sex and relevant clinical traits as covariates. Within each family of tests, *P* values were adjusted by the Benjamini–Hochberg procedure and significance assessed at adjusted *P* (*P*_adj_) < 0.05.

## Code Availability

The COMET software, together with documentation and a worked example, is available on GitHub (https://github.com/fanglab/COMET).

## Data Availability

Long-read methylation calls (including cell-type-resolved calls) and methylation array data generated in this study will be released on Zenodo and GEO upon publication. Raw sequencing reads and participant-level data from human donors will be available through controlled access, subject to donor consent and institutional approvals; requests can be sent to the corresponding author. Cell-type proportions and cell-type-resolved methylation levels for all samples used in this study are provided in **Supplementary Tables 3–5**. The curated reference panels are available on Zenodo at https://zenodo.org/records/24064938. Whole-genome BS-seq methylation calls for sorted blood and colon cell populations from Loyfer et al.^2^ are available in GEO under accession number GSE186458. BS-seq methylation calls and ChIP-seq histone modification calls for sorted blood cell populations from BLUEPRINT^19^ are available at https://ftp.ebi.ac.uk/pub/databases/blueprint/. Single-cell methylation calls for brain cell types from Tian et al.^25^ are available at https://cellxgene.cziscience.com/collections/fdebfda9-bb9a-4b4b-97e5-651097ea07b0.

ChIP-seq histone modification calls from ROADMAP^20^ are available at https://egg2.wustl.edu/roadmap/data/. Processed PCHi-C data from Javierre et al.^88^ are available through the Open Science Framework at https://osf.io/u8tzp. ENCODE candidate cis-regulatory element annotations are available at https://screen.wenglab.org/. Processed single-cell brain epigenomic data (H3K27ac, H3K4me3, and ATAC-seq) from Nott et al.^92^ are available on GitHub at https://github.com/nottalexi/brain-cell-type-peak-files. The brain single-nucleus and single-cell RNA-seq datasets reanalyzed were obtained from the CZ CELLxGENE Discover Census^100^ (https://cellxgene.cziscience.com).

