## Supplementary Figures for "Cell-type-resolved DNA methylation analysis using bulk long-read sequencing across human tissues"

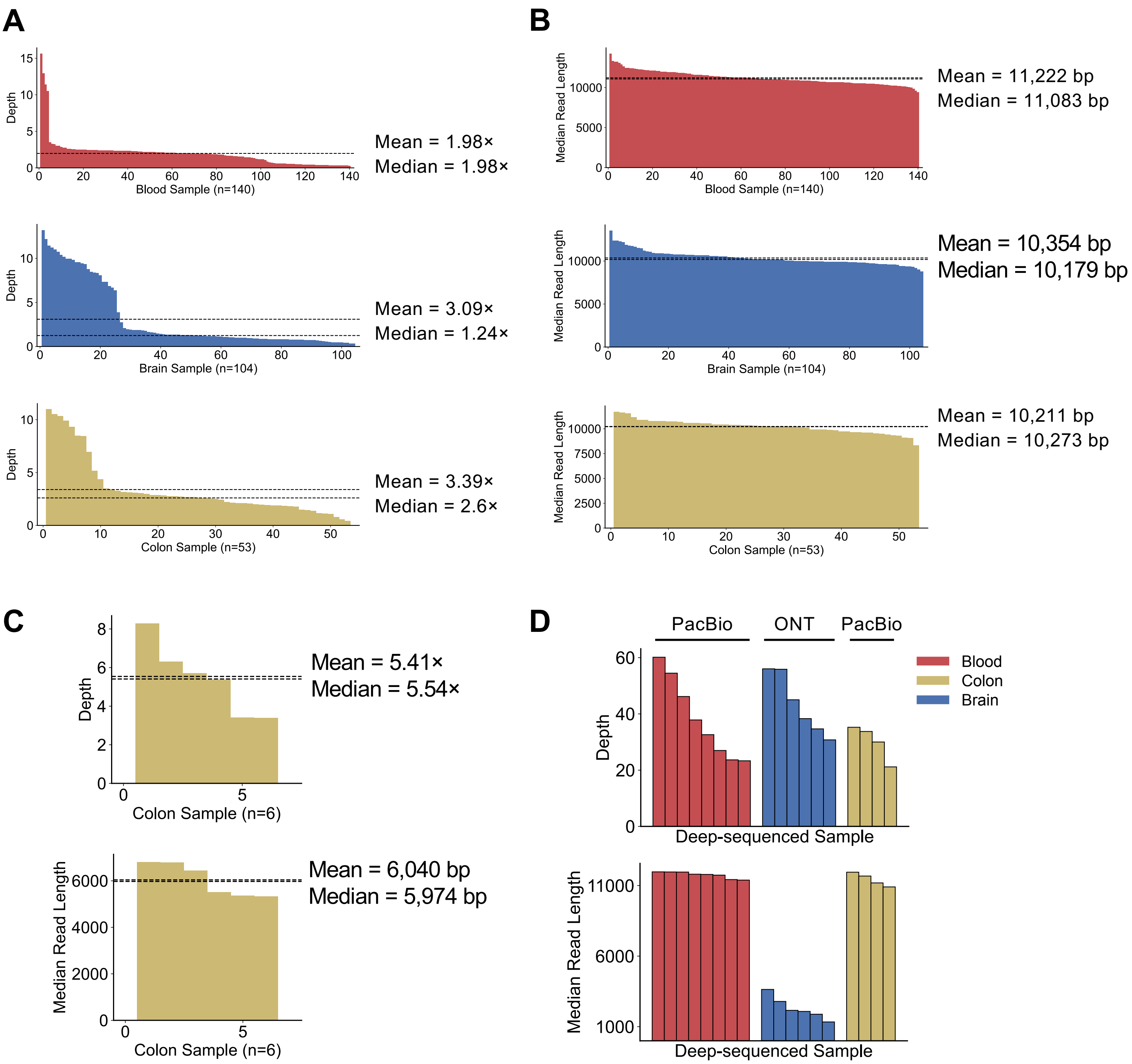


**Figure S1. Sequencing depth and read length across tissues for shallow and deep long-read datasets.**
(**A–B**) Genome-wide sequencing depth (**A**) and median read length (**B**) for the shallow-sequenced cohorts, comprising 140 blood, 104 brain and 53 colon samples, all PacBio-sequenced. (**C**) Genome-wide sequencing depth (upper) and median read length (lower) for 6 ONT-sequenced colon samples at shallow coverage. (**D**) Genome-wide sequencing depth (upper) and median read length (lower) for samples used in the deep-coverage analysis, comprising 6 blood, 6 brain and 4 colon samples. Blood and colon samples were PacBio-sequenced, and brain samples were ONT-sequenced, as indicated above the panel. In each panel, each bar represents one sample, bars are ordered by decreasing value. In (**A–C**) and horizontal dashed lines indicate the mean and median across samples of that tissue.

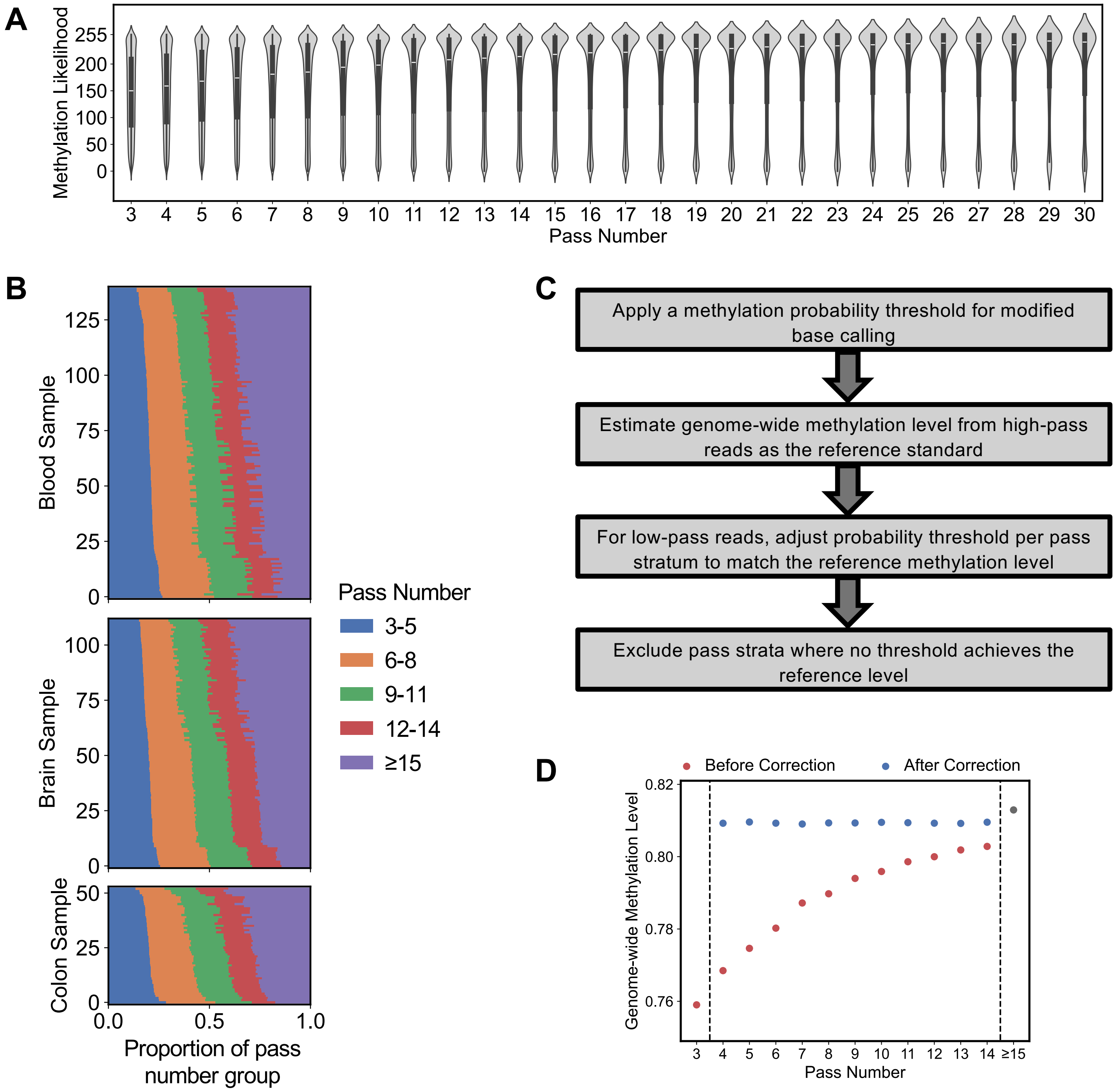


**Figure S2. Pass number correction for SMRT long-read methylation calling.** **(A)** Distribution of methylation likelihood (ML tag) values across pass numbers 3 to 30 for a representative sample, illustrating the systematic shift toward lower and more ambiguous values in low-pass reads relative to high-pass reads. **(B)** Pass number distributions across all cohort samples in this study, highlighting the inter-sample variability that motivates per-sample threshold calibration. **(C)** Schematic of the pass number correction pipeline implemented in COMET: a fixed probability threshold is applied to reads with ≥15 passes to establish a reference genome-wide methylation level, and pass-specific thresholds are then calibrated for reads with <15 passes to match this reference. Pass strata where no threshold achieves the reference level or where usable read depth is insufficient are excluded from further analysis. **(D)** Genome-wide CpG methylation levels at each pass number stratum before and after correction, for the representative sample in **(A)**. Corrected levels (blue) match the ≥15-pass reference (gray) from 4 passes onward; uncorrected levels (red) rise steadily with pass number. Reads with 3 passes have no threshold achieves the reference level and were excluded after correction.

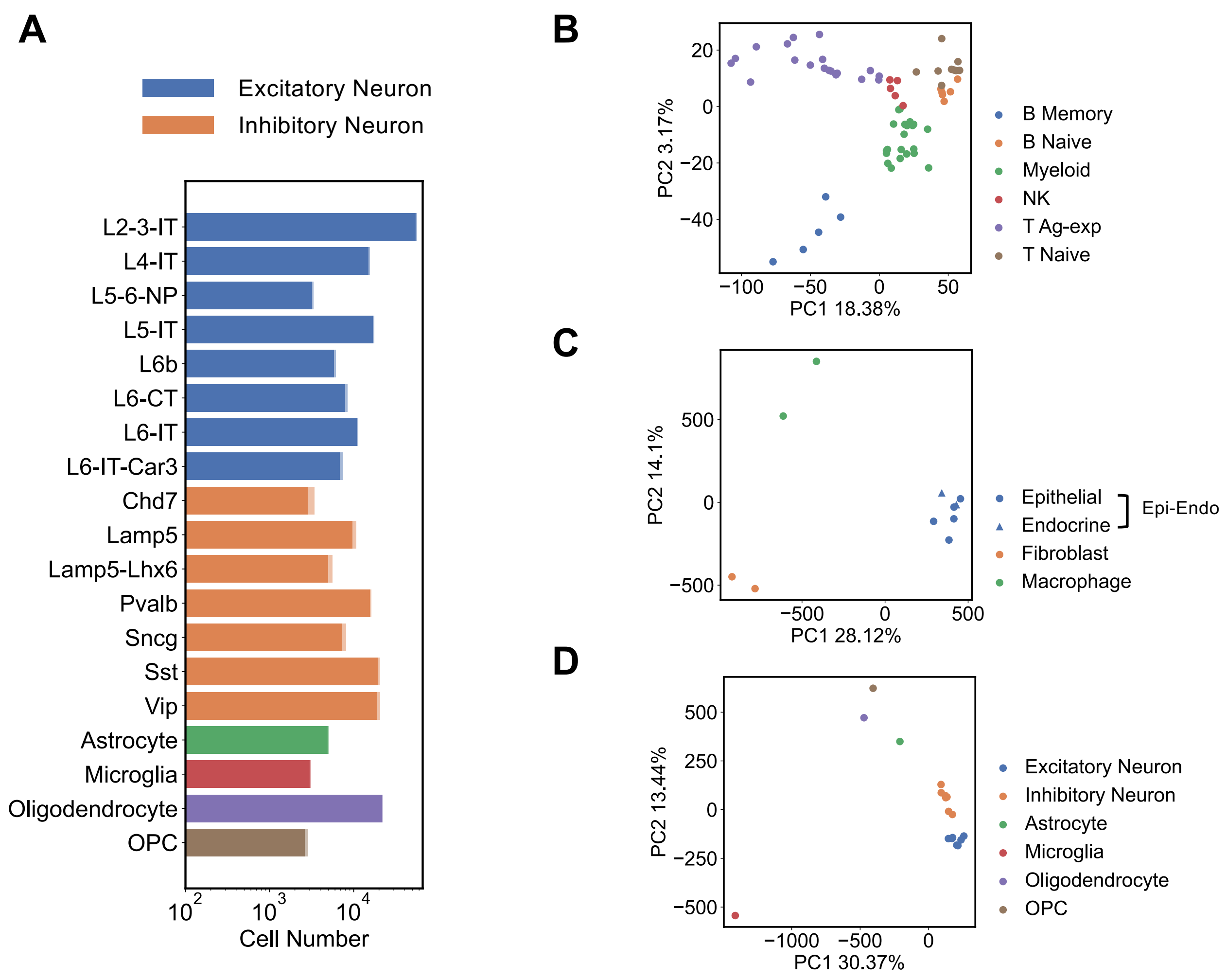


**Figure S3. Curated methylation reference panels across tissues.**
**(A)** Number of single cells per brain cell type contributing to the brain reference panel[1], shown on a log scale. For each cell type, the light shade gives the cell number reported in Tian et al. and the dark shade the number of cells successfully downloaded and used. **(B–C)** Principal component analysis of genome-wide CpG methylation for cell types in the **(B)** blood and **(C)** colon reference panels, based on high-depth BS-seq of sorted cells[2,3]. Each point represents one sorted-cell sample; axes show the first two principal components with the percentage of variance explained. **(D)** Principal component analysis of genome-wide CpG methylation for brain reference panel cell types, derived from single-cell methylation sequencing data aggregated to pseudo-bulk profiles per cell type, corresponding to **(A)**. Excitatory and inhibitory neuron subtypes are shown separately to assess whether their methylation profiles are sufficiently distinct to warrant independent representation or can be consolidated into a single class.

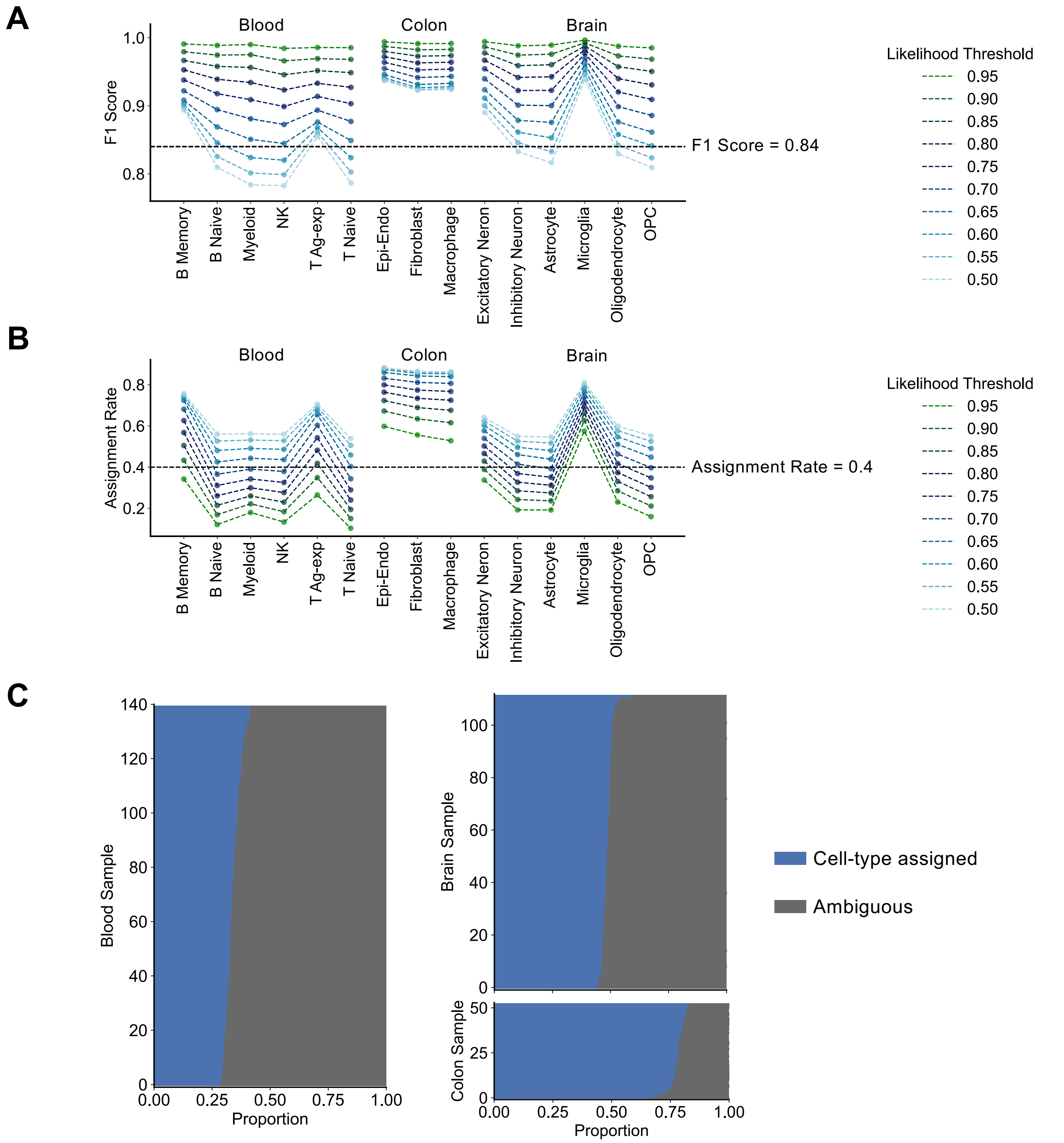


**Figure S4. Simulation-based evaluation of cell type assignment by COMET.** **(A)** F1 scores for each cell type in the blood, colon, and brain reference panels, computed from simulated reads of known cell-type origin at likelihood thresholds from 0.50 to 0.95 in increments of 0.05. The horizontal dashed line marks F1 score = 0.84, the minimum achieved by all cell types at the selected thresholds (0.65 for blood and brain, 0.50 for colon). **(B)** Assignment rate for each cell type in the blood, colon, and brain reference panels, computed from simulated reads of known cell-type origin at likelihood thresholds from 0.50 to 0.95 in increments of 0.05. The horizontal dashed line marks assignment rate = 0.4, the minimum achieved by all cell types at the selected thresholds (0.65 for blood and brain, 0.50 for colon). **(C)** Proportion of reads assigned to a cell type versus left ambiguous in each real sample from our cohort, under the selected thresholds (0.65 for blood and brain, 0.50 for colon). Each horizontal bar represents one sample, ordered by assignment rate.

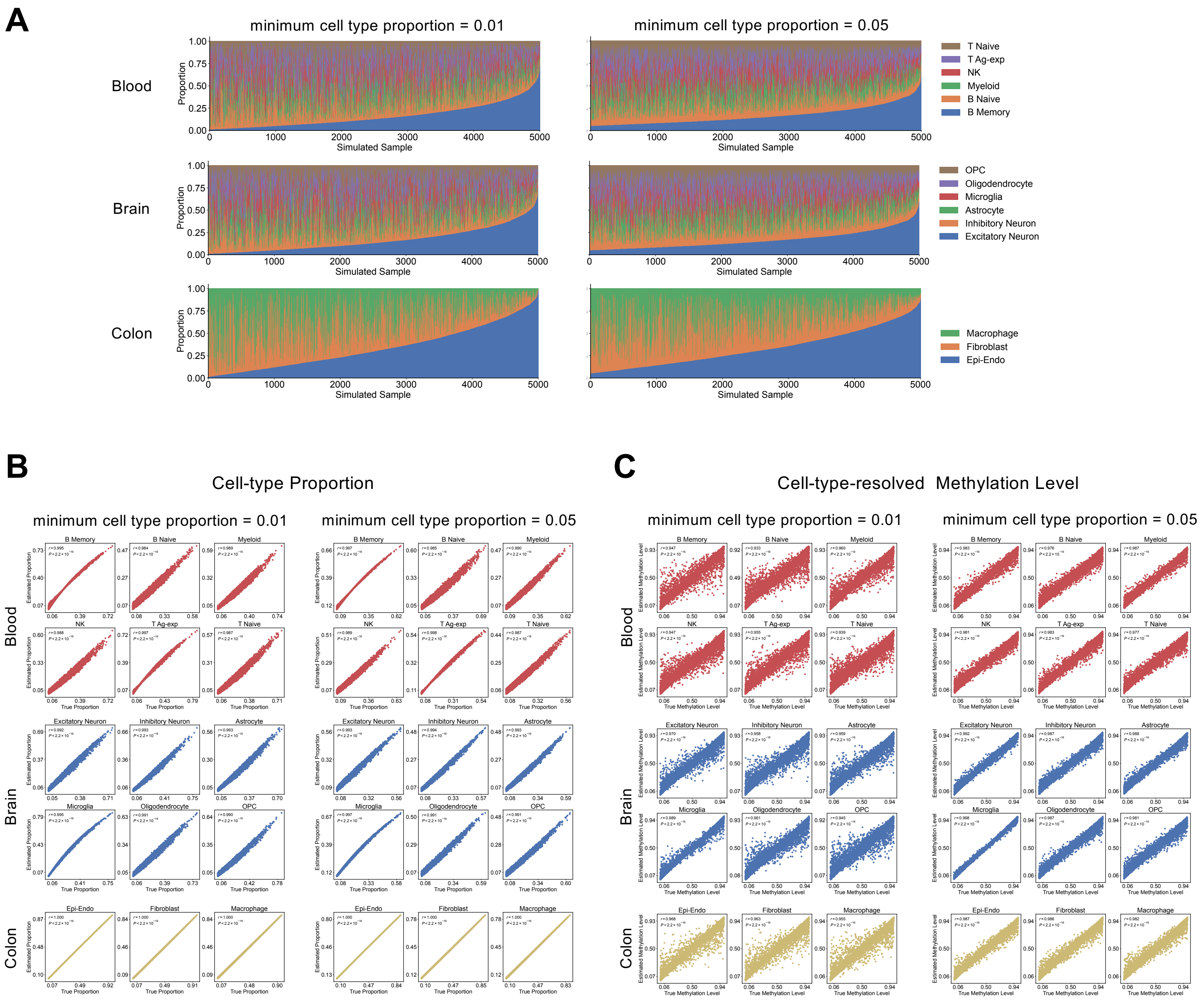


**Figure S5. Simulation-based evaluation of cell-type quantification analysis by COMET.**

**(A)** Composition of simulated mixtures for each tissue. Each bar represents one simulated sample with known cell-type proportions, generated such that every cell type exceeds a defined minimum proportion. **(B)** Scatter plots showing the correspondence between known cell-type proportions from simulated reads and proportions estimated by COMET, across all three tissues. Each point represents one simulated mixture. Axes show known proportion (x-axis) and COMET-estimated proportion (y-axis). Pearson's *r* and associated *p* value are labelled for each cell type. **(C)** Scatter plots showing the correspondence between known cell-type methylation levels from simulated reads and cell-type-resolved methylation levels estimated by COMET. Each point represents the methylation level of one cell type from one simulated sample. Axes show known methylation level (x-axis) and COMET-estimated methylation level (y-axis). Pearson's *r* and associated *P* value are labelled for each cell type.

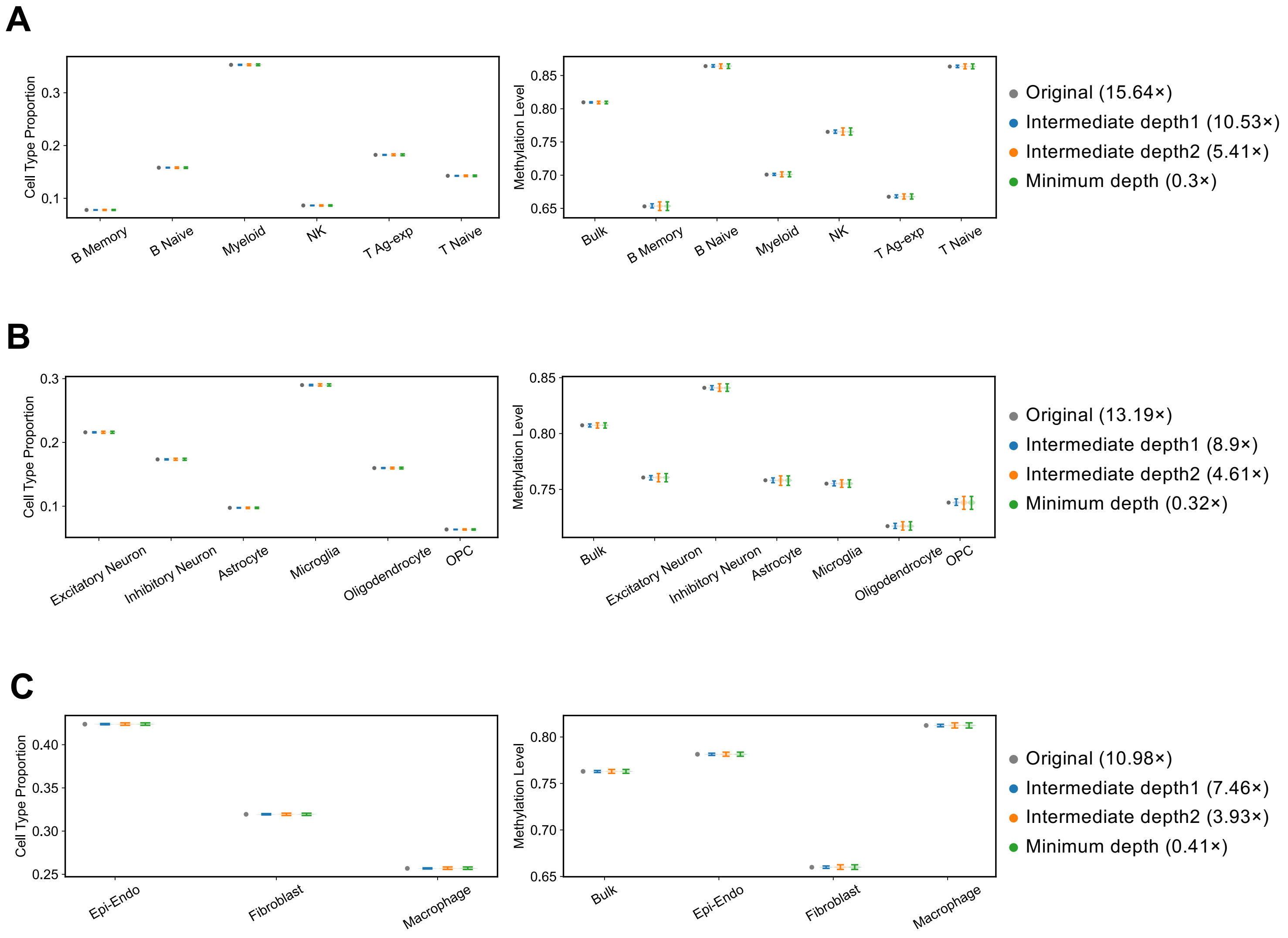


**Figure S6. Cell-type quantification analysis by COMET is robust to sequencing depth.**

**(A–C)** The highest-depth sample from each tissue cohort was downsampled 2,000 times to each of three target depths, two intermediate depths and the lowest depth observed in that cohort, for **(A)** blood, **(B)** brain, and **(C)** colon. Left panels show cell-type proportions and right panels show genome-wide methylation levels for bulk tissue and for each cell type. Grey points indicate values from the original, undownsampled sample, with its depth given in the legend. Blue, orange, and green violins show the distribution across the 2,000 downsampling replicates at each target depth, with depths given in the legend for each tissue.

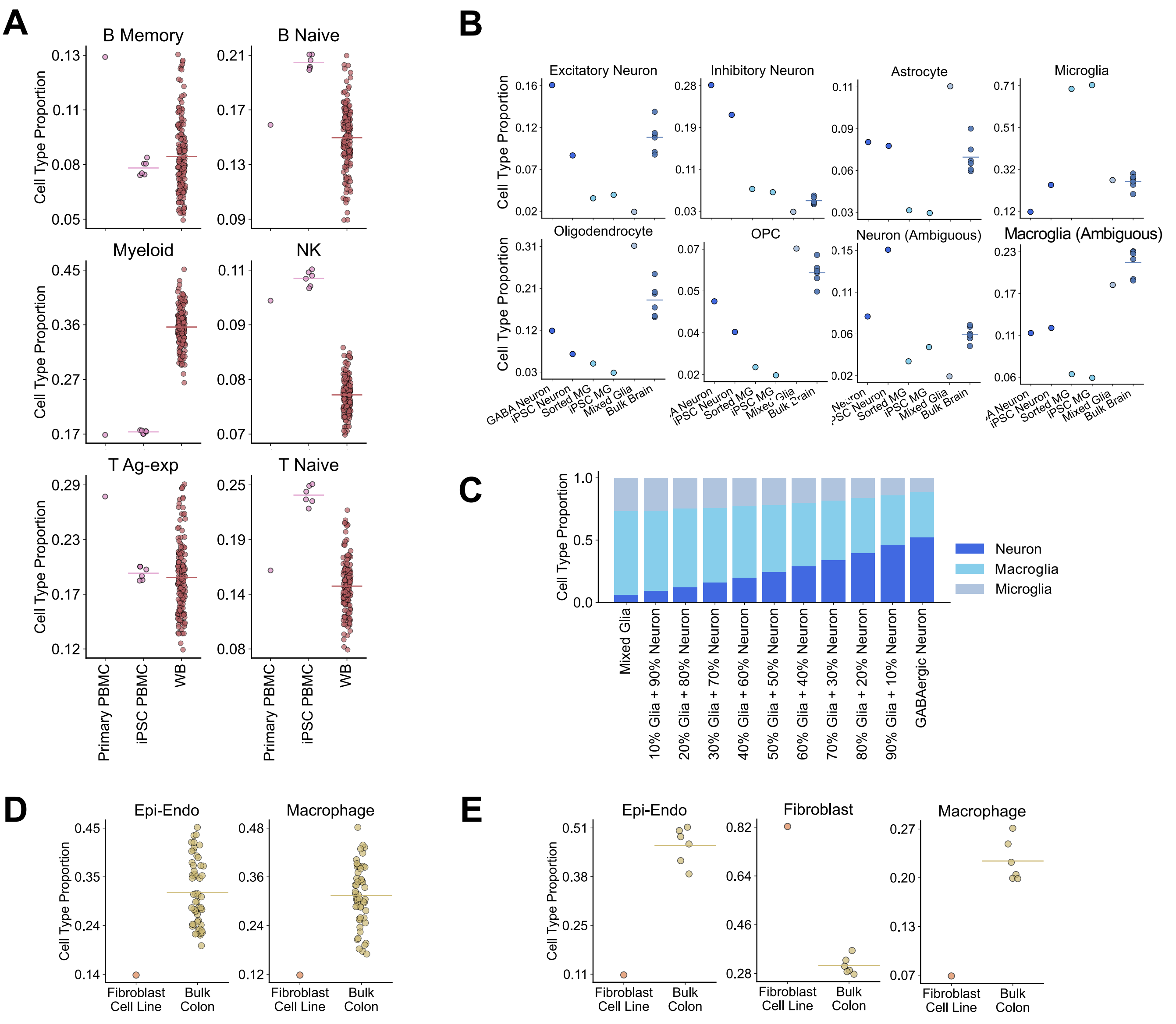


**Figure S7. Real-sample evaluation of cell type assignment.**
**(A)** Cell-type proportions estimated from assigned reads across the six blood cell types in PacBio data, shown for one primary PBMC sample, six iPSC-derived PBMC samples, and 140 whole-blood (WB) samples. **(B)** Cell-type proportions across the six brain cell types and two broad cell groups (ambiguous neuron and ambiguous macroglia) in ONT data, shown for one GABAergic neuron sample, one iPSC-derived neuron sample, one sorted microglia sample, one iPSC-derived microglia sample, one mixed-glia sample, and six bulk brain samples. **(C)** Cell-type proportions of the mixed-glia and GABAergic neuron samples alongside titration samples generated by mixing reads from the two at defined proportions, showing recovery of the imposed composition gradient. **(D)** Cell-type proportions for Epi-Endo and macrophage in PacBio colon data, shown for one colon fibroblast cell line and bulk colon samples; the corresponding fibroblast result is shown in **Fig. 2G**. **(E)** Cell-type proportions across the three colon cell types in ONT data, shown for one colon fibroblast cell line and six bulk colon samples. In **(A, B, D, and E)**, each point represents one sample and horizontal lines indicate the group mean.

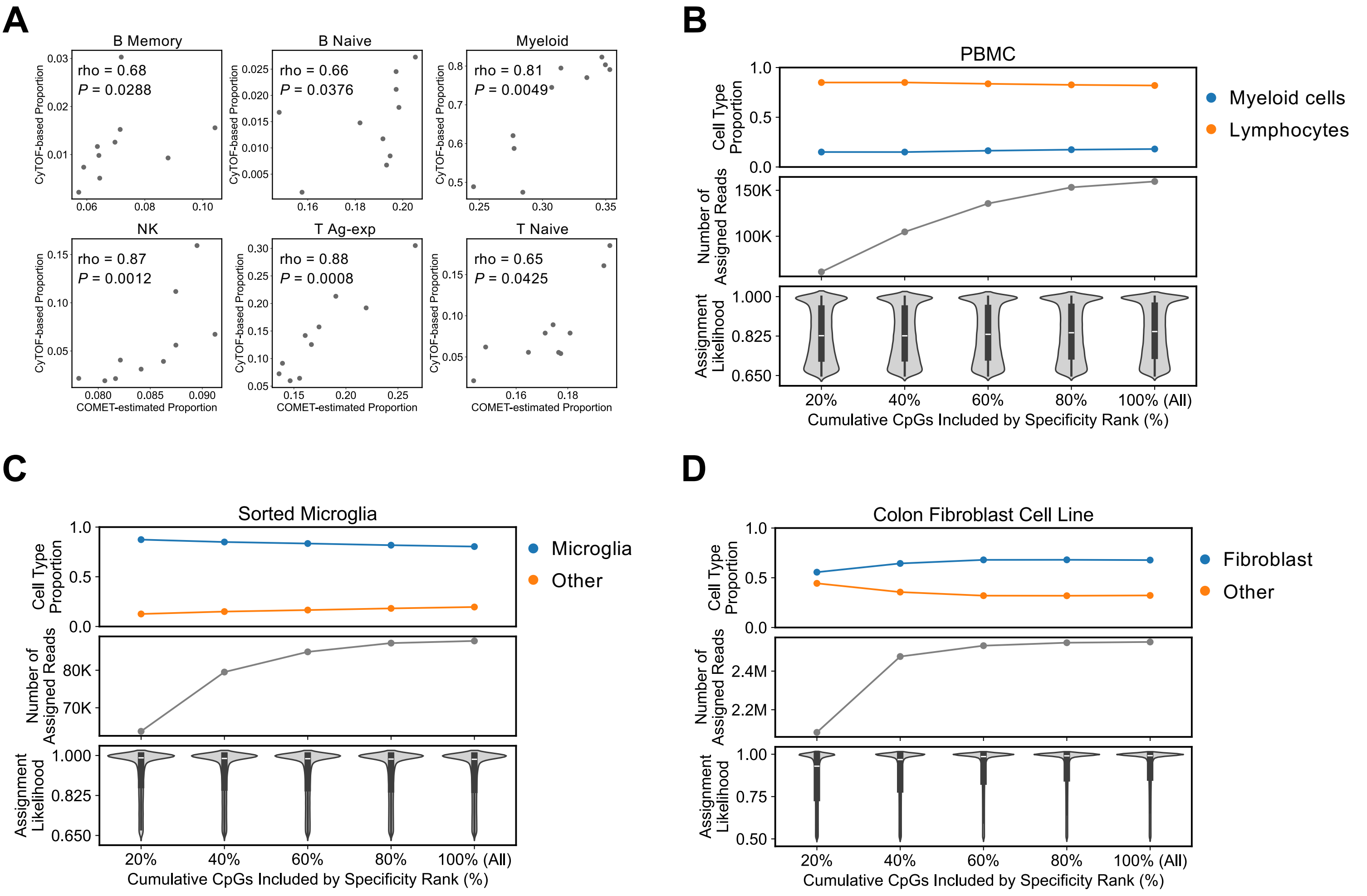


**Figure S8. COMET proportions are concordant with CyTOF and stable across reference panel sizes.**

**(A)** COMET-estimated versus CyTOF-based cell type proportions in an independent set of 10 blood samples profiled by both PacBio SMRT sequencing and CyTOF, shown separately for each of the six blood cell types. Each point represents one sample, and Spearman's rho and *P* value are given in each panel. **(B–D)** Samples of known cellular composition were assigned with COMET using the **(B)** blood, **(C)** brain, and **(D)** colon reference panels, restricting the panel to the top 20%, 40%, 60%, 80%, or 100% of CpGs ranked by cell-type specificity (x-axis, cumulative). The samples shown are a PBMC preparation in **(B)**, a sorted microglia sample in **(C)**, and a colon fibroblast cell line in **(D)**, each dominated by a known cell type. Top: estimated cell-type proportions, shown for the expected dominant population and the remainder. Middle: total number of reads receiving a confident cell-type assignment. Bottom: distribution of assignment likelihoods across all assigned reads, shown as violins with the median and interquartile range overlaid. Proportion estimates and likelihood distributions are essentially unchanged as lower-specificity CpGs are added, whereas the number of assigned reads rises steadily, most steeply between the top 20% and 40%.

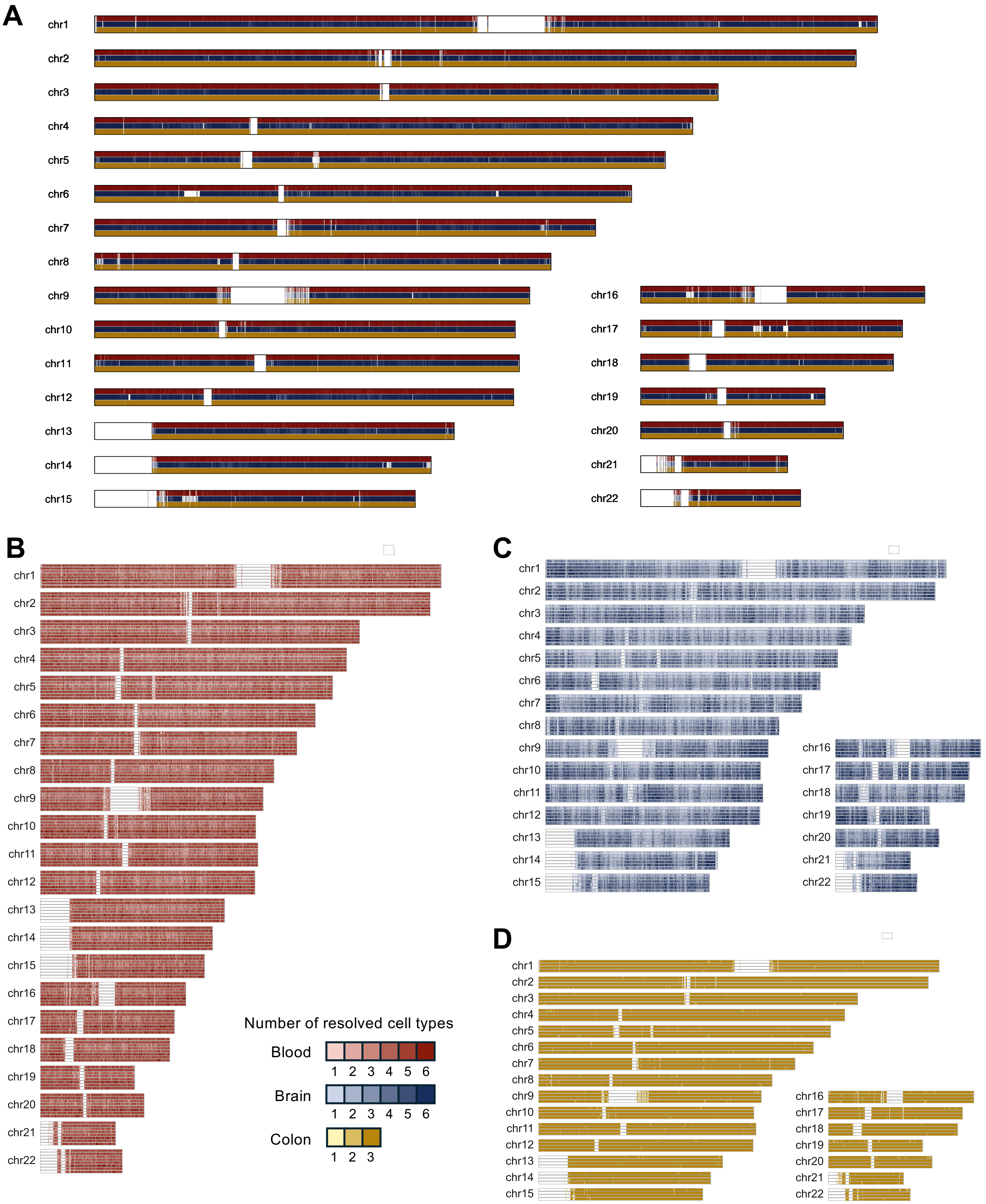


**Figure S9. Number of cell types resolved by COMET across the genome.**
**(A)** Number of cell types resolved across the genome, based on cell-type-assigned reads aggregated across all shallow PacBio-sequenced samples of each tissue. Within each chromosome, tracks run from top to bottom as blood (red), brain (blue), and colon (yellow) with six, six and three cell types respectively. **(B–D)** Number of cell types resolved across the genome for **(A)** 8 blood, **(B)** 6 brain, and **(C)** 4 colon deep-sequenced samples. Within each chromosome, one track is drawn per sample. In each color scheme of (**A–D**), lighter shades indicate fewer cell types resolved and darker shades more. Regions where no cell type could be resolved are left blank.

The red and blue marks indicate methylated and unmethylated CpGs, respectively

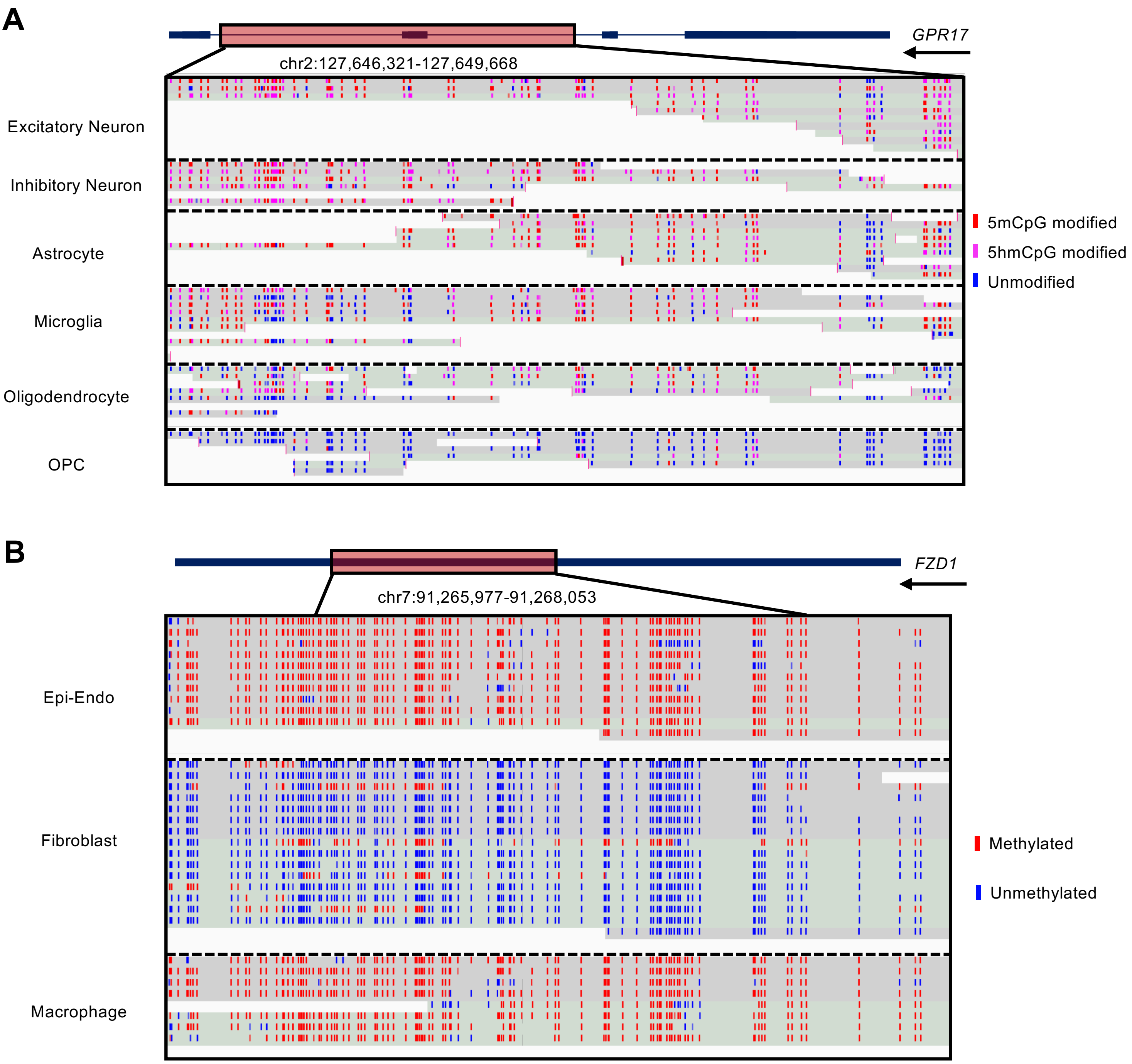


**Figure S10. Read-level visualization of cell-type-specific methylation at marker loci in deep-sequenced blood and colon samples.**
**(A)** Genome browser view of the *GPR17* locus (chr2:127,646,321–127,649,668) in a representative deep ONT-sequenced brain sample, showing reads assigned to each of the six brain cell types. OPC-assigned reads are largely unmodified across the region, whereas reads from the other five cell types are predominantly modified. The red, magenta and blue marks indicate 5mCpG-modified, 5hmCpG-modified and unmodified CpGs, respectively. **(B)** Genome browser view of the *FZD1* locus (chr7:91,265,977–91,268,053) in a representative deep-sequenced colon sample, showing reads assigned to each of the three colon cell types. Fibroblast-assigned reads carry markedly more unmethylated CpGs than Epi-Endo- and macrophage-assigned reads. The red and blue marks indicate methylated and unmethylated CpGs, respectively. In both panels, each row represents one read, grouped by assigned cell type and separated by dashed lines. Arrow under the gene name indicates the orientation of the gene.

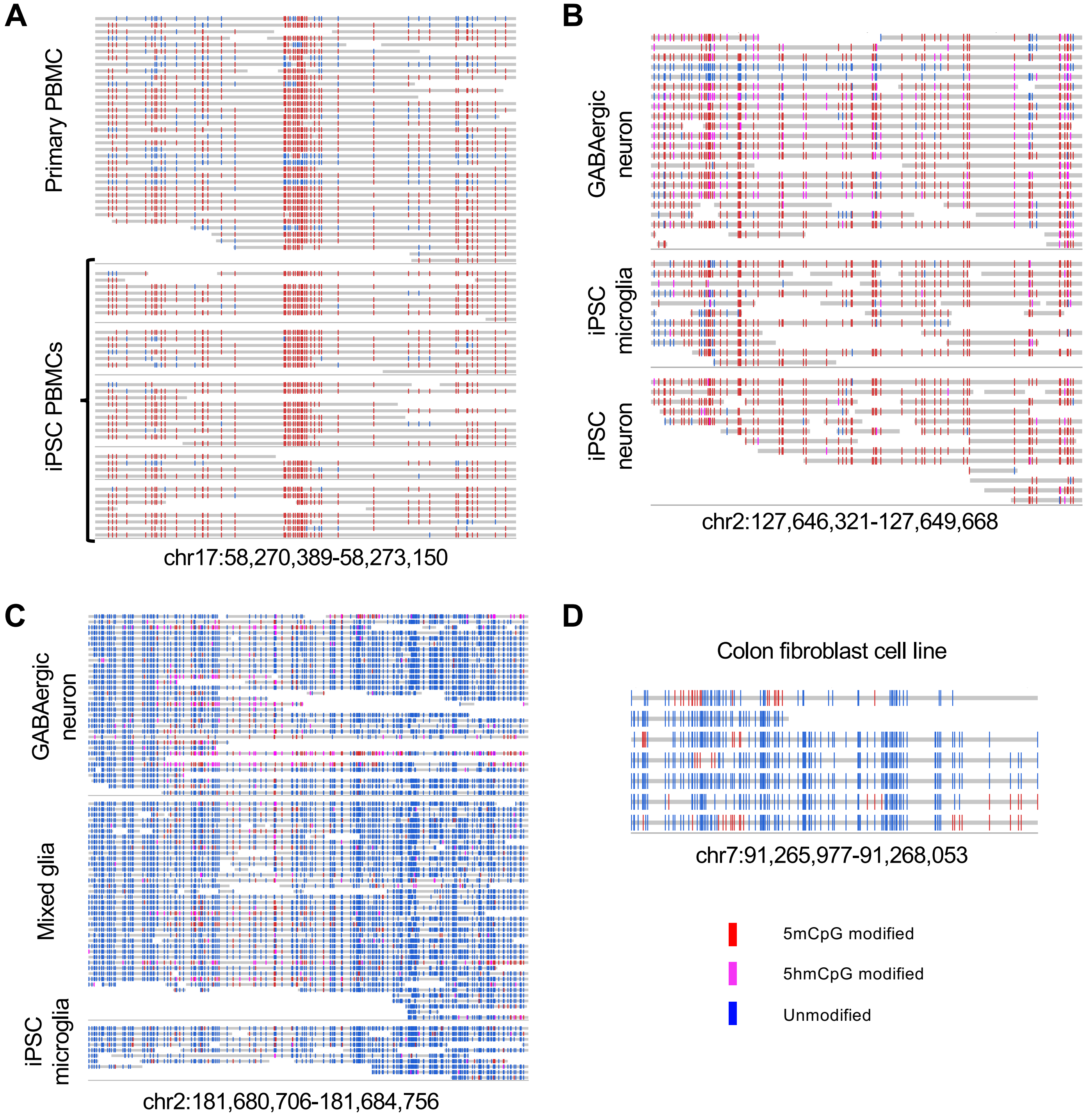


**Figure S11. Cell-type-specific modification patterns confirmed in samples of known cell-type identity.**

Read-level modification at the four example loci of **Fig. 2E–F** and **Supplementary Fig. 10A–B**, shown in samples composed predominantly of a single cell type. Each row represents one read. Red, magenta and blue marks indicate 5mCpG-modified, 5hmCpG-modified and unmodified CpGs, respectively. **(A)** *MPO* (chr17:58,270,389–58,273,150; **Fig. 2E**) in primary and iPSC-derived PBMC samples, which are lymphocyte-enriched and show predominantly methylated reads. **(B)** *GPR17* (chr2:127,646,321–127,649,668; **Supplementary Fig. 10A**) in GABAergic neuron, iPSC-derived microglia and iPSC-derived neuron samples, none of which contains OPCs and all of which are predominantly modified. **(C)** *NEUROD1* upstream (chr2:181,680,706–181,684,756; **Fig. 2F**) in GABAergic neuron, mixed glia and iPSC-derived microglia samples, none of which is an excitatory neuron population and all of which lack the 5hmCpG signal seen on excitatory-neuron-assigned reads. **(D)** *FZD1* (chr7:91,265,977–91,268,053; **Supplementary Fig. 10B**) in a colon fibroblast cell line, showing predominantly unmethylated reads.

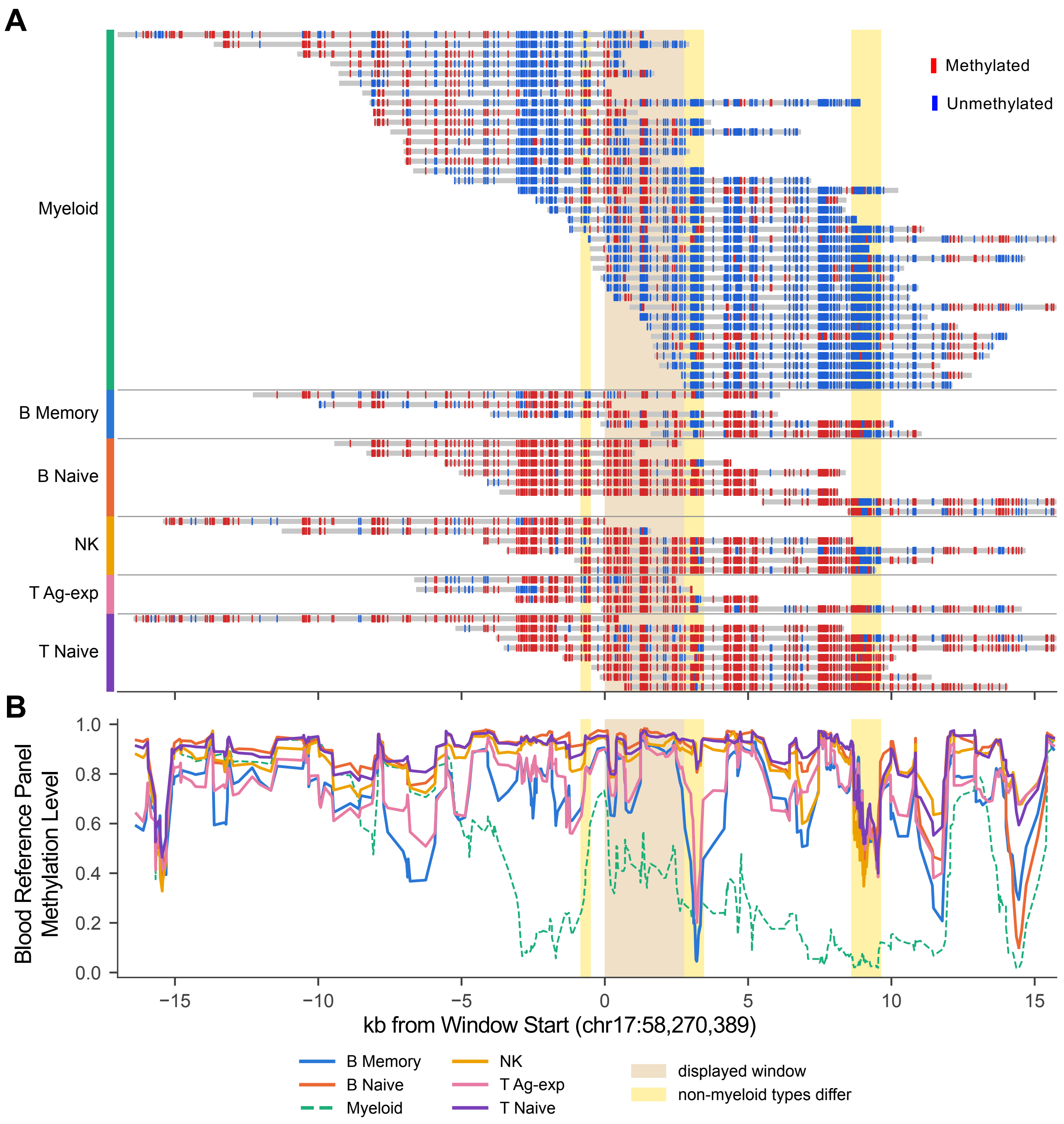


**Figure S12. Cell-type assignment of reads overlapping a window draws on CpGs outside it.**
**(A)** Expanded view of COMET-assigned reads for the blood sample in **Fig. 2E**. Each row is one primary alignment. Reads are ordered by assigned cell type. Blue and red mark unmethylated and methylated CpGs. Brown shading marks the displayed window from **Fig. 2E**, chr17:58,270,389–58,273,150, where myeloid reads at the top are largely unmethylated and the other five types are largely methylated. This pattern separates myeloid from the rest but leaves the five non-myeloid types indistinguishable from one another. Yellow shading marks flanking regions where the non-myeloid types differ from one another and track their reference profiles. **(B)** Blood reference panel methylation level for the six blood cell types across the same interval and x-axis as **(A)**. The dashed line is myeloid, and solid lines are the other five. The x-axis gives distance in kb from the window start at chr17:58,270,389.

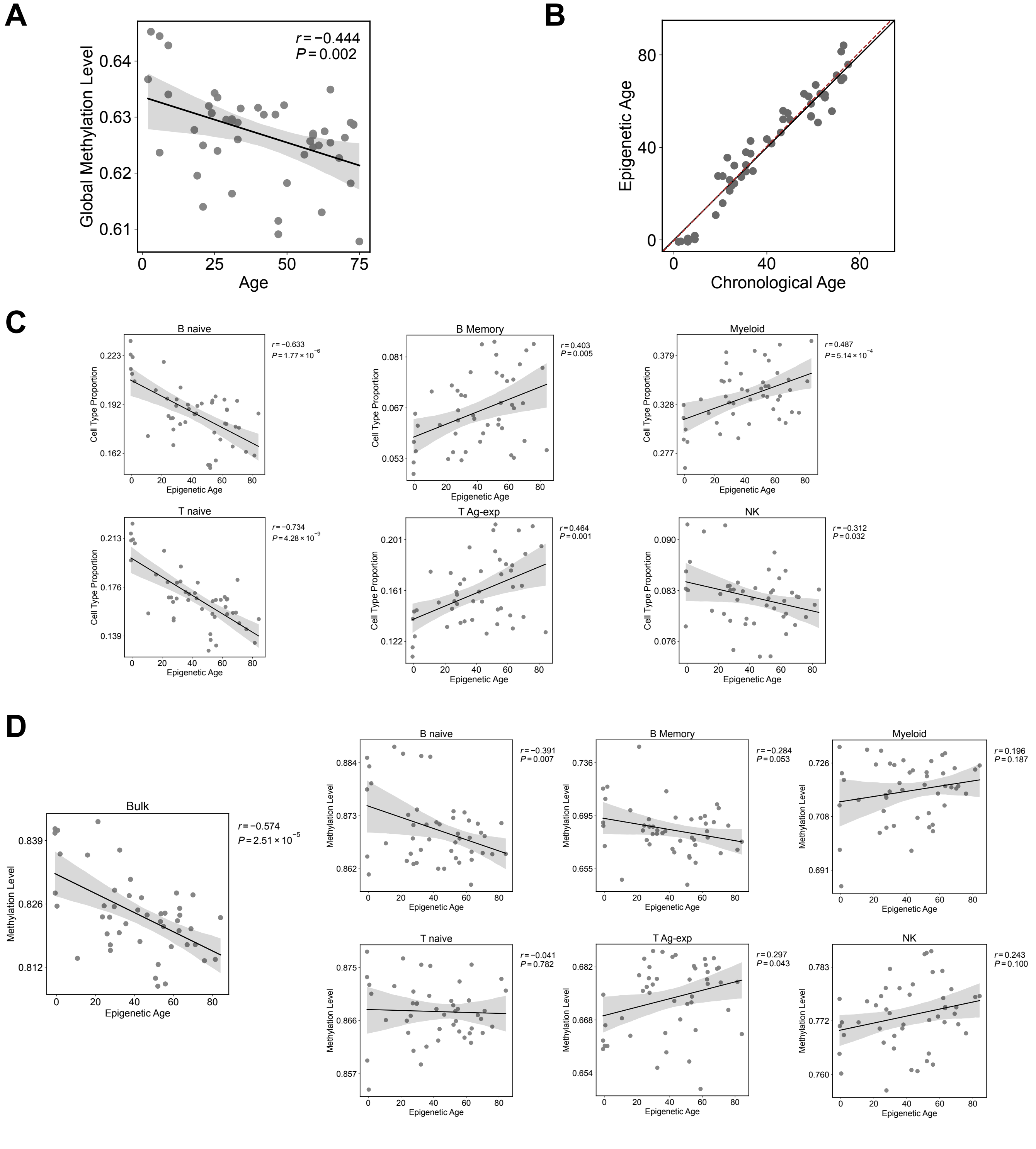


**Figure S13. Methylation array-based epigenetic age compared with long-read cell-type quantification measures.**

Analyses use the 47 blood samples with matched methylation array and long-read data. **(A)** Methylation array-based global methylation level against chronological age. **(B)** Methylation array-based Horvath clock age[4] against chronological age. The solid black line marks the diagonal where epigenetic age equals chronological age; the red dashed line is the fitted regression. **(C)** Methylation array-based Horvath clock age against the proportion of each of the six blood cell types, estimated from long-read data. **(D)** Methylation array-based Horvath clock age against genome-wide methylation level in bulk blood (left) and in each cell type resolved from long-read data (right). Throughout, points are individual samples, lines are linear fits, and shaded bands are 95% confidence intervals. Correlations are Pearson's *r* adjusted for sex, with *r* and the associated *P* value shown in each panel.

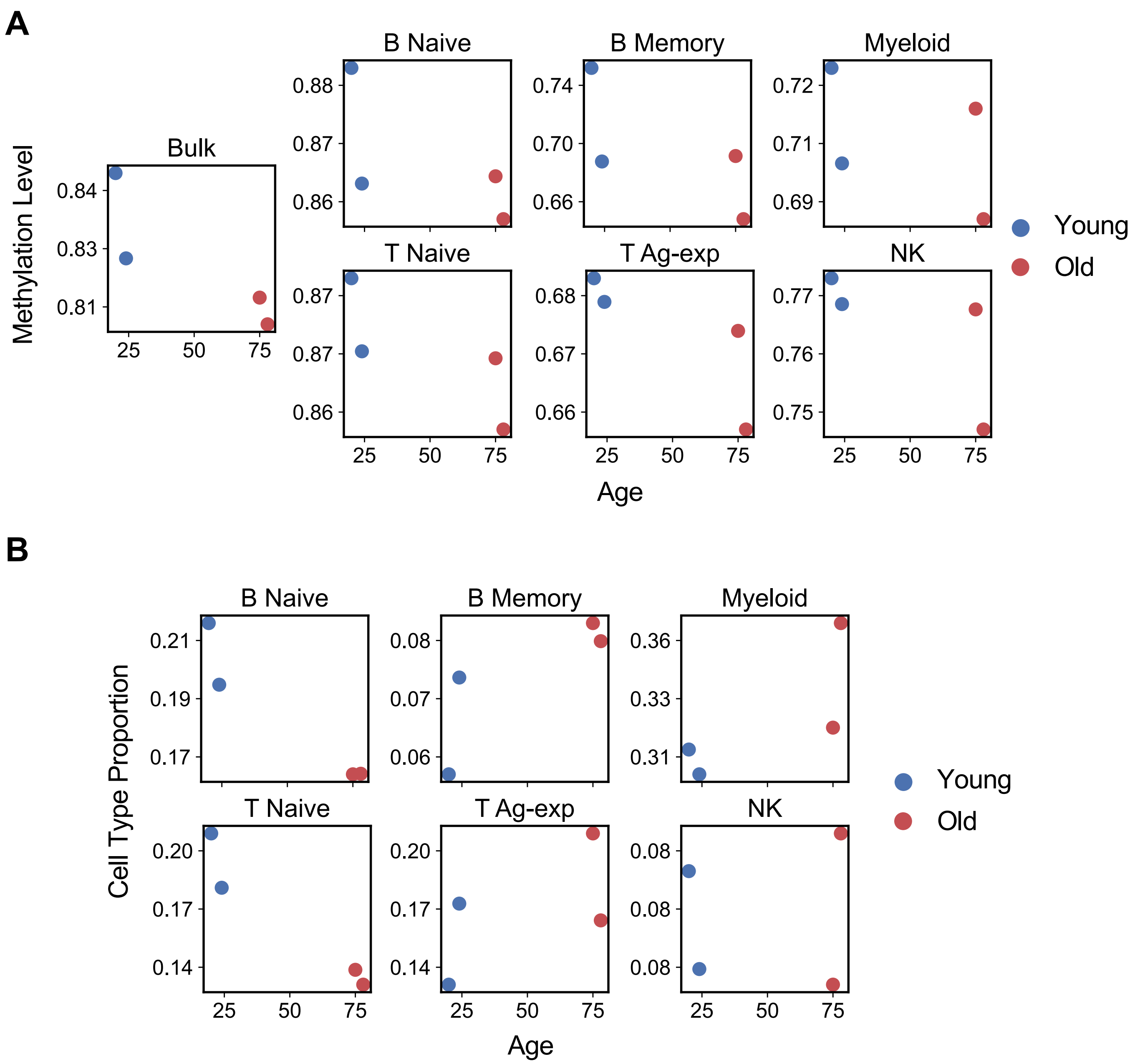


**Figure S14. Cell-type quantification measures of the four blood samples selected for deep sequencing.**
Four blood samples selected for high-coverage sequencing, two young (blue) and two old (red), plotted against chronological age. **(A)** Genome-wide methylation level in bulk blood (left) and in each of the six cell types resolved by COMET (right). **(B)** Proportion of each of the six cell types. Each point represents one sample.

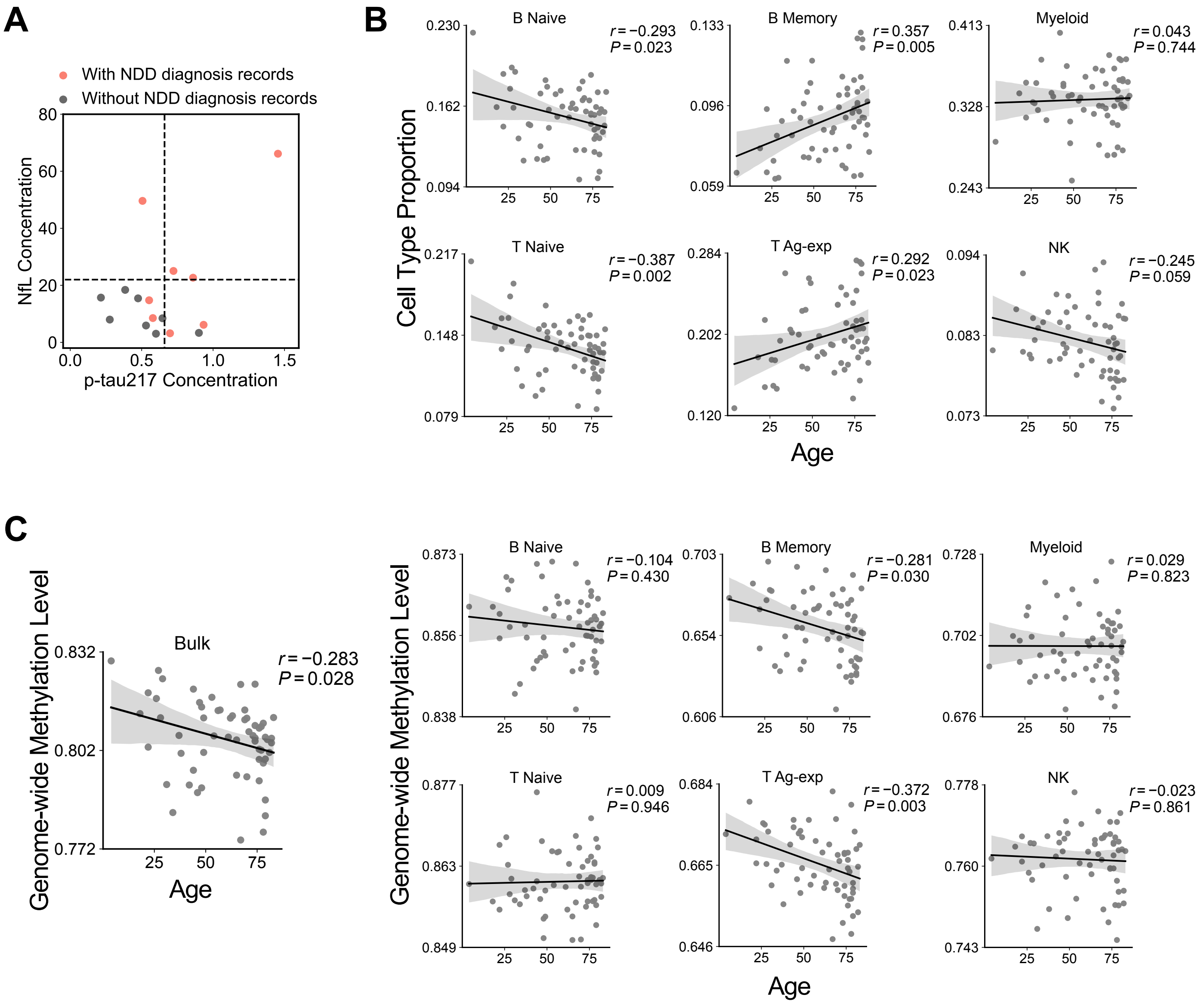


**Figure S15. Identification of resilience candidates among APOE4 carriers and cell-type aging patterns in APOE4 carriers.**
**(A)** Chart review of NDD diagnosis records, plasma p-tau217 and NfL for all APOE4 carrier blood samples aged ≥75 years. Each point represents one subject, with p-tau217 concentration on the x-axis and NfL concentration on the y-axis. Dashed lines mark the thresholds used to flag elevated NDD risk, 0.66 for p-tau217 and 22 for NfL[5]. Red points indicate subjects with an NDD diagnosis on record and grey points those without. Grey points falling below both thresholds, in the lower left quadrant, were defined as resilience candidates (n = 7), and all remaining subjects were classified as non-resilient (n = 9). **(B)** COMET-estimated cell-type proportions across age in APOE4 carriers. **(C)** COMET-estimated genome-wide methylation levels across age in APOE4 carriers, shown for bulk blood (left) and each cell type (right). In **(B–C)**, each plot gives the indicated measure (y-axis) against subject age (x-axis), with solid lines showing linear regression fits and shaded areas 95% confidence intervals. Pearson's *r* and *P* values, adjusted for sex, are shown in each plot.

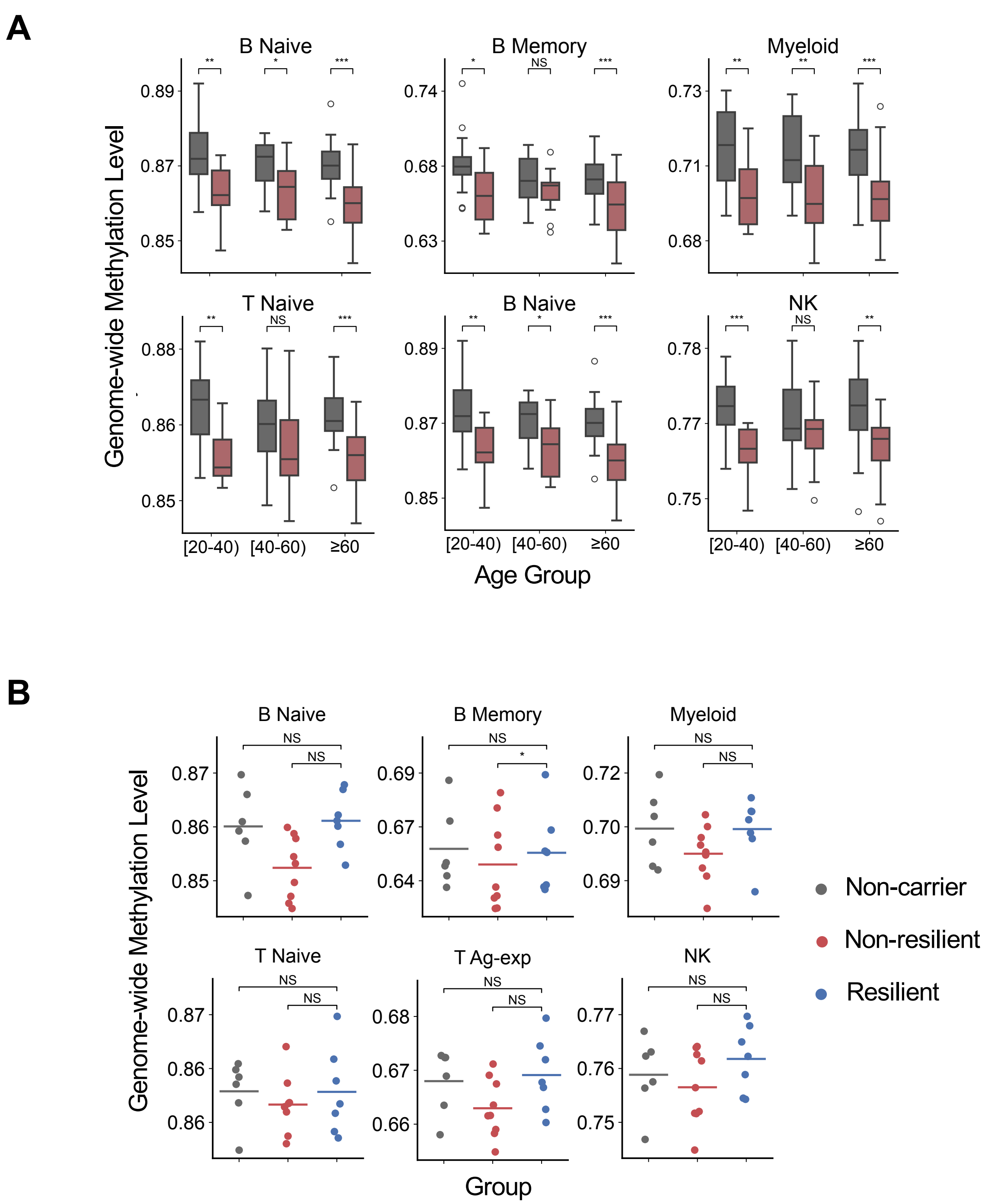


**Figure S16. Cell-type-resolved methylation across APOE4 genotype and resilience groups.**
**(A)** COMET-estimated genome-wide methylation levels for each blood cell type in APOE4 non-carriers (grey) and carriers (red), shown separately for three age groups. Boxes give the interquartile range and median, whiskers extend to 1.5× the interquartile range, and points beyond are outliers. **(B)** The same measures in non-carrier (grey), non-resilient (red) and resilience candidate (blue) subjects, all aged ≥75. Each point represents one sample and horizontal lines indicate group means. In both panels, asterisks indicate Benjamini–Hochberg-adjusted $P$ values from mixed-effects models adjusted for age and sex, with *$P_{adj} < 0.05$, ** $P_{adj} < 0.01$ and *** $P_{adj} < 0.001$. NS, not significant.

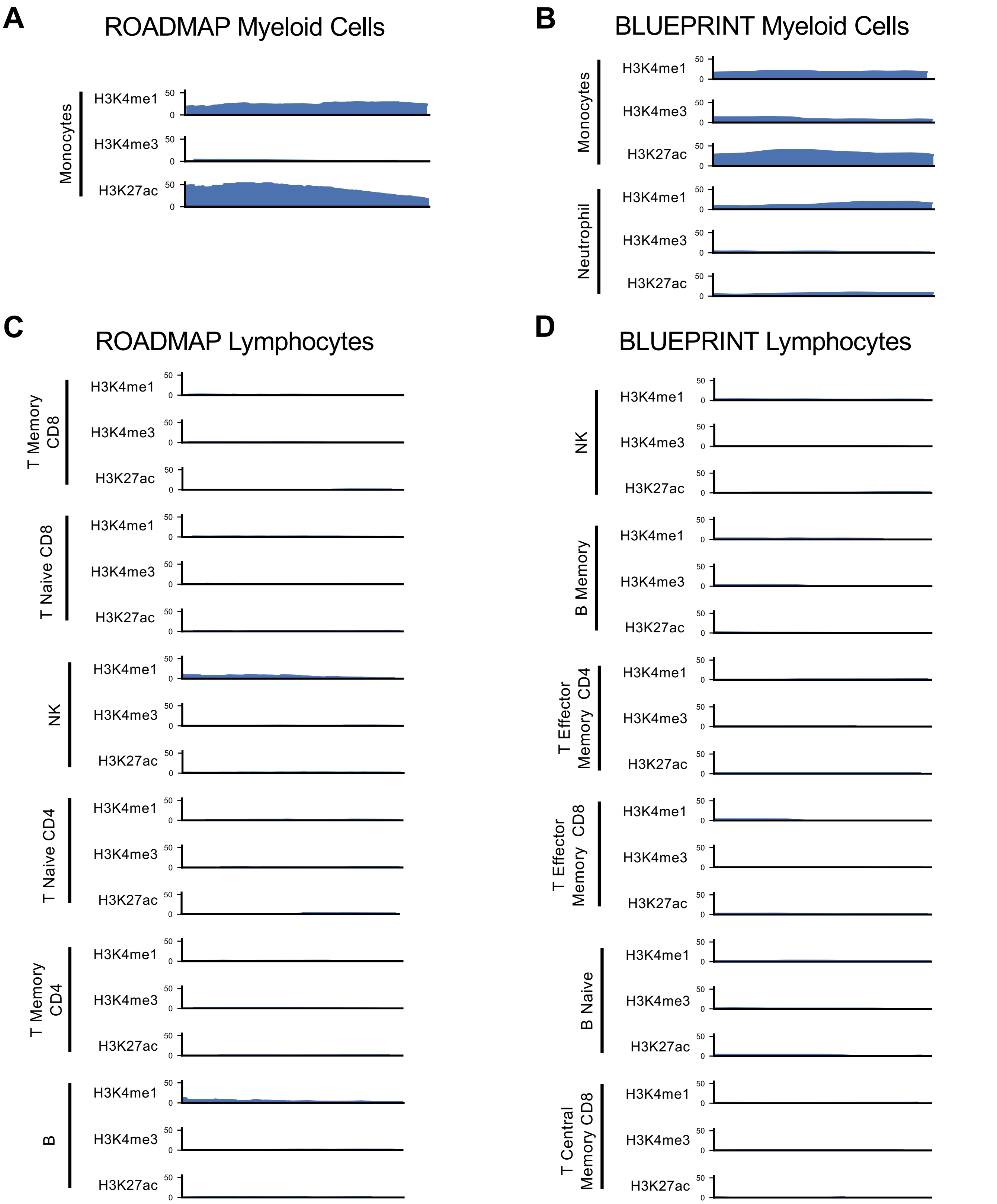


**Figure S17. Histone modification profiles at chr2:113,124,820–113,125,150 across individual myeloid and lymphocyte populations.**

**(A–D)** H3K4me1, H3K4me3 and H3K27ac signal (top to bottom within each cell population) across the region, shown for individual cell types rather than aggregated myeloid or lymphocyte tracks. Panels give **(A)** ROADMAP[6] myeloid populations, **(B)** BLUEPRINT[3] myeloid populations, **(C)** ROADMAP lymphocyte populations and **(D)** BLUEPRINT lymphocyte populations.

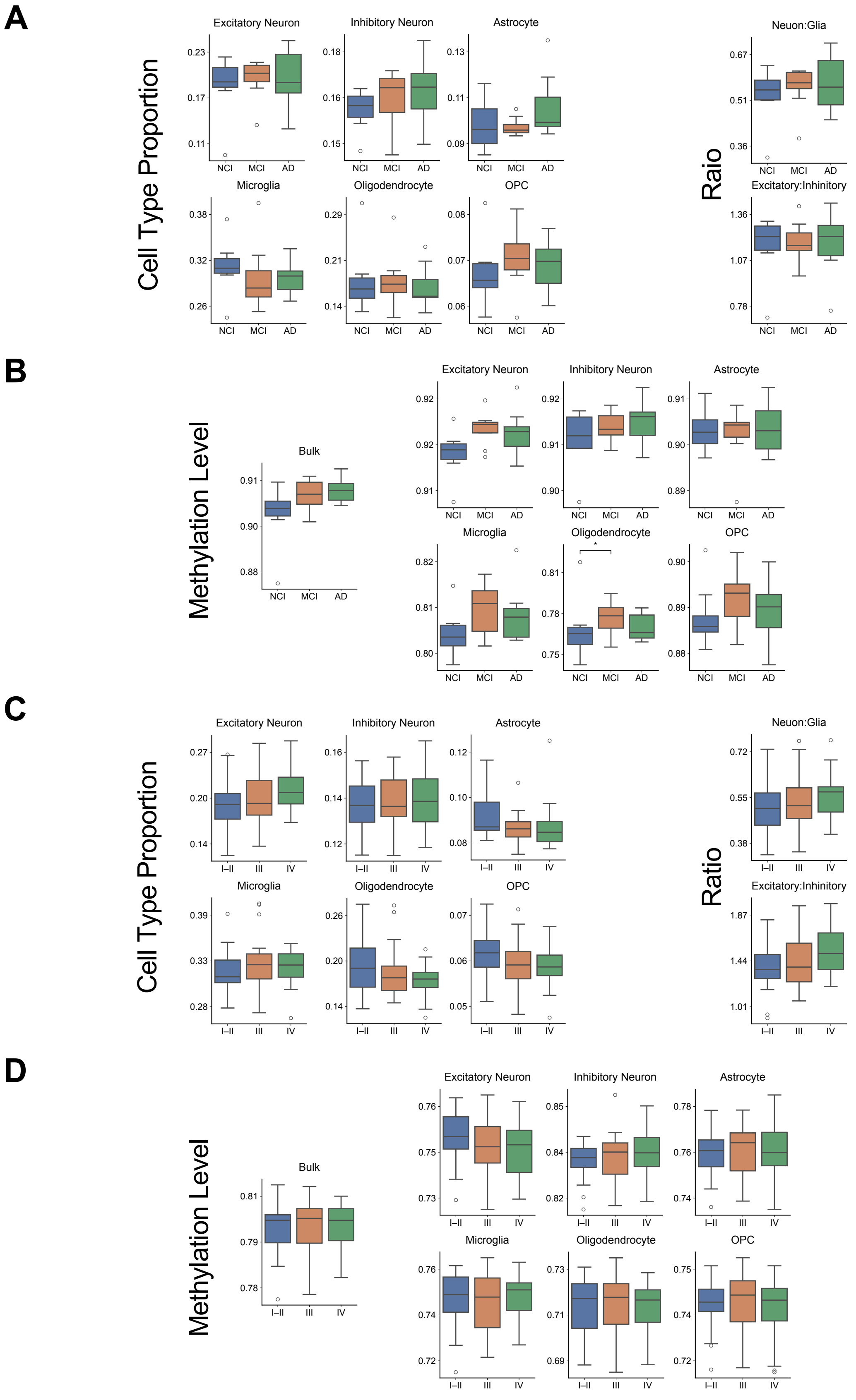


**Figure S18. Cell-type-resolved proportion and methylation measures against clinical traits in two brain cohorts.**

**(A–B)** NPBB cohort (n = 28), across cognitive diagnosis groups (NCI, no cognitive impairment; MCI, mild cognitive impairment; AD, Alzheimer's disease). **(A)** Bulk and cell-type-resolved methylation levels. **(B)** Cell-type proportions (left) and derived ratios (right, neuron-to-glia and excitatory-to-inhibitory). **(C–D)** PWG cohort (n = 76), across Braak stage groups (I–II, III, IV). **(C)** Bulk and cell-type-resolved methylation levels. **(D)** Cell-type proportions (left) and derived ratios (right). Asterisks indicate Benjamini–Hochberg-adjusted $P$ values from mixed-effects models adjusted for age and sex, with the bracket marking the significant pairwise contrast. $*P_{adj} < 0.05$.

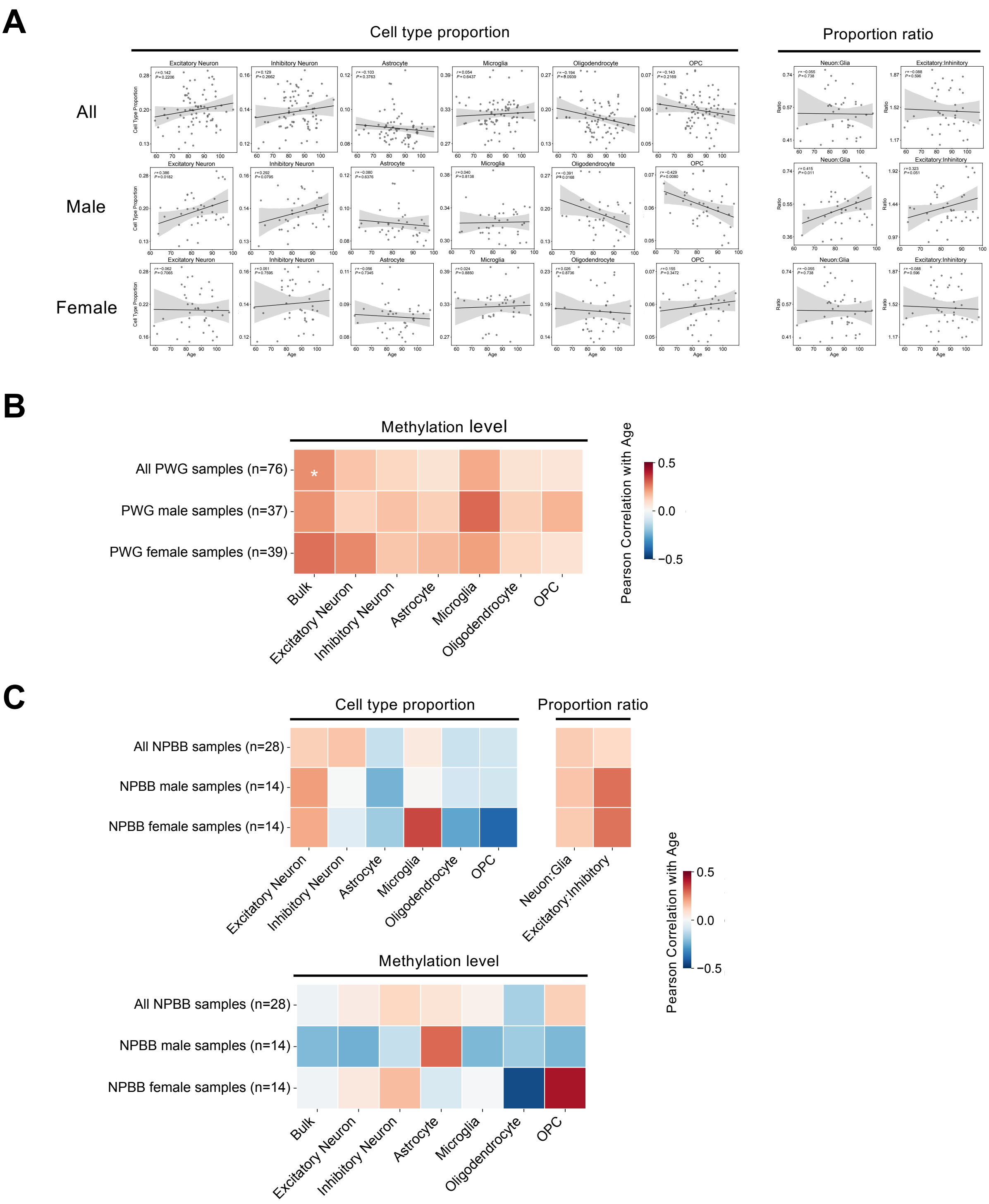


**Figure S19. Age associations of cell-type composition and methylation in the PWG and NPBB cohorts, overall and by sex.**

**(A)** Per-sample data underlying the correlations summarized in Fig. 6B. Cell-type proportions and proportion ratios (y-axis) are plotted against age (x-axis) in all PWG samples (n = 76), PWG male samples (n = 37) and PWG female samples (n = 39), top to bottom. Each point represents one sample, with solid lines giving linear regression fits and shaded areas 95% confidence intervals. Braak stage-adjusted Pearson's *r* and P values are shown in each plot. **(B)** Associations between age and bulk or cell-type-resolved methylation level in the same three PWG groups. Color gives the Pearson correlation coefficient, with asterisks indicating Braak stage-adjusted *P* values, *$P_{adj}$ < 0.05 and **$P_{adj}$ < 0.01. **(C)** The same correlation analyses in the NPBB cohort, shown for all NPBB samples (n = 28), male samples (n = 14) and female samples (n = 14). Upper panels give cell-type proportions and proportion ratios, lower panels give bulk and cell-type-resolved methylation. Color gives the Pearson correlation coefficient, adjusted for AD status. No association reached significance.

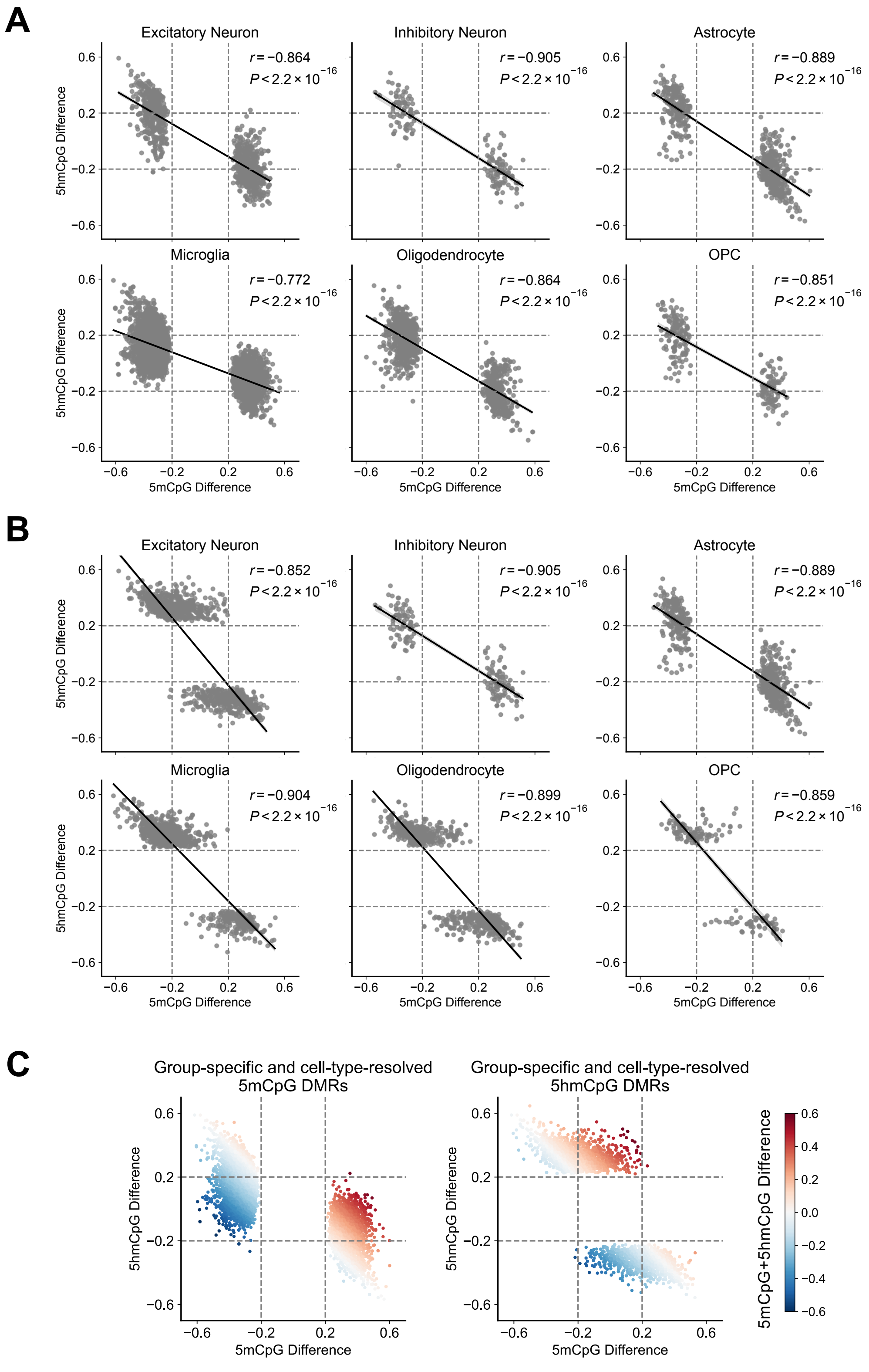


**Figure S20. Reciprocal 5mCpG and 5hmCpG changes at group-specific and cell-type-resolved DMRs.**

**(A–B)** Relationship between the 5mCpG difference (x-axis) and the 5hmCpG difference (y-axis) at group-specific and cell-type-resolved **(A)** 5mCpG DMRs and **(B)** 5hmCpG DMRs, shown separately for each brain cell type. Each point represents one DMR, with the solid line giving the linear fit and the shaded area its 95% confidence interval. Pearson's *r* and *P* values are shown in each plot, and dashed lines mark the ±0.2 difference threshold used for DMR identification. **(C)** The same comparison with all six cell types combined, for 5mCpG DMRs (left) and 5hmCpG DMRs (right), colored by the summed 5mCpG + 5hmCpG difference. Points nearer the anti-diagonal, where the two changes are closest to equal and opposite, show the strongest cancellation in the combined signal.

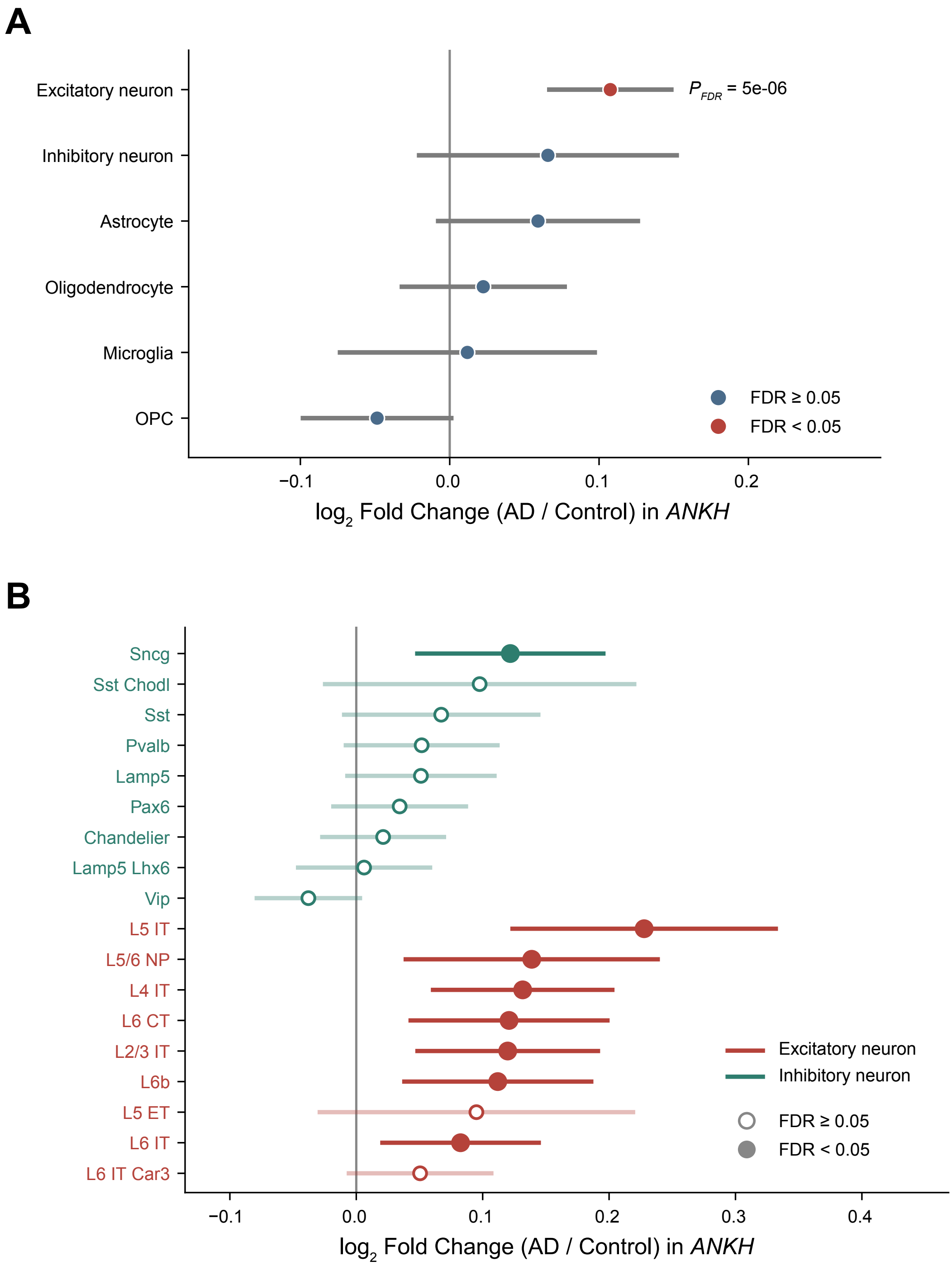


**Figure S21. *ANKH* expression is increased in excitatory neurons in Alzheimer's disease.**
**(A)** Differential expression of *ANKH* between AD and control donors across the six brain cell classes, from donor-level pseudobulk of public single-nucleus and single-cell RNA-seq data[7–14] from human cortex. Points show the $\log_2$ fold change (AD / control) estimated by ordinary least squares with study, region and sex as covariates and donor-clustered robust standard errors; horizontal lines show 95% confidence intervals. Red, FDR < 0.05; blue, FDR ≥ 0.05 (Benjamini–Hochberg across cell classes). The FDR-adjusted *P* value is shown for excitatory neurons, the only class reaching significance. **(B)** The same analysis at neuronal subclass resolution, for excitatory (red) and inhibitory (green) subclasses, ordered by effect size within each class. Filled circles, FDR < 0.05; open circles, FDR ≥ 0.05 (Benjamini–Hochberg across subclasses).

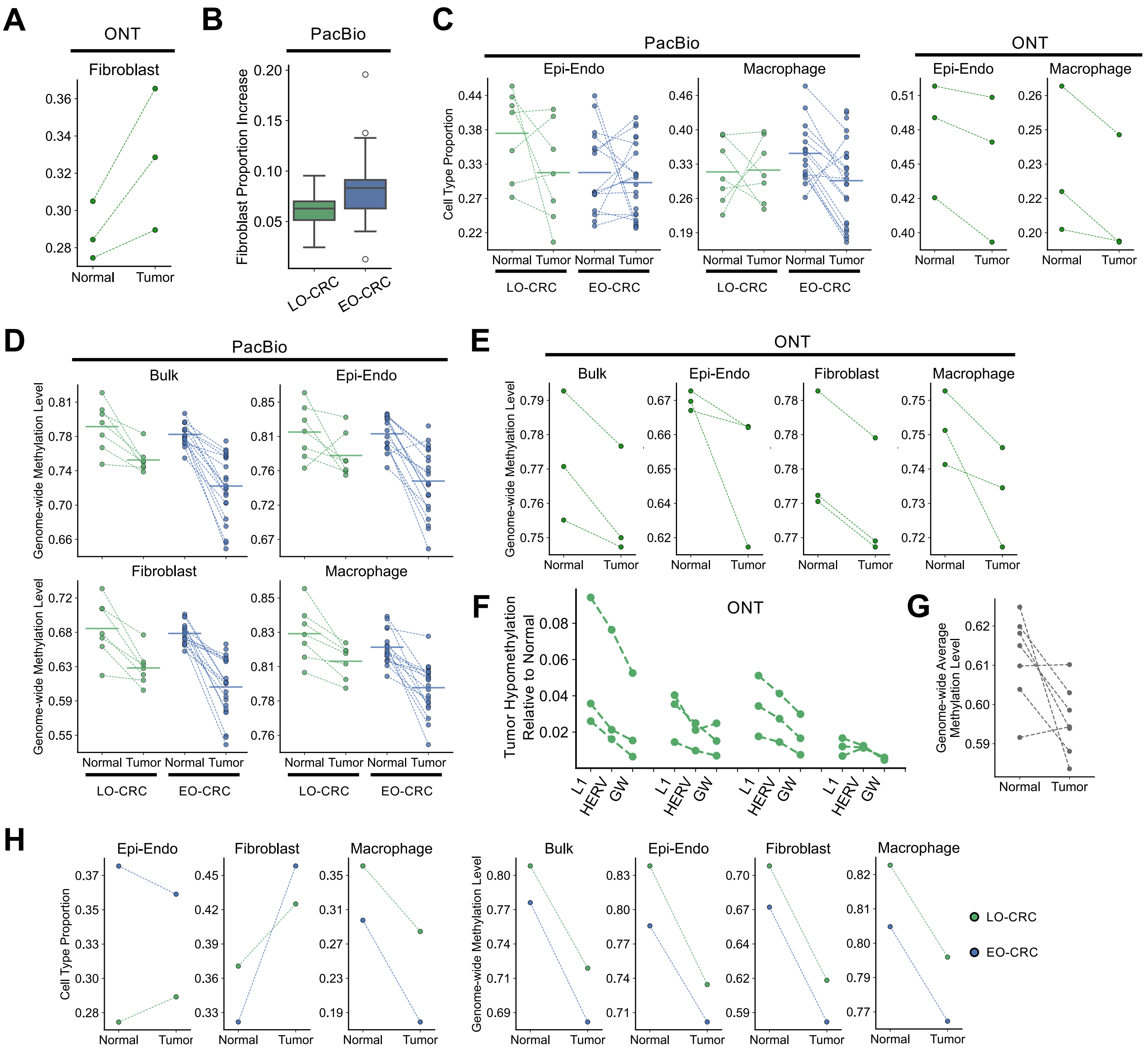


**Figure S22. Cell-type composition and methylation changes in paired CRC tumor and adjacent normal tissue.**

**(A)** COMET-estimated proportions of fibroblast in 3 LO-CRC pairs sequenced on ONT. **(B)** Fibroblast proportion increase (tumor minus matched adjacent normal) in LO-CRC (n = 7) and EO-CRC (n = 17) pairs sequenced on PacBio. The corresponding fibroblast result is also shown in **Fig. 6B**. **(C)** COMET-estimated proportions of Epi-Endo and macrophage in 7 LO-CRC and 17 EO-CRC pairs sequenced on PacBio (left) and 3 LO-CRC pairs sequenced on ONT (right). **(D–E)** COMET-estimated genome-wide methylation levels in bulk tissue and each cell type for **(D)** the 7 LO-CRC and 17 EO-CRC pairs sequenced on PacBio and **(E)** the 3 LO-CRC pairs sequenced on ONT. **(F)** Loss of methylation in tumor relative to matched adjacent normal (y-axis) in the ONT-sequenced pairs, quantified genome-wide (GW) and at L1 and HERV elements (x-axis) for bulk tissue and each cell type. **(G)** Genome-wide average methylation measured by methylation array in the 7 PacBio-sequenced LO-CRC pairs. **(H)** COMET-estimated cell-type proportions (left) and genome-wide methylation levels in bulk tissue and each cell type (right) for the two deep-sequenced pairs, one LO-CRC and one EO-CRC. Throughout, each point represents one sample and dashed lines connect the tumor and adjacent normal samples of the same pair. In **(A–F, H)**, green indicates LO-CRC and blue EO-CRC, with horizontal lines in **(C)** and **(D)** giving group means.

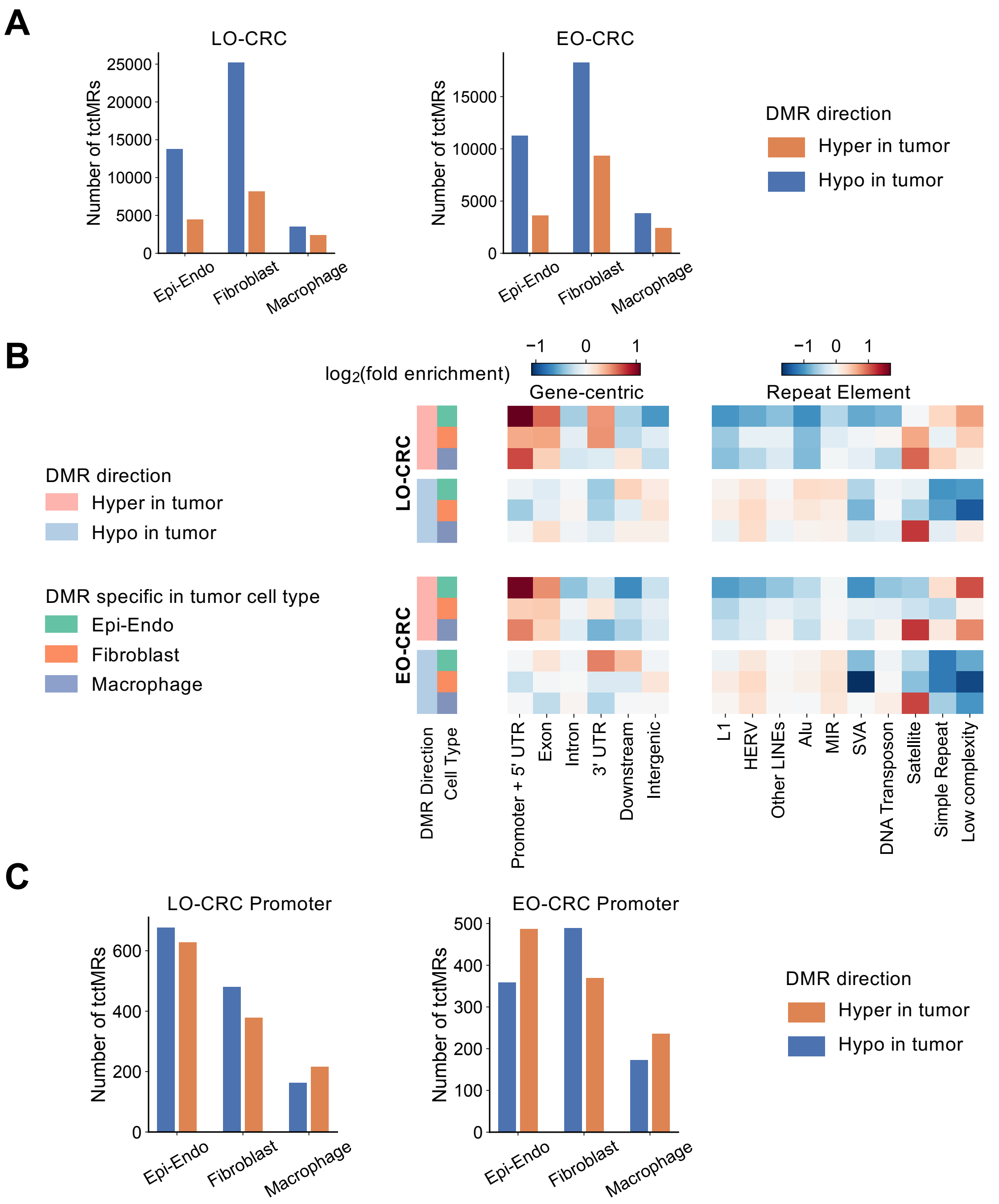


**Figure S23. Distribution and genomic context of tctDMRs in the deep-sequenced CRC pairs.**

**(A)** Number of tctDMRs identified in each of the three colon cell types, shown separately for the LO-CRC (left) and EO-CRC (right) pair. Orange and blue bars give tctDMRs hypermethylated and hypomethylated in tumor relative to matched adjacent normal. **(B)** Fold enrichment of tctDMRs at gene-centric features (left) and repeat element classes (right), shown for the LO-CRC (top) and EO-CRC (bottom) pair. Each row is a tctDMR set defined by the direction of the methylation difference and the cell type to which it is specific, as indicated by the annotation bars on the left, and each column is an annotation category. Colour gives $\log_2$(fold enrichment). **(C)** Number of tctDMRs mapping to gene promoters in each cell type, shown separately for the LO-CRC (left) and EO-CRC (right) pair, with bars colored by the direction of the methylation difference as in **(A)**.

A

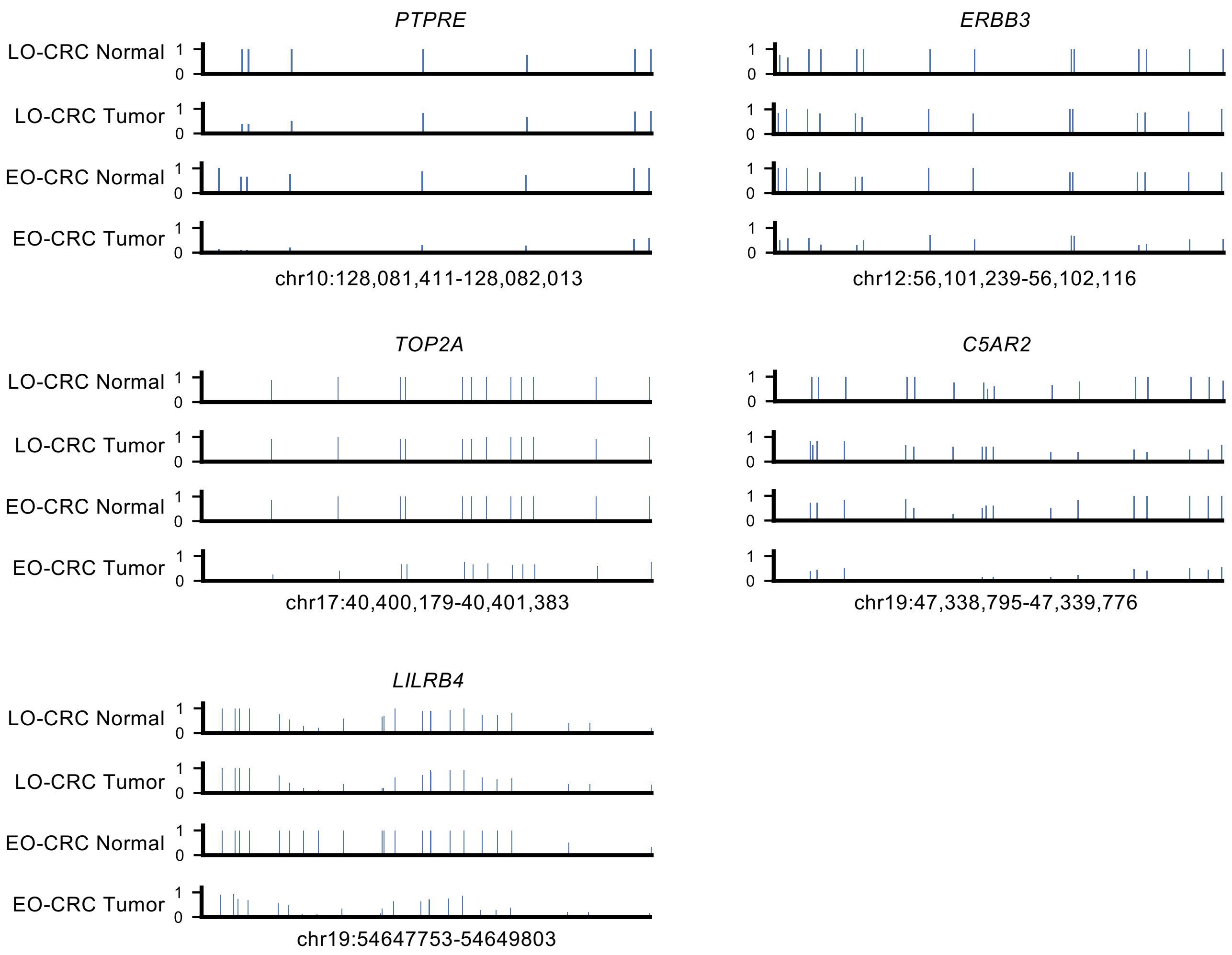


B

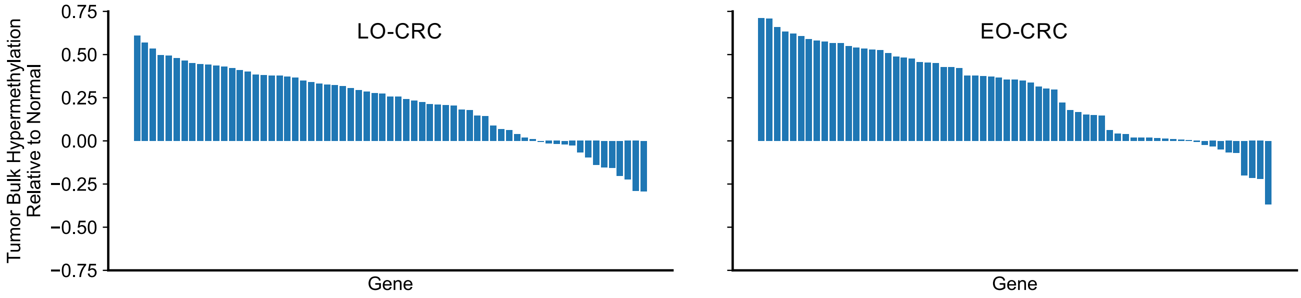


**Figure S24. Epi-Endo-resolved methylation at genes marking an EO-CRC-enriched epithelial state, and bulk methylation at CRC screening genes.**

**(A)** Epi-Endo-resolved methylation across *PTPRE*, *ERBB3*, *TOP2A*, *C5AR2* and *LILRB4*[15], the five genes showing reduced methylation in EO-CRC tumor relative to EO-CRC adjacent normal. For each gene, four tracks are shown, from top to bottom: LO-CRC normal, LO-CRC tumor, EO-CRC normal and EO-CRC tumor. Each vertical bar represents one CpG, with height giving its methylation level from 0 to 1, and the displayed interval is indicated below each panel. These genes are highly expressed in an epithelial cell state enriched in EO-CRC relative to LO-CRC in scRNA-seq data. **(B)** Bulk methylation difference between tumor and matched adjacent normal at the promoters of the 65 genes used in CRC screening panels, shown for the LO-CRC pair (left) and the EO-CRC pair (right). Each bar represents one gene, ordered by the size of the difference, with positive values indicating tumor hypermethylation. Per-gene methylation levels are given in **Supplementary Table 14**.

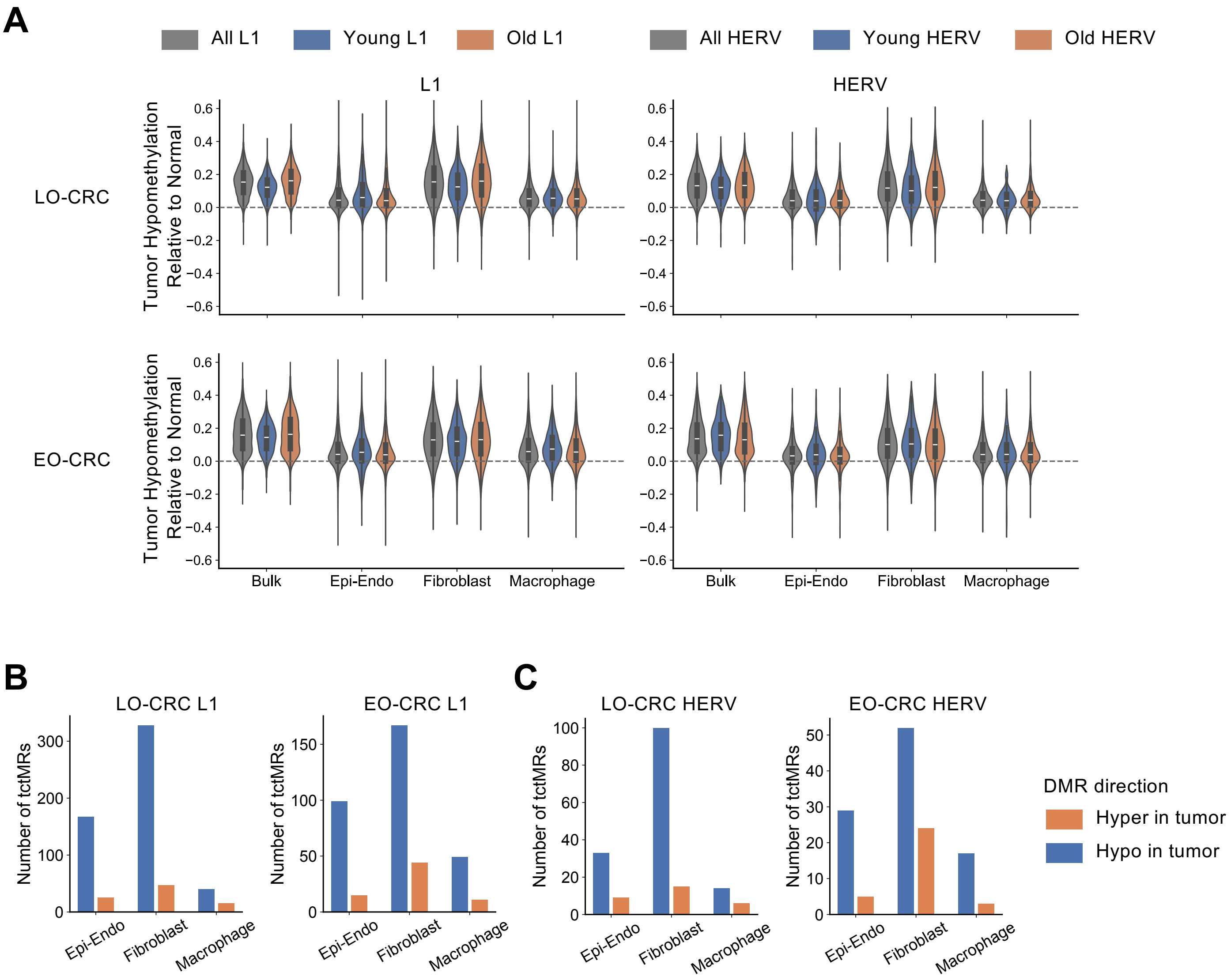


**Figure S25. CRC methylation changes at L1 and HERV elements in the deep-sequenced pairs.**

**(A)** Bulk and cell-type-resolved methylation loss in tumor relative to matched adjacent normal at L1 (left) and HERV (right) elements, shown for the LO-CRC (top) and EO-CRC (bottom) pair, with positive values indicating tumor hypomethylation. Elements are grouped by evolutionary age, with grey giving all elements, blue the young subfamilies (L1HS and L1PA2 for L1, HERVK for HERV) and orange the remaining older subfamilies. Violins show the distribution across elements, with the white line giving the median, and the dashed line marks no change. **(B–C)** Number of tctDMRs falling within **(B)** L1 and **(C)** HERV elements in each of the three colon cell types, shown for the LO-CRC and EO-CRC pair. Orange and blue bars give tctDMRs hypermethylated and hypomethylated in tumor relative to matched adjacent normal.
