## Supplementary Tables for "Cell-type-resolved DNA methylation analysis using bulk long-read sequencing across human tissues": Supplementary Table Legend.docx

**Supplementary Table 1.** **Sequencing and quality control metrics for all samples profiled in this study.** Per-sample sequencing metrics for the 297 samples sequenced on the PacBio Revio platform (140 blood, 53 colon, 104 brain samples) and the 6 colon samples sequenced on ONT P2, giving tissue, sequencing platform, base modification calling model, depth, median read length and total yield. The 18 deep-sequenced samples are indicated.

**Supplementary Table 2.** **Samples of defined cell-type composition used for method validation.** Sequencing metrics for the samples used to evaluate COMET against known cell-type content, giving sample type, tissue validated, sequencing platform, base modification calling model, depth, median read length and total yield.

**Supplementary Table 3.** **COMET-estimated cell-type proportions and cell-type-resolved methylation levels for all blood samples.** Per-sample proportion of reads assigned to each of the six blood cell types (B Memory, B Naive, myeloid, NK, T Ag-exp, T Naive), and the genome-wide 5mCpG methylation level at bulk level and computed within each cell type.

**Supplementary Table 4.** **COMET-estimated cell-type proportions and cell-type-resolved methylation levels for all brain samples.** Per-sample proportion of reads assigned to each of the six brain cell types (excitatory neurons, inhibitory neurons, astrocytes, microglia, oligodendrocytes and OPC), and the genome-wide 5mCpG+5hmCpG modification level at bulk level and computed within each cell type.

**Supplementary Table 5.** **COMET-estimated cell-type proportions and cell-type-resolved methylation levels for all colon samples.** Per-sample proportion of reads assigned to each of the three colon cell types (Epi-Endo, fibroblast, macrophage), and the genome-wide 5mCpG methylation level at bulk level and computed within each cell type.

**Supplementary Table 6.** **Characteristics of the blood cohort.** Donor-level information for the 140 whole blood samples, including source cohort, age at collection, sex, APOE genotype, APOE ε4 carrier status, NDD-related clinical record, plasma p-tau217 and neurofilament light values where available, resilience status and availability of matched methylation array data.

**Supplementary Table 7.** **Decomposition of age-related bulk blood methylation change into compositional and intrinsic contributions.** For each comparison between consecutive age bins ([0,20) versus [20,40), [20,40) versus [40,60), [40,60) versus ≥60): the per-cell-type intrinsic and compositional contributions, the covariance and measurement residual terms, each reported as its contribution to the bulk methylation change and as its percentage share of the total bulk methylation decrease.

**Supplementary Table 8.** **Group-specific, cell-type-resolved differentially methylated regions in blood.** DMRs called between the APOE44 resilience candidates, non-resilient APOE44 carriers and APOE4 non-carrier controls from the 6 deep-sequenced blood samples, reported with genomic coordinates, number of CpGs, group specificity, methylation direction, cell type and per-sample methylation level.

**Supplementary Table 9.** **Regulatory annotation of group-specific myeloid differentially methylated regions in blood.** Annotation of the myeloid-resolved, group-specific DMRs against ROADMAP histone modification states, derived promoter and enhancer states, and against promoter-capture Hi-C interactions, giving the annotated regulatory class, the number of significant interactions, the myeloid cell types supporting them and the linked genes.

**Supplementary Table 10.** **Characteristics of the brain cohort.** Donor-level information for the 104 postmortem brain samples, giving cohort, brain region, age at death, sex, CERAD score and Braak stage for all samples, and additionally clinical cognitive status, combined ADNC score and final diagnostic classification for the 28 NPBB samples.

**Supplementary Table 11.** **Group-specific, cell-type-resolved 5mCpG and 5hmCpG differentially methylated regions in brain, with gene annotation.** DMRs called separately for 5mCpG and 5hmCpG from the 6 deep-sequenced NPBB samples, reported with genomic coordinates, number of CpGs, group specificity, modification type, cell type, direction and per-sample modification level, together with the annotated gene name, GENCODE gene ID and annotated gene region.

**Supplementary Table 12.** **Characteristics of the colorectal cancer cohort.** Specimen-level information for the 59 early-onset and late-onset colorectal cancer specimens, including individual ID, age at diagnosis, sex, specimen type, sequencing platform and availability of matched methylation array data. Tumor and adjacent normal specimens from the same patient share an individual ID.

**Supplementary Table 13.** **Tumor- and cell-type-specific differentially methylated regions in colorectal cancer, with annotation.** tctDMRs identified from the two deep-sequenced matched tumor–normal pairs, reported with genomic coordinates, number of CpGs, CRC type, cell type of origin, direction of change, the contributing normal and tumor sample IDs, and the methylation level in each cell type in both specimens. Annotation is given against ROADMAP 15-state ChromHMM chromatin states of colonic mucosa, CpG island context, gene-centric features and candidate promoters based on GENCODE v45, and RepeatMasker repeat elements including full-length L1 and HERV classification.

**Supplementary Table 14.** **Cell-type-resolved methylation at colorectal cancer screening panel genes.** The 65 genes of the QIAseq Targeted Methyl Human Colorectal Cancer Panel, with the candidate promoter region evaluated for each gene (1,500 bp upstream to 500 bp downstream of the TSS). For each gene, the table gives the gene name, the candidate promoter coordinates and the number of CpGs in the region, followed by methylation at bulk level and resolved to each of the three colon cell types in the four deep-sequenced samples. Each sample and cell type is reported as the methylated call count, the unmethylated call count and the resulting methylation level. Cell-type-resolved methylation levels are shown as “(depth <3×)” where that cell type was covered at less than 3× in that sample.

**Supplementary Table 15.** **Antibodies and staining reagents.** Each antibody is listed with its target, clone, source and catalogue number. The upper table gives the antibodies used for fluorescence-activated sorting of CD49f^+^ long-term hematopoietic stem cells prior to iPSC reprogramming. The lower table gives the mass cytometry immune profiling panel, with the metal isotope conjugated to each antibody and whether it belongs to the Maxpar Direct core panel or was spiked in as an add-on.
